# Genetic association data are broadly consistent with stabilizing selection shaping human common diseases and traits

**DOI:** 10.1101/2024.06.19.599789

**Authors:** E. Koch, N. J. Connally, N. Baya, M. P. Reeve, M. J. Daly, B. M. Neale, E. S. Lander, A. Bloemendal, S. Sunyaev

## Abstract

Results from genome-wide association studies (GWAS) enable inferences about the balance of evolutionary forces maintaining genetic variation underlying common diseases and other genetically complex traits. Natural selection is a major force shaping variation, and understanding it is necessary to explain the genetic architecture and prevalence of heritable diseases. Population genetics models that make different assumptions about selection at the phenotypic level also make different predictions about genetic architectures. We develop an inference framework to test existing population genetics models based on the joint distribution of allelic effect sizes and frequencies of trait-associated variants. Here, we analyze multiple datasets for anthropometric traits, metabolic traits, and binary diseases—both early-onset and post-reproductive. We test models of neutral evolution, long-term directional selection (selection against a trait or against disease risk) and stabilizing selection, which acts to preserve an intermediate trait value or disease risk. We evaluate pleiotropic models of stabilizing and purifying selection, where the fitness effects of each variant are influenced by a multitude of selected traits. We find that most traits show consistency with stabilizing selection and suggest that selection is pleiotropic.

## Introduction

Many human traits, including diseases, are known to be highly heritable, and affect the lifespan or reproduction of individuals. These qualities are the basis of natural selection, which acts to shape phenotypic distributions and reduce disease prevalence by removing maladaptive genotypes. With complex traits, a long-standing question in evolutionary biology is how substantial heritable variation is maintained in the presence of selection acting to reduce variation [1, 2]. In human biomedical genetics, the same question appears in the seeming contradiction between the high prevalence of complex diseases in the population and their observed reduction in fitness [3, 4, 5].

It has been proposed that some conditions, such as breast cancer, have persisted due to their post-reproductive onset. Classic theories of aging put forth the idea that selection declines with age and that late-acting harmful effects can persist when their effects on fitness earlier in life are weakly adverse, neutral, or beneficial [6, 7, 8, 9]. In the simplest version of this argument, a trait with no effect on the chance or ability to reproduce would, despite being harmful to individuals, have little or no effect on fitness. In the absence of other evolutionary forces, genetic variation underlying such a trait could evolve nearly neutrally and could occur at high frequency. This argument cannot explain polygenic conditions that obviously occur earlier in life, such as schizophrenia, which is estimated to reduce reproduction by up to 75% [10, 11, 12], but continues to have a population prevalence of 1% [13]. Even for late-onset traits, the premises of this explanation are challenged by model organism experiments and analysis of post-reproductive mortality and morbidity [14, 15, 16]. More generally, genetic variation underlying such phenotypes may also have earlier, less severe effects at reproductive ages. These early-acting effects, both on the focal trait itself and through other traits with shared genetic variants, may nevertheless be sufficient to induce effective natural selection.

More recent studies have addressed the question of trait-associated selection in the context of genome-wide association studies (GWAS). These studies focused on how negative selection shapes the relationship between variant frequency and the magnitude of trait effects. This formalizes the well-known intuition that GWAS should not identify common variants of large effect that explain a substantial fraction of the heritability of complex traits [17, 18]. Random-effects models based on this idea have produced a consistent conclusion across quantitative traits and complex diseases: on average, negative selection acts more strongly on variants with larger trait effects [19, 20, 21, 22, 23, 24, 25]. While clearly identifying the phenomenon, these studies did not explore the manner in which selection acts and whether it was associated with a particular trait direction. In order to explain the maintenance of trait variation and disease in a population, we must consider which among multiple models of selection could plausibly generate this pattern of negative selection. Selection is most readily modeled at the level of individual variants, where variant frequencies and inheritance define a clear genetic system. Direct inference of selection on traits requires measuring fitness components for phenotypes themselves, so here we ask a more restricted question about how different models of trait-level selection would appear among the genetic variants associated with each phenotype.

For schizophrenia—and other complex human diseases with measurable fitness costs—it is natural to propose a model in which selection acts to remove risk-increasing alleles, but is balanced by an incessant influx of new mutations (mutation-selection-drift balance) [26]. This directional selection against risk alleles could result from selection on liability, but could also arise in a model where disease liability reflects disruptions to biological processes that have been optimized over long periods of evolutionary time, such that most new mutations are selected against and increase risk. This directional selection assumption seems to be supported by the observation that variants causing monogenic familial versions of diabetes, breast cancer, and other diseases are rare in the population [27, 28]. Similar observations have been made for some quantitative traits. For example, large-effect coding mutations, on average, lead to a decrease in height and hand grip—possibly counterbalanced by directional selection [29, 30, 31]. However, it has not been shown that selection on common variants acts directionally, and there may be other explanations.

Under the above model, selection relevant for variation in the trait acts primarily on the trait itself. Alternatively, variation underlying the trait may impact many other traits that collectively determine fitness (pleiotropic selection). In the extreme, the importance of the trait value on fitness is negligible, and the trait variation is mostly determined by other (pleiotropic) consequences of genetic variation.

Even models with selection primarily acting directly on the trait have different flavors. Directional selection reduces maladaptive traits in a population, such as high risk for a (binary) disease, or harmfully high levels of a quantitative trait [14, 15, 32, 2]. Another model, stabilizing selection, works to maintain a trait at its current level in the population, while also reducing variance [33, 34, 35, 36, 32, 37, 1, 2]. The stabilizing model is commonly assumed for many quantitative traits, such as blood pressure, which can be dangerous at either extreme. The average value of the trait corresponds to the highest fitness. The fitness is reduced with the increasing deviation from the average. Stabilizing selection models were recently re-introduced to human complex trait genetics by Simons *et al*. (2018) [37], and have provided quantitative explanations for the frequencies and effect magnitudes of variants discovered by human GWAS [38], the relationship between quantitative human phenotypes and lifespan [39], and the enrichment of large-effect cis-regulatory variants at low frequencies [40].

It is therefore instructive to contrast models of stabilizing and directional selection from the perspective of individual alleles. Under the directional model, selection acts against the risk allele, and the direction of the phenotypic effect determines the direction of selection. Under the stabilizing model, the consequence of optimal fitness for an intermediate value, is that selection acts against the minor (less common in the population) allele, and the allele frequency determines the direction of selection [33, 34, 35].

Binary traits, such as disease status, cannot exist at intermediate levels within individuals by definition. On the other hand, the liability of a binary trait (the genetic contribution to its occurrence) can be intermediate and therefore subject to stabilizing forms of selection. In some cases, liability is known to be mediated through an observable trait, as with blood pressure and hypertension [41]. For other traits, liability may correspond to an underlying trait such as glucose level for diabetes [42] and FEV1/FVC ratio for asthma [43], or it may be an abstracted measurement of genetic risk. If stabilizing selection acts on liability for many diseases, it could help explain their counter-intuitive prevalence. For example, schizophrenia’s prevalence could be high because stabilizing selection has maintained intermediate levels of schizophrenia liability in populations by preventing evolution to both high and low extremes. However, this model has not been deeply explored for diseases, as it is not intuitive that individuals with an unusually low risk of disease would have a fitness disadvantage.

At the same time, widespread pleiotropic effects of genetic variants on multiple traits [44] must also be considered when explaining maintenance of trait or disease variation. Pleiotropic models assume that variants affect multiple traits, sometimes to the extent that selection on genetic variation underlying a given trait is only minimally concerned with the trait itself but rather with myriad unrelated (pleiotropic) consequences. The literature offers two categories of such models. In one category of models, variants affect many traits. Contributing variants are affected by selection on the focal trait, but also on other traits they influence. We call this category of models “multidimensional”. An early version was suggested by Zhang and Hill [45], and Simons *et al*. [37] developed an inference-oriented framework to describe the relationship between selection and effect sizes in GWAS. The other category of pleiotropic models assumes that phenotypically relevant variants are mostly deleterious due to unknowable effects on numerous unmeasured traits. The focal trait has no meaningful or measurable contribution to selection, but its genetic architecture is determined by selection on its underlying variants. Such models proposed by Barton [46], Kondrashov and Turelli [47] and Eyre-Walker [48] may be called “pure pleiotropy” or “apparent stabilizing selection.” Despite their different perspectives, these pleiotropic models share some properties with simple models of stabilizing selection — selection maintains the focal trait at intermediate level and reduces its variance. From the viewpoint of individual alleles, these models differ. Under the multidimensional model, selection acts against minor alleles. Under the apparent selection model, selection acts against derived alleles (those arising due to new mutations).

The discovery of associated variants from genome-wide association studies (GWAS) offers a window into the evolutionary dynamics and genetic architecture of many traits [1, 2, 49] and an opportunity to test which of the above scenarios is most consistent with human genetic variation. We take a set of traits with sufficient numbers of associated variants (minimum 40 after filtering), and examine the relationship between the effect size and frequency of the trait-increasing (or risk-increasing) allele. Visualizing this relationship in a scatter plot reveals that most traits share a similar distribution, with a U-shaped curve that resembles a “smile” (Figure 1). In measuring the relationship between signed effect sizes and frequency, these plots are similar to “trumpet” plots [29, 50], but are organized to facilitate observations about evolution and selection.

**Figure 1:**
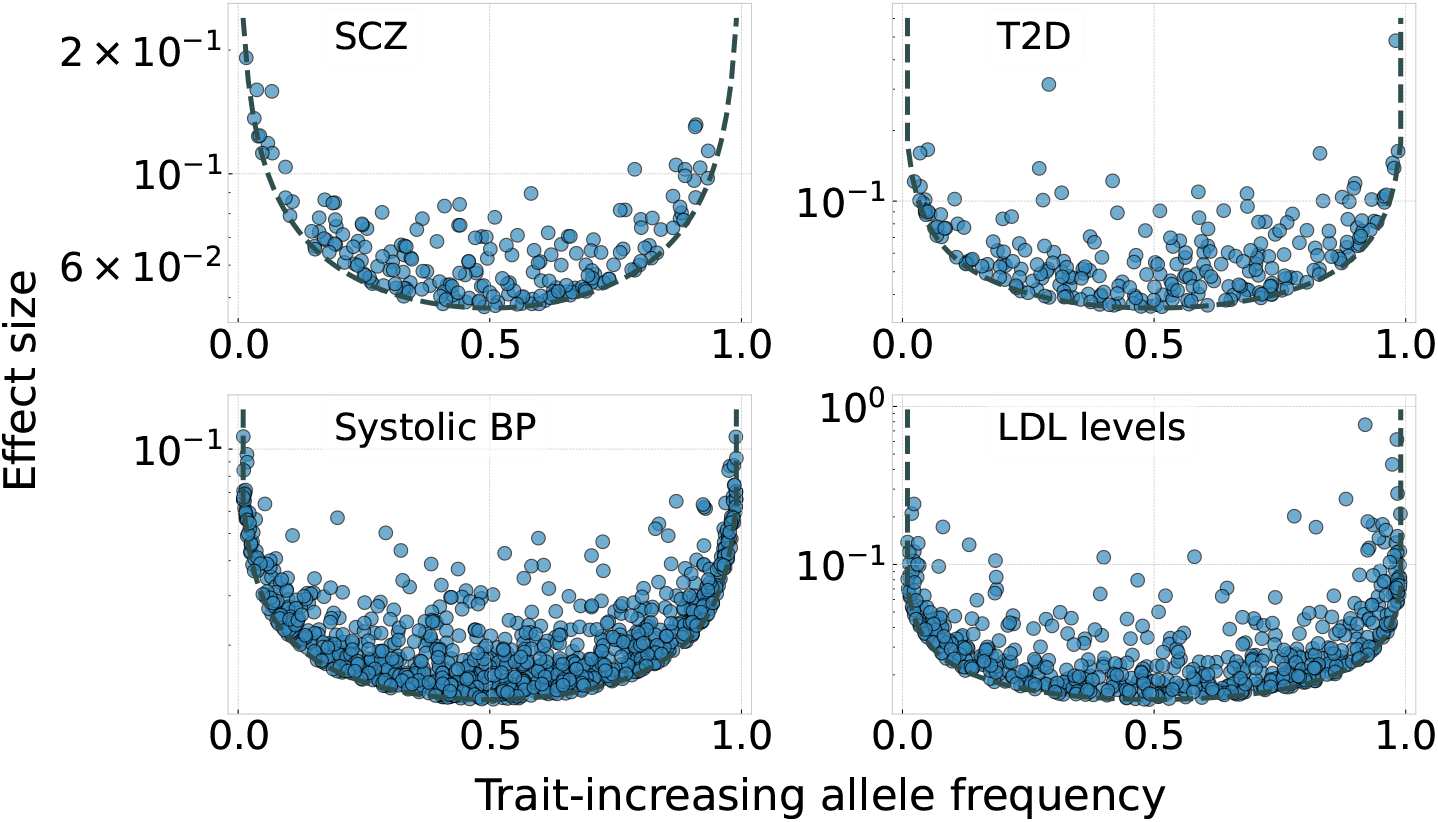
A U-shaped distribution for the effect sizes and allele frequencies after polarizing by direction of effect. For four example traits we plot the GWAS-estimated effect sizes of trait or disease risk-increasing alleles against their frequencies. For disease traits, the effect size is calculated as the odds-ratio minus one. We include variants that are both genome-wide significant and have an estimated variance contribution exceeding a threshold variance associated with 5 × 10^−8^.

The “smiles” curves suggest some initial inferences about selection on these traits. The lower edge of the curve represents the limits of statistical power, and is not reflective of selection. The upper edge of the curve is the result of an absence of large-effect variants at intermediate frequencies. This is clear evidence of selection: under neutrality, large-effect variants could drift freely to any frequency, whereas selection prevents them from doing so. If selection were directional, acting either to decrease or increase the trait, we would expect to see large-effect variants at one side of the distribution (where the effects of selection are balanced against the influx of new mutations), but not at both sides (Figure 2b). The symmetry of the smile is consistent with the symmetry of stabilizing selection (direct or apparent), which acts against both trait-increasing and trait-decreasing variants. Additionally, the location of the upper edge and spread of observed variants may be influenced by pleiotropy.

**Figure 2:**
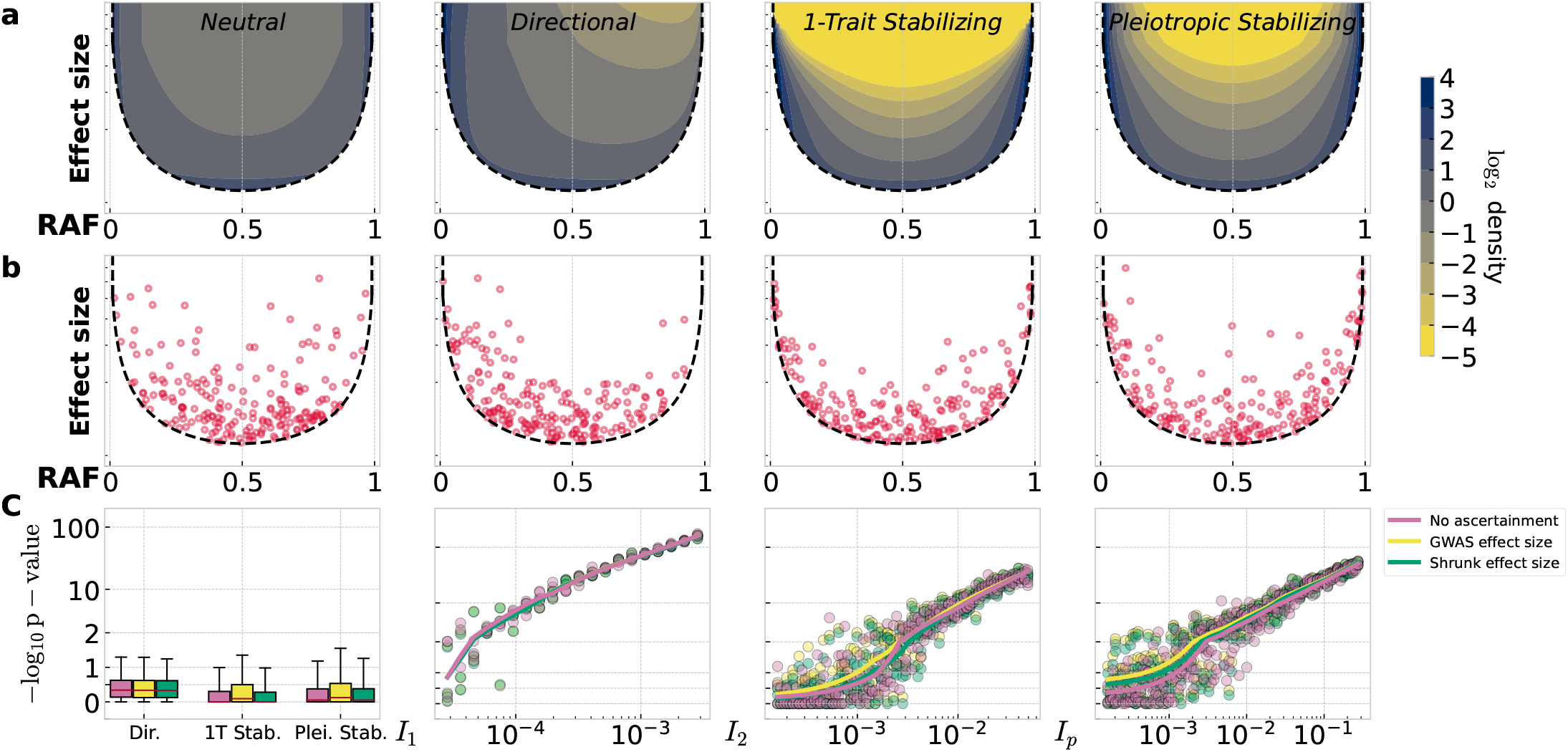
Selection leaves an identifiable pattern in the distribution of GWAS hits. a) The probability densities for the joint distributions of variant frequency and effect size show the expected shapes of smiles under different models. Neutrality produces a more filled-in smile. Directional selection produces a more asymmetric smile. Single-trait stabilizing selection produces a symmetric smile that is not filled-in. Pleiotropic stabilizing selection produces a thicker smile than stabilizing selection on one trait. b) Simulations show examples of the distributions of variant effect sizes and risk-increasing (or trait-increasing) allele frequencies (RAF) expected under different models. Ascertainment is modeled using a variance-contribution threshold. An absence of selection results in a large number of large-effect common variants. Directional selection is modeled assuming mutational equilibrium. New mutations are highly biased by their effect direction on the trait, and most are deleterious. Stabilizing selection is considered at single-trait (1D) and pleiotropic (hD) extremes. Both result in a U-shape, with symmetry in RAF space and a lack of high-frequency, large-effect variants. c) Simple models using variant frequencies conditional on effect size are able to detect selection. Evidence for selection is quantified using p-values relative to the neutral model where selection intensities are zero, values of 1 and above can be considered strong evidence. Neutral simulations rarely produce false selection signals. Selection is reliably detected in all models once the intensity exceeds a threshold. *I*_1_, *I*_2_, and *I*_*p*_ represent the intensity of selection in the directional, single-trait stabilizing, and pleiotropic stabilizing models respectively. Simulation results presented here used the distribution of effect sizes estimated for common IBD variants and a model of European population history from Tennessen *et al*. (2012) [57]. Each simulation was fixed at 200 ascertained loci.

## Methods and results

### Models and Inference

We analyzed the consistency of observed GWAS data with models of selection on complex traits, including a neutral model used to represent the case where selection is either undetectable or does not manifest within GWAS at the current resolution. These models can be characterized by the change in frequency of a variant under selection as a function of the variant’s effect on the focal trait [35, 51, 1, 52] (Table 1). We also analyzed models in which a neutral focal trait is influenced by selection from the pleiotropy of its contributing variants. Such “apparent selection” models can make very similar predictions to trait-selection models, but we attempted to distinguish them through two features: the frequency-dependent strength of selection and the scaling of selection with trait effects (Supplement I). For this reason, we leave purely pleiotropic models to later in the text and start by analyzing the consistency of observed GWAS data with models of neutrality and four different forms of trait selection.

**Table 1:**
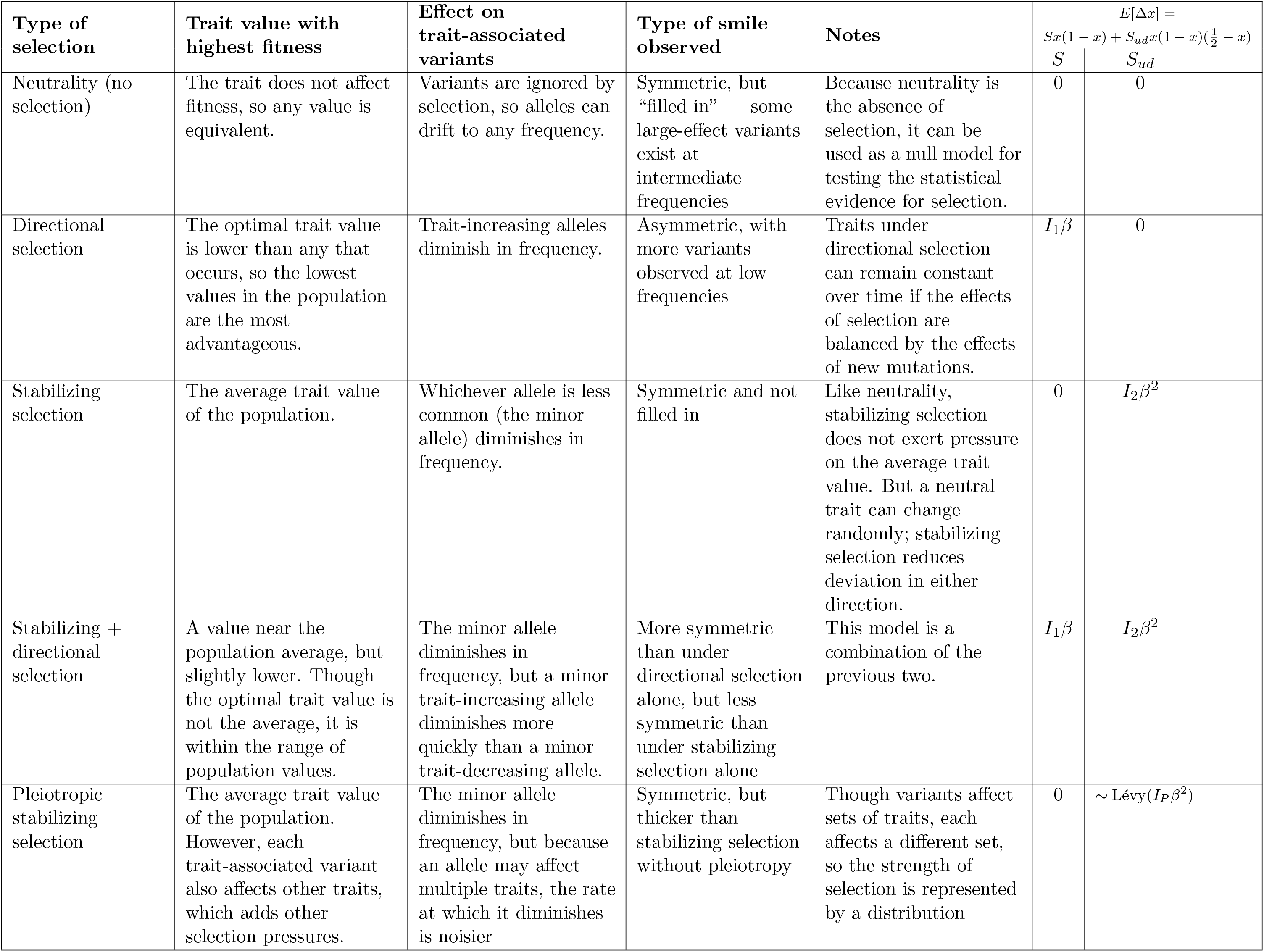
A summary of different selection models, their effects on smiles plots, and their parameters. Each selection model assumes a different relationship between traits and fitness, which leads to different effects on trait-associated variants, and qualitatively different smiles plots. The parameters *S, S*_*ud*_, *S*_*P*_ are the coefficients for directional selection, underdominant selection (which occurs when a trait is under stabilizing selection), and pleiotropic selection. They are used to model the expected change in allele frequency, 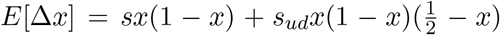. Directional examples were written as though selection is acting against a maladaptive trait, but would be reversed if selection is acting to promote a beneficial trait. The “pure pleiotropic” or apparent stabilizing selection model applies additive selection (*S*) without underdominant selection (*S*_*ud*_). The effects of pure pleiotropic selection depend on whether an allele is ancestral or derived, not whether it is trait-increasing or trait-decreasing. It is considered separately from other models (Figure 4a), so it is omitted from this table.

For the models of neutrality and trait selection, the dynamics of frequency changes can be used to predict the distribution of ascertained trait-associated variants for the model (Supplement B.4) under the assumption of mutation-selection-drift equilibrium (Figure 2a,b). The four models of trait selection are directional selection [51, 2, 26], stabilizing selection [35, 51, 1, 2], a combination of directional and stabilizing selection [53], and pleiotropic stabilizing selection (the “multidimensional” model [45, 37, 38]). By varying the selection parameters shown in Table 1, we can find the best possible fit for a given model for the observed data. The strength of this fit is compared against the neutral model using a likelihood ratio test, and likelihood differences, adjusting for which models use one or two parameters are employed to judge the strength of evidence between different models of selection.

### Application to the UK Biobank and FinnGen datasets

We initially fit each model to the GWAS summary statistics for 27 traits: 13 quantitative traits and 14 binary disease traits (Section A.1, Figure 3a,b). These included GWAS from the UK Biobank, FinnGen, and GWAS of individual diseases. For six diseases, we could not reject the neutral model at the adjusted significance threshold (Bonferroni-adjusted, 0.05/27). This was possibly due to power limitations for asthma, gallstones, glaucoma, and uterine fibroids, which all had fewer than 100 GWAS hits retained. IBD (199) and hypothyroidism (277) both had substantially more ascertained variants, making a power explanation less likely. All other traits showed greater consistency with stabilizing selection than with neutral or directional models. The pleiotropic model of stabilizing selection proposed by Simons *et al*. [37] was consistently a better fit than the single-trait stabilizing selection model, though the difference varied among traits (Figure S12).

**Figure 3:**
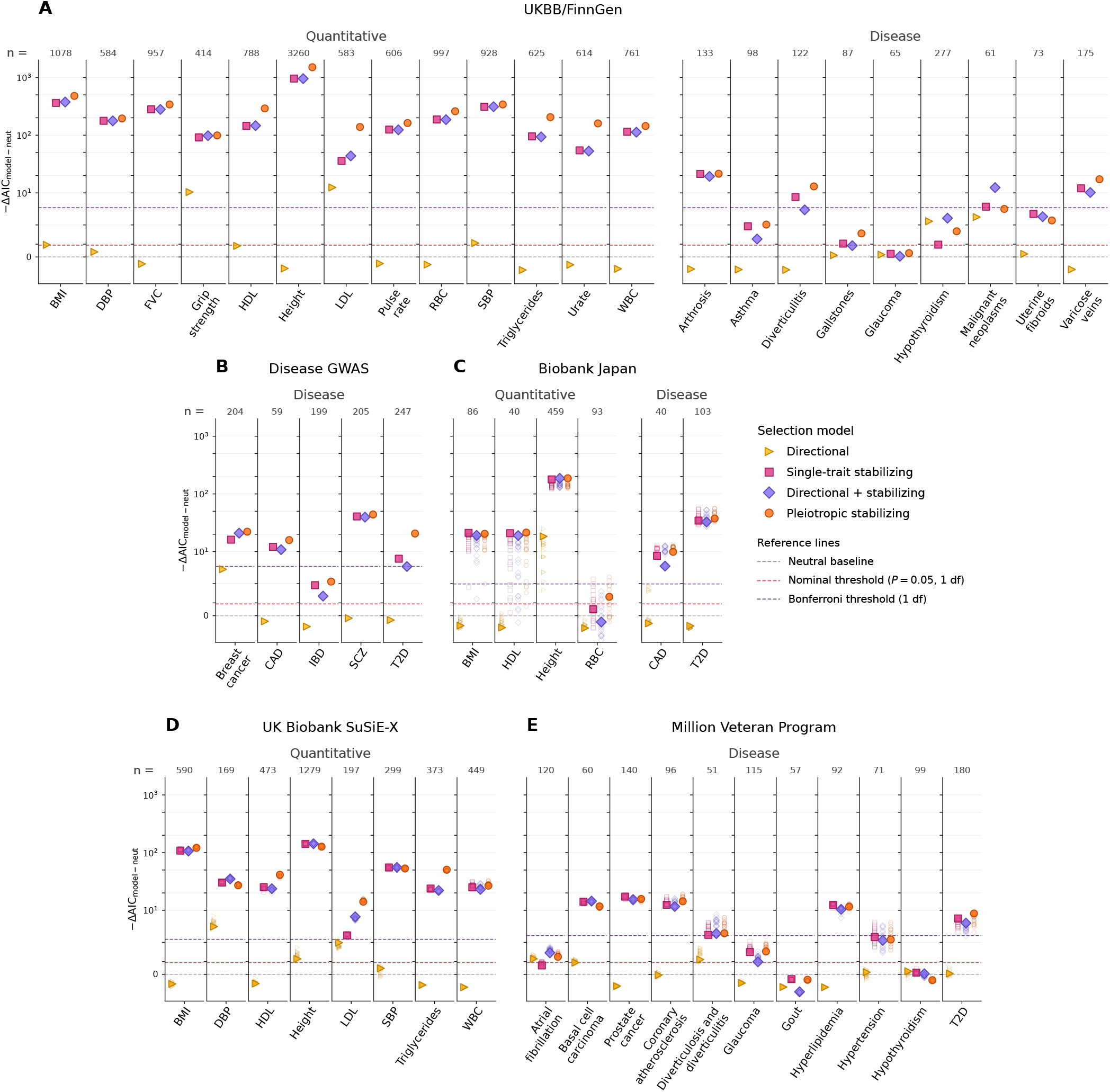
Model comparison favors stabilizing selection across categories of traits. We fit models of selection to each trait, estimating the intensity of selection under each model, and summarized results using AIC-scale likelihood differences relative to a neutral model. Reference lines show the corresponding one-parameter likelihood ratio test thresholds for *P* = 0.05 and Bonferroni-adjusted *P* = 0.05*/n*. To account for ascertainment bias in effect sizes we shrunk effect sizes by fitting the prior distribution of effect sizes using ASH [55] and computing posterior means from estimated GWAS effect sizes and standard errors. We use a variance-contribution threshold corresponding approximately to the p-value threshold 5 × 10^−8^ and also filter explicitly on p-value. In addition to the directional and stabilizing models shown in Figure 2, we also fit a model that combined directional and single-trait stabilizing selection. Larger filled points show the primary fit displayed for each trait, while faint open points show fits from repeated samples of one variant per locus or credible set where sampling was performed. a) The UKBB/FinnGen quantitative and disease traits from the original analysis. b) The five individual disease GWAS from the original analysis. c) Replication results for BBJ traits using the high-clumped version of the data. d) Replication results for UK Biobank quantitative traits using SuSiE-X fine-mapping, with one variant sampled from each credible set. e) Replication results for MVP disease traits using European-ancestry fine-mapping, again with one variant sampled from each credible set.

### Addressing potential confounders

The nature of GWAS data introduces technical complications to this inference. As a conservative strategy, we ascertain GWAS data to only include variants with genome-wide significance (*p <* 5 × 10^−8^) and exceeding a threshold variance contribution (Supplements B.2 & B.3). We explicitly model this ascertainment (Supplements B.4 and D). Next, true effect sizes and the identity of causative variants are unknown, and only their GWAS sampling estimates are available. The effect size estimates have a tendency to be larger than the true effects—a phenomenon referred to as winner’s curse [54]. To correct for winner’s curse, we do not use raw effect size estimates from GWAS, but fit selection models using shrunken estimates of trait effects [55] as well as a maximum likelihood adjustment for winner’s curse (Supplements D,D.2); both methods are supported by simulations (Figure 2b,c). Figure 3 reports shrunken effect size estimates. Our ability to identify which specific variants influence the trait is limited by linkage disequilibrium (LD)—the correlation of GWAS statistics for variants near one another. We analytically studied the impact of LD on our inference (Supplement B.1). We also tested robustness of our inference with respect to both winner’s curse and LD in full GWAS simulations (Supplement C.3).

Simulations under each model demonstrated the extent to which each produces a distribution of variant effects and frequencies similar to the characteristic “smile” observed in GWAS data, and whether models of trait selection can be distinguished in practice (Figure 2b,c). We found that, while the GWAS ascertainment boundary produced a superficial U-shape in many scenarios, statistical discrimination between models was less ambiguous. In simulated GWAS of neutral traits, inflated test statistics and false positives occurred if GWAS estimates of effect sizes were treated as true effects. Strategies to account for winner’s curse in GWAS largely removed this inflation in simulated data (Figure S40). When simulating selection, neutral models were rejected once the strength of selection became sufficiently high.

Last, the evolutionary demographic history of a population can alter the distribution of variants, even in the absence of selection [56, 57]. We incorporated realistic demographic history of the European population in the analysis [57] (Supplement E).

Figure 2c and Supplements A, D, E, G, and C present robustness analyses and simulations, and detail how these issues were handled in the inference. In order to further validate our findings, we performed several follow-up analyses to evaluate their sensitivity to the ancestry of the study population, uncertainty in causative variant identity, and other potential study-specific effects.

### Replication studies

While association studies in non-European ancestry populations have grown recently, in most instances sample sizes are not sufficient to produce the number of significant loci necessary for our analyses (Figure S3, Tables S2,S3). One major exception to this is the Biobank Japan [58]. We repeated the analyses of the original traits on GWAS that were also performed in Biobank Japan, using a model of Japanese demographic history [59] (Figure S32). We replicated signals of stabilizing selection for five quantitative traits and two disease traits (Figure 3c) despite association counts being far fewer for all traits. Signals of stabilizing selection remained robust when original effect-size estimates were replaced with matched BBJ estimates, providing an additional check against ascertainment bias in the original studies (Supplement G; Figures S17–S19). However, these analyses were hampered by the frequent absence of rarer original variants from the BBJ GWAS (Figure S18) [60].

Our original analysis represented each locus using the variant with the strongest association signal, after LD clumping. In reality there is uncertainty about which variant has a true effect on the trait, and this uncertainty propagates into effect sizes and variant frequencies. Fine-mapping methods quantify this uncertainty [61]. We used published fine-mapping of the same UK Biobank GWAS (SuSiEx, [62]) we originally analyzed to assess this uncertainty by sampling variants from associated loci proportional to their posterior inclusion probabilities and re-fitting models to each sample. This analysis replicated support for stabilizing selection models across all eight quantitative traits we re-analyzed (Figure 3d).

We similarly analyzed fine-mapping results of disease traits from a phenome-wide association study of the Department of Veterans Affairs Million Veteran Project (MVP) [63]. While MVP included ~30% of non-European ancestry, we had to restrict our analysis to the European ancestry sub-sample to obtain a sufficient number of phenome-wide significant (*p <* 4.6 × 10^−11^) hits to fit selection models. The MVP analysis therefore accounts for uncertainty in variant frequencies, as well as study-specific factors, but not any effects of continental ancestry. For the five traits represented in both the original and MVP datasets, we were able to replicate our findings: models of stabilizing selection fit better than neutral or directional models for coronary artery disease, diverticulitis, and type 2 diabetes, while we failed to find significant evidence for selection in glaucoma and hypothyroidism (Figure 3e). These similarities are not surprising given the shared ancestry of the study populations, but they support the idea that the initial results were not driven primarily by study-specific factors. We also found that GWAS data were consistent with stabilizing selection for two cancer traits not present in the original data: basal cell carcinoma and prostate cancer.

We concluded from these follow-up analyses that our inference of widespread consistency of stabilizing selection models with GWAS summary data was robust to major sources of potential bias. While we found no instances where the model of directional trait selection had higher likelihood than stabilizing models, the model combining directional and stabilizing selection provided the best fit for malignant neoplasms (UKBB/FinnGen) and DBP (SuSiE-X). This may indicate a directional component to selection in these traits, but could also reflect an unrelated mutational asymmetry or greater power to detect risk variants than protective ones [64]. Additionally, because the directional signal was only present in the fine-mapped version of DBP, it may be due to differences in which loci were retained between fine-mapping and standard GWAS clumping. In three other traits (LDL levels, grip strength, and height (BBJ)) the directional model out-performed neutrality, but each of these traits was still most consistent with pleiotropic stabilizing selection.

Pleiotropic stabilizing selection models had higher likelihoods than single-trait stabilizing models for 26 of 28 traits and diseases with significant selection signals in the UKBB/FinnGen, individual disease GWAS and BBJ analyses. This is consistent with previous studies showing that pleiotropic stabilizing models provide a good fit to the distribution of GWAS hits [37, 38]. In diagnostic comparisons, pleiotropic models achieved higher likelihoods by better fitting the joint occurrence of many low-frequency variants near the ascertainment boundary and a smaller number of common variants with appreciable effects (Figure S15). However, this preference was less consistent within the fine-mapping follow-up, where the pleiotropic model had a higher likelihood in 8 of 14 significant cases. We found that this was likely due to reported fine-mapping results preferentially leaving out loci near the GWAS ascertainment boundary (Figure S16, Supplement A.5). This likely reflects both the exclusion of loci with high uncertainty in causality among a large number of variants, where fine-mapping often fails, and the initial process by which variants were defined for inclusion in fine-mapping. Because near-boundary variants provide evidence for selection generally, and pleiotropic models are best suited to fit the resulting mixture of low-frequency near-boundary variants and common variants with appreciable effects (Figure S45), we treat consistency with stabilizing selection in fine-mapping datasets as a conservative test biased against pleiotropy.

### Models of pleiotropic purifying selection

Finally, we turned our attention to the question of whether the consistency with stabilizing selection we have noted is better explained by other models. In models of “pure pleiotropy” (“apparent” selection), the focal trait is unimportant and negative selection on trait-affecting variants is determined by a myriad of pleiotropic effects. Instead of enumerating another host of models, we focused on two additional predictions made by stabilizing models which would not obviously be true under alternatives. The first of these is that polygenic stabilizing selection appears equivalent to under-dominant selection at the per-variant level [33, 34, 35]. That is, the strength of selection becomes weaker as variants reach higher frequencies, due to compensating polygenic shifts from the rest of the genome, and even changes signs at 50% such that the minor allele is always disfavored. This allows variants to reach higher frequencies than under strict purifying selection (Figures S45). We tested whether models with additive selection (the assumption of the “purely pleiotropic” purifying selection) or underdominant models (characteristic of stabilizing selection) find stronger support in the data. When fitting the models, we kept the relationship between phenotypic effect sizes and selection coefficients the same as in stabilizing selection models to keep all features of the models except for effective underdominance intact. Using additive selection rather than the “underdominance” effect did not significantly improve the likelihood under the pleiotropic model for any trait, while the fit was significantly worse for 8 of 27 traits (Figure 4a).

**Figure 4:**
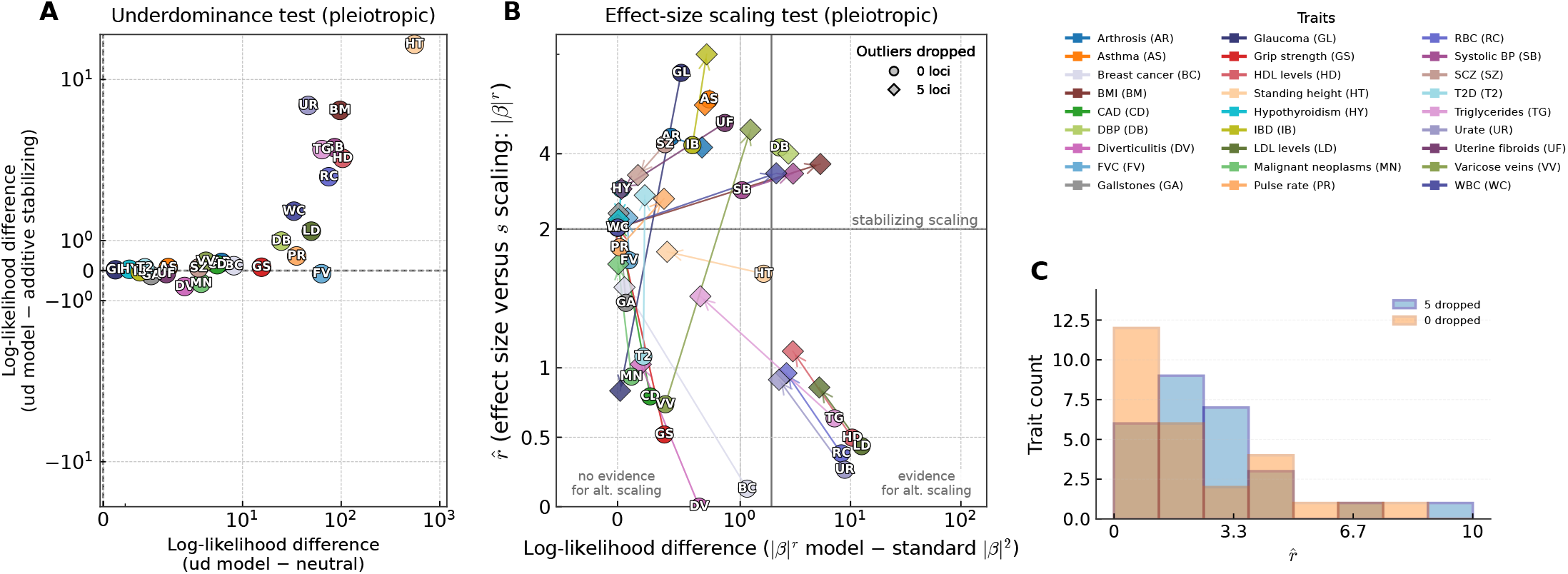
Most traits are consistent with underdominant selection and the effect size to selection scaling predicted by stabilizing selection. We evaluated two additional predictions made by stabilizing models of selection. a) For each original trait, we compare the likelihood of pleiotropic stabilizing selection models under underdominant and additive models of the site frequency spectrum. Positive values on the y-axis indicate that the underdominant form provides the better fit, while positive values on the x-axis indicate support for selection overall relative to neutrality. b) We fit models where the scaling coefficient *r* between effect size and selection (*s* = |*β*|^*r*^) was allowed to vary from the relationship under stabilizing selection (*r* = 2). Top outlier points were identified using leave-one-out likelihood deviation from the pleiotropic model. The estimated *r* and evidence for the model with free scaling versus the standard pleiotropic stabilizing model are compared for traits from the UKBB/FinnGen and disease GWAS sets. The vertical dashed line marks the nominal significance threshold, 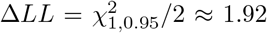, for the one-parameter extension from standard stabilizing scaling to free scaling. Sensitivity to potential outlier loci was assessed by re-fitting after dropping different numbers of outliers. c) The distribution of estimated scaling factors with and without outlier removal and refitting.

The second prediction made by models of stabilizing selection is that the selection coefficient arising from selection on the focal trait is proportional to *β*^2^ (Table 1) as long as the fitness function is sufficiently smooth near the optimum [37]. Alternative models of negative selection do not generally predict this specific scaling. We therefore fit models allowing the strength of selection to scale with |*β*|^*r*^ (Supplement I). Under pleiotropic stabilizing selection, 6 of 27 original traits showed significantly higher likelihood under the *r*≠ 2 model than under the standard *r* = 2 model (Figure 4b,c). This group included diastolic blood pressure, for which the estimated 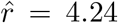 was about twice the stabilizing expectation, as well as several molecular quantitative traits, including lipid traits (LDL, HDL, triglycerides, RBC, and urate). Evidence for deviations from the *r* = 2 scaling attenuated as outlier loci were removed, though significant differences remained for five traits even after removing the top five outlier loci, and three additional traits showed evidence for *r >* 2. These results were not strongly dependent on whether the pleiotropic stabilizing or *r* = 0 model was used to define outliers (Figures S48). Likelihoods were more often improved in single-trait selection models when allowing *r*≠ 2 (Figure S47). However, these did not meaningfully improve overall model fit relative to pleiotropic stabilizing models, except for the outlier traits mentioned above. (Figure S49). In most cases, after considering these two additional predictions, pleiotropic stabilizing selection still appears most consistent with GWAS data. This conclusion is violated in a handful of traits that appear to have only weak evidence that selection differs between different forms of the relationship between phenotypic and fitness effects. Whether this is driven by true irrelevance of focal trait values, a large number of biological outlier loci, or other model violations remains unknown.

The comparison of natural selection models described above has not addressed the estimation of the intensity of selection on variants underlying phenotypic variation. Although quantitative estimation of selection strength is a more nuanced problem than model comparison, our statistical approach can be used to estimate selection coefficients. Uncertainty intervals for stabilizing fits to the original traits were broad, particularly for the pleiotropic intensity parameters (Figure S14). As shown in Figures S21–S29 the inferred selection strengths predicted for associated variants are of low-to-moderate magnitude. Supplementary Note B.7 discusses these estimates and their robustness.

## Discussion

Our results show that, across traits and datasets, selection is common and largely consistent with stabilizing models. In the original analysis of 27 traits, 21 showed evidence of selection based on the distribution of frequencies and effect sizes of associated variants. These 21 traits were best fit by stabilizing selection, usually pleiotropic and in one case with a minor directional component. Follow-up analyses in Biobank Japan and in UK Biobank and MVP fine-mapping largely supported the same conclusion: when selection was detectable, stabilizing models generally fit better than neutral or directional models. The evidence strongly favored pleiotropic stabilizing selection for many traits in the original and BBJ analyses, although the distinction between single-trait and pleiotropic stabilizing selection was less consistent in the fine-mapped datasets.

The intuitively appealing model of directional selection against disease is not strongly supported by the observed genetic architectures of traits. This is despite a power imbalance in the ability to detect risk vs. protective alleles, with risk variants being easier to observe for any binary trait with a low prevalence [64]. However, we cannot fully rule out a role for a directional component of selection. The few directional selection signals we do observe for diseases point to greater selection against risk alleles (*I*_1_ is consistently negative, Figures S13, S29). As noted in the introduction, rare variants with especially strong effects show a bias in their direction of effect for quantitative traits, which have much lower power difference between trait-increasing and trait-decreasing variants [29, 30, 31]. Higher-powered GWAS may uncover more imbalances for both quantitative and disease traits.

Higher-powered GWAS may also uncover signs of selection in additional traits. Four of the six traits for which we could not reject neutrality (asthma, gallstones, glaucoma, uterine fibroids) had fewer genetic associations than most other traits. Interestingly, IBD and hypothyroidism showed only weak or non-significant signatures of selection despite having substantially more associated variants, indicating that low power may not be the only issue. Selection associated with liability may therefore be lower for these traits than for other diseases. It is also possible that, for IBD, a reduced ability to detect low-frequency large-effect variants results from a skew towards common variants in the genotyping array or the fact that IBD is partly a mixture of Crohn’s disease and ulcerative colitis [65, 66].

Generally, traits are consistent with two forms of stabilizing selection: single-trait or pleiotropic. Numerous lines of evidence point to widespread pleiotropy for trait-associated variants [67, 68, 69], but these observations do not necessarily rule out single-trait models of stabilizing selection. The pleiotropy considered here refers to a large number of mutationally uncorrelated fitness dimensions [53, 37], which may not be satisfied if pleiotropic variant effects occur through narrow genetic pathways. Earlier work has argued that quantitative traits are under pleiotropic, not single-trait stabilizing selection [37], using a variance-contribution test to distinguish those models. Here, we use allele frequencies conditional on effect size to compare a broader set of selection models. We note that the variance-contribution test has limited ability to discriminate between forms of selection [37] (Supplement H). Still, in the original analysis, pleiotropic stabilizing selection models had higher likelihoods than single-trait models for all 13 quantitative traits and seven of the eight diseases with significant evidence of selection.

It is possible that the distinction between the single trait and pleiotropic models in our analysis is overstated. We simulated GWAS data under the two models and found much more limited power to distinguish between them (Supplement C.1, Figures S34–S36). Pleiotropic models also tended to produce unrealistically high estimates of selection coefficients for associated variants, especially in the original GWAS analyses (Figures S22, S24, S26, and S28; Supplement B.7). We speculate that this bias may be remedied by explicit modeling of the underlying distribution of variant effects [38] (Supplement B.7). It may be that other sources of heterogeneity in allelic effect sizes and frequencies (ascertainment, stratification, etc.) are better absorbed into the pleiotropic likelihood. For instance, it is difficult to model uncertainty in effect sizes and frequencies due to linkage disequilibrium, which is evident from variation among independent GWAS studies in different populations [70] (some variance may be explained by gene-environment interactions [71]). Neither simulations of this process nor our attempts to match effect sizes from independent populations and GWAS strongly favored pleiotropic models (Supplements G,C.3; Figures S41,S20). Alternatively, our inference may correctly point to a high degree of pleiotropy, but the relationship between a variant’s effect on the focal trait and its effects on other fitness-relevant traits that may be more heterogeneous within any given GWAS than current parameterization allows [72].

Two lines of evidence suggest that inferred stabilizing selection results from actual phenotypic selection to maintain intermediate trait values, whether many pleiotropic traits per variant or one at a time. Frequencies were more consistent with a model where selection decreases for high MAF variants (underdominance) and with selection coefficients that are proportional to squared effect sizes, on average. While this consistency is encouraging, neither is sufficient to demonstrate actual historical fitness effects associated with the current human phenotypes for which GWAS was performed. Even in the single-trait case and supposing real phenotypic selection, the object of that selection may be a genetically correlated trait [73, 74]. This is most readily apparent in the case of disease traits, where disease status itself, strictly speaking, cannot be under stabilizing selection. Liability, a separate trait, must be postulated as something capable of taking intermediate values.

If unconstrained by a need to model phenotypes, it is always possible to design a model of “apparent” selection that matches the variant-to-fitness function implied by a model of directional or stabilizing selection. If one’s goal is simply to parameterize that relationship, this is not necessarily a problem. Many useful inferences have been made using *ad-hoc* models of selection, though explicit pleiotropic stabilizing models appear to have the potential to substantially improve goodness-of-fit [38]. We are left to ask what it would actually look like to have a pleiotropic model, generating purifying selection against derived causative variants but without stabilizing selection on the focal or other traits. Such alternative models are not currently well-defined and additional theoretical work is needed. One possibility would consist of a large number of sub-traits under directional mutation-selection balance, but whose impact (per subtrait) on the focal trait are balanced between positive and negative changes. For example, genome-wide selection might favor increased protein stability, while decreased stability of individual proteins could have either positive or negative trait effects depending on position in pathways.

Pleiotropic stabilizing models, often favored by likelihood, also do not imply that the focal phenotype makes a large direct contribution to fitness. In the high-dimensional setting, variants with large effects on the focal trait are expected to have large aggregate effects on many other selected traits. The observed relationship between effect size and selection therefore arises even when fitness consequences are dominated by pleiotropic dimensions. While “apparent” has been used to describe models where negatively selected pleiotropic effects induce the appearance of stabilizing selection on an otherwise neutral trait, it is not clear that this is not also a fitting description of the multidimensional pleiotropic model when the number of selected dimensions becomes sufficiently large.

The finding of selection against some protective variants has several implications. First, it helps explain disease prevalence and variation in traits. Diseases, including those which appear to be post-reproductive, do not have a neutral genetics basis. For selection to decrease the genetic component of disease risk would require it to increase the prevalence of low-frequency protective genetic variants—which our results show has not been a consistent feature of human evolution.

Second, this selection reinforces the importance of deep phenotypic assessment in genetics-guided drug discovery [75]. Efforts to identify potential drug targets often examine existing genetic variation—”experiments of nature”—for protective variants [76, 77]. Selection against these variants may indicate that lowering the risk of a disease is inherently associated with a fitness cost or trade-off at the level of population evolution. Alternatively, even if lowering disease risk in isolation is itself associated with minor costs, it may be that all possible genetic routes (and corresponding drug targets) to doing so lead to harmful changes in other, seemingly unrelated, traits. The single-trait model of stabilizing selection might suggest the first of these scenarios, while the pleiotropic model might suggest the latter. This perspective could ultimately prove helpful in predicting whether the other phenotypic effects associated with a protective variant will lead to disqualifying risk.

As the sizes of GWAS grow, more traits will have a sufficient number of associated variants for the type of analysis presented here. Indeed, trumpet plots are becoming a standard part of the presentation of well-powered GWAS results. When possible, selection analyses like those presented here may provide useful insight into the mode and strength of selection on identified variants. Combining better-powered studies with more advanced models may also allow us to better distinguish between models of selection that produce similar genetic architectures, expanding on the scope of this study. These models may leverage sub-significant variants, combine multiple traits, leverage linkage disequilibrium between causal variants, or compare across populations [49]. Stabilizing selection models make explicit predictions that can be tested. For instance, variants with greater pleiotropy should, all else being equal, experience stronger selection, and haplotypes carrying opposite effect variants should be found at higher frequencies than those with concordant effect variants [78, 79].

We have shown that, across a broad array of traits, selection is common, but does not appear to have acted directionally over extended periods of evolutionary time. Instead, the genetic architecture of these traits is consistent with forms of stabilizing selection, indicating that selection has kept trait values and disease liabilities at intermediate levels. The extent to which this reflects direct selection on the focal trait or liability, rather than selection on pleiotropic effects of the same variants, remains unclear. This does not rule out adaptive shifts, but implies that they were accomplished without major changes to genetic architectures and did not result in large-scale fixation and the establishment of a strong mutational bias. Our results are therefore not inconsistent with recent findings of pervasive adaptive evolution in recent Eurasian history [80]. Allele frequency changes are substantial enough to be detectable in ancient DNA time series, but subtle enough to appear as noise in the smile-shaped distribution of GWAS hits and fail to dominate signals of stabilizing selection left over longer evolutionary time scales. Our findings help explain the high prevalence of some diseases and health-associated traits and raise important questions about the relationship between these traits and fitness.

## Data availability

Data are available upon publication through the figshare associated with this paper.

## Code availability

All code written for this project is publicly available at https://github.com/emkoch/smilenfer

## Supplemental Methods and Figures

### A GWAS Summary Statistics and Pipeline

#### A.1 GWAS sources

The original analysis was performed using the set of traits from UK Biobank and FinnGen.

##### A.1.1 Pan-UK Biobank [81]

Height, body mass index (BMI), LDL cholesterol, HDL cholesterol, diastolic blood pressure, systolic blood pressure, triglycerides, urate levels, red blood cell count (RBC), white blood cell count (WBC), grip strength, forced vital capacity (FVC), and pulse rate.

##### A.1.2 Meta-analysis of Pan-UK Biobank and Finngen [82]

Arthrosis, asthma, diverticulitis, gallstones, glaucoma, hypothyroidism, malignant neoplasms, uterine fibroids, and varicose veins.

##### A.1.3 Biobank Japan [58]

Height, body mass index, LDL cholesterol, HDL cholesterol, diastolic blood pressure, systolic blood pressure, triglycerides, red blood cell count, asthma, breast cancer, coronary artery disease, gall-stones, type II diabetes, uterine fibroids

##### A.1.4 Million Veteran Program [63]

Height, body mass index, LDL cholesterol, HDL cholesterol, diastolic blood pressure, systolic blood pressure, triglycerides, red blood cell count, white blood cell count, arthrosis, asthma, breast cancer, coronary artery disease, diverticulitis, gallstones, glaucoma, hypothyroidism, inflammatory bowel disease, type II diabetes, varicose veins.

##### A.1.5 Trait-targeting GWAS

Breast cancer [83], coronary artery disease (CAD) [84], inflammatory bowel disease (IBD) [85], schizophrenia (SCZ) [86], and type II diabetes (T2D) [87].

#### A.2 GWAS processing

GWAS summary statistics were taken from the studies referenced in the previous section. All variants other than biallelic SNPs were removed, and the 24,000,000 - 36,000,000 bp region of chromosome 6 was removed in order to avoid complications from the extended linkage disequilibrium and unusual selection properties of the human leukocyte antigen (HLA) region.

#### A.3 Selecting variants for analysis

Each plot and model test uses variants that are approximately linkage-independent. We first selected variants using a stepwise procedure. The most significant variant is selected, and all other variants within 100 kb are removed from the data. The most significant variant remaining is selected, and neighboring variants are again removed. This process is repeated until no two significant variants fall within 100 kb of one another. The stepwise method prioritizes variants by their original significance; prioritizing based on ASH-correct significance would create a bias toward variants with higher minor allele frequencies. After pruning summary statistics to a minimum distance of 100 kb, some variant pairs in high LD remained. We then calculated *r*^2^ between all pairs within 1 Mb and *r*^2^ *>* 0.2 using EUR samples in the TOPMed reference panel [88]. Any pairs for which TOPMed LD was not available were looked up in the 1000 Genomes EUR samples [89]. Conservatively, any remaining pairs for which both references failed were assumed to have *r*^2^ = 1. For each variant we then calculated the maximum *r*^2^ for any variant with a lower p-value and filtered out variants where this max *r*^2^ *>* 0.2. Traits based on Pan-UK Biobank and Finngen meta-analysis use a window size of 500 kb, as these are the publicly reported summary statistics.

#### A.4 Biobank Japan

We re-analyzed traits from Biobank Japan (BBJ) that overlapped the primary trait set and had sufficient numbers of variants for model fitting [58]. These included eight quantitative traits (height, BMI, LDL cholesterol, HDL cholesterol, diastolic blood pressure, systolic blood pressure, triglycerides, and red blood cell count) and six disease traits (asthma, breast cancer, coronary artery disease, gallstones, type 2 diabetes, and uterine fibroids). BBJ summary statistics were processed into the same format as the primary GWAS tables, with alleles polarized so that frequencies corresponded to the trait- or risk-increasing allele.

To account for LD among BBJ variants, we annotated each variant with the maximum LD to any more significant nearby variant within 1 Mb. LD was computed primarily using East Asian reference data from TOP-LD [88]. Variant pairs absent from TOP-LD were queried from 1000 Genomes reference data [89]. As for European-ancestry GWAS, we used an *r*^2^ reporting threshold of 0.2 and a minor allele-frequency cutoff of 0.01. Variants with max *r*^2^ ≥ 0.2 to a more significant variant were removed, and variants in the MHC region were excluded. Because BBJ was analyzed in a Japanese-ancestry cohort, likelihoods were calculated using the Japanese demographic model described in Section E. We computed a variance-contribution threshold corresponding to 5 × 10^−8^ using the median effective sample size for each BBJ trait.

For the main BBJ replication analysis, loci were clumped using a 500 kb window while prioritizing the variant with the largest trait- or risk-increasing effect size in each region. This high-effect clumping was used for the primary BBJ estimates shown in the main text. As sensitivity analyses, we repeated the BBJ fits using an otherwise identical 500 kb clumping procedure that prioritized the smallest *p*-value, and using 20 random clumpings in which one variant was sampled from each 500 kb region.

**Table S2:** Per-phenotype locus counts for the BBJ overlap set. Counts in the original column are the retained loci from the primary analyses. Counts in the BBJ column are the retained loci after the 500 kb high-effect clumping used for the main BBJ replication analysis.

| Phenotype | Original | BBJ |
| --- | --- | --- |
| *Quantitative traits* |  |  |
| BMI | 1078 | 86 |
| Diastolic blood pressure | 584 | 22 |
| HDL cholesterol | 788 | 40 |
| Height | 3260 | 459 |
| LDL cholesterol | 583 | 30 |
| Red blood cell count | 997 | 93 |
| Systolic blood pressure | 928 | 37 |
| Triglycerides | 625 | 28 |
| *Disease traits* |  |  |
| Asthma | 98 | 20 |
| Breast cancer | 204 | 8 |
| Coronary artery disease | 59 | 40 |
| Gallstones | 87 | 15 |
| Type 2 diabetes | 247 | 103 |
| Uterine fibroids | 73 | 17 |

#### A.5 UK Biobank SuSiE-X fine-mapping

We used published SuSiE-X fine-mapping results for UK Biobank quantitative traits [62]. Eight traits overlapped the original UK Biobank quantitative trait set and had available fine-mapped credible sets: BMI, diastolic blood pressure, HDL cholesterol, height, LDL cholesterol, systolic blood pressure, triglycerides, and white blood cell count. For each trait, we used the SuSiE-X credible-set table containing European-ancestry effect-size estimates, standard errors, allele frequencies, and posterior inclusion probabilities. Alleles were polarized so that frequencies corresponded to the trait-increasing allele.

To propagate uncertainty in the identity of the causal variant, we generated 20 replicate datasets for each trait. In each replicate, posterior inclusion probabilities were normalized within each SuSiE-X locus and credible set, and one variant was sampled according to its normalized posterior inclusion probability. Rows without an assigned SuSiE-X credible set were retained as single-variant loci with sampling probability one. We used the fine-mapped effect-size estimates directly, without shrinkage. After sampling, we calculated variance contributions from the sampled effect sizes and trait-increasing allele frequencies. We used the median effective sample size from the corresponding original UK Biobank GWAS to define the variance-contribution threshold associated with 5 × 10^−8^, retained sampled variants with *v > v*^∗^ and trait-increasing allele frequency between 0.01 and 0.99, and fit the same selection models as above using the European demographic model.

To better understand the differences between the original clumped UK Biobank analysis and the SuSiE-X re-analysis, we examined the extent to which loci from the original GWAS were represented in the reported fine-mapping results as a function of their distance above the ascertainment boundary. For each trait, we grouped original loci by 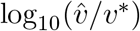, where 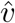 is the estimated variance contribution of the original locus and *v*^∗^ is the threshold corresponding to 5 × 10^−8^, and calculated the fraction of loci in each bin that mapped to at least one reported credible set (Figure S16). Loci close to the ascertainment boundary were found less often in in the fine-mapping output, whereas loci with larger variance contributions relative to the boundary were retained much more consistently. This likely reflects the exclusion of loci with diffuse posterior support (low PIP across many variants) and potentially duplications of single causative variants due to insufficient LD filtering. Because loci near the ascertainment boundary contribute information about the lower edge of the smile and about the extent to which moderately selected variants are represented, their preferential exclusion makes the fine-mapping analysis a conservative comparison to the original clumped analysis and likely contributes to the weaker support for pleiotropic stabilizing selection observed in the SuSiE-X results.

#### A.6 Million Veteran Program fine-mapping

We used fine-mapping results from the Million Veteran Program (MVP) phenome-wide association study [63]. The released table contained variant-level fine-mapping results across multiple ancestry-specific analyses. We restricted to the European-ancestry analysis and to traits annotated as PheCodes. The traits analyzed were atrial fibrillation, basal cell carcinoma, coronary atherosclerosis, diverticulosis and diverticulitis, glaucoma, gout, hyperlipidemia, hypertension, hypothyroidism, prostate cancer, and type 2 diabetes. For each trait, we retained rows matching the reported PheCode description, and treated each combination of reported locus and population-specific signal identifier as a fine-mapped signal. We used the European-ancestry effect-size estimate, standard error, effect-allele frequency, and credible-set-level posterior inclusion probability. Alleles were polarized so that frequencies corresponded to the risk-increasing allele.

To account for uncertainty in the causal variant within each fine-mapped signal, we generated 20 replicate datasets per trait. In each replicate, one variant was sampled from each signal according to its reported credible-set-level posterior inclusion probability. We used the MVP fine-mapped effect-size estimates directly, without shrinkage. For each trait, the variance-contribution threshold was defined using the median effective sample size estimated from the MVP fine-mapped variants and the MVP phenome-wide significance threshold 4.6 × 10^−11^. After sampling, variants were restricted to effect-allele frequency between 0.01 and 0.99, polarized by the sign of the MVP effect-size estimate, and retained only if their variance contribution exceeded this threshold. We then fit the same selection models as above using the European demographic model.

**Table S3:** Per-phenotype matched genome-wide significant locus counts in non-European MVP populations. Counts were obtained by intersecting the retained project loci with population-specific MVP fine-mapped signals significant at *P* ≤ 4.6 × 10^−11^. For continuous traits, counts use the corresponding mean phenotype rows in the MVP fine-mapping table.

| Phenotype | AFR | AMR |
| --- | --- | --- |
| *Quantitative traits* |  |  |
| BMI | 8 | 4 |
| Diastolic blood pressure | 2 | 0 |
| HDL cholesterol | 16 | 14 |
| Height | 25 | 29 |
| LDL cholesterol | 5 | 5 |
| Red blood cell count | 4 | 6 |
| Systolic blood pressure | 4 | 7 |
| Triglycerides | 7 | 10 |
| White blood cell count | 8 | 4 |
| *Disease traits* |  |  |
| Arthrosis | 0 | 0 |
| Asthma | 1 | 0 |
| Breast cancer | 0 | 0 |
| Coronary artery disease | 1 | 2 |
| Diverticulitis | 0 | 0 |
| Gallstones | 0 | 0 |
| Glaucoma | 1 | 1 |
| Hypothyroidism | 1 | 5 |
| Inflammatory bowel disease | 0 | 0 |
| Type 2 diabetes | 11 | 3 |
| Varicose veins | 0 | 0 |

### B Model and Likelihood

#### B.1 Risk allele frequencies are robust to linkage disequilibrium

Most trait-associated variants in GWAS are not themselves causative but arise due to tagging of a causative variant by LD; furthermore, fine mapping does not generally identify single causative variants with high confidence. Our selected LD-independent variants thus serve as proxies for causative variants driving associations at the respective loci. A key concern is then how well the proxy allele frequencies and effect sizes used in inference approximate those of the respective causative variants. In particular, while it is intuitive and well known that variants in tight LD have similar minor allele frequencies and similar absolute effect sizes, a question remains as to sign, namely whether the *risk* allele frequency might “flip”, for example from around 0.1 to around 0.9. The answer is that it does not. We provide both an analytic argument and an empirical demonstration.

The analytic argument rests on the observation that, with respect to a given trait, *a causative trait-increasing allele and a tagging trait-increasing allele are in positive LD*. In detail, let *X*_*a*_, *X*_*b*_ be causative and tagging variants respectively, coded {0, 1, 2} as usual with respect to reference/alternate alleles A/a, B/b defined so that the alternate allele is increasing for trait *Y* in the sense that the GWAS regression coefficient *β*_*a*_ = Cov(*Y, X*_*a*_)*/* Var(*X*_*a*_) *>* 0 and similarly *β*_*b*_ *>* 0 (for simplicity here we omit additional covariates and all sampling noise in the GWAS coefficient estimates). The signed LD is the Pearson correlation coefficient

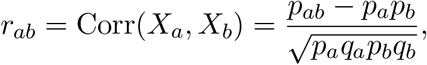

where *p*_*a*_ = 1 − *q*_*a*_, etc. are allele frequencies and *p*_*ab*_ is the frequency of the ab haplotype. The frequency bounds derived here are given in terms of haplotype frequencies because these determine the sign of LD. Writing *Y* = *β*_*a*_*X*_*a*_ + *ε* with Cov(*X*_*a*_, *ε*) = Cov(*X*_*b*_, *ε*) = 0 (for simplicity here we assume a single causal variant per locus), one finds 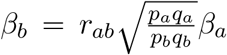. It follows that *r*_*ab*_ *>* 0. Incidentally, we note that the effect size relationship simplifies nicely on the variance explained scale: with 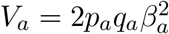 and 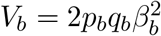, we have 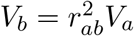.

The classical bounds on LD in terms of marginal allele frequencies may be summarized

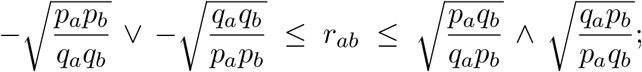

these represent four inequalities, two of which are nontrivial in a given instance, and which saturate respectively when one of the four haplotype frequencies *p*_*ab*_, *p*_*AB*_, *p*_*aB*_, *p*_*Ab*_ drops to zero. In particular, *r*_*ab*_ = 1 only when *p*_*a*_ = *p*_*b*_, while *r*_*ab*_ = −1 only when *p*_*a*_ = *q*_*b*_. The former extends quantitatively to the case *r*_*ab*_ *>* 0, where we obtain the bounds

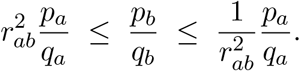

Inverting the function *p* 1→ *p/*(1 − *p*), which is monotonic for 0 *< p <* 1, via *t* ↦ *t/*(1 + *t*) yields bounds on *p*_*b*_ in terms of *p*_*a*_ and *r*_*ab*_. For example, with *p*_*a*_ = 0.2 and 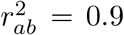 we find 0.18 ≤ *p*_*b*_ ≤ 0.22; with *p*_*a*_ = 0.2 and 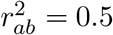 we find 0.11 ≤ *p*_*b*_ ≤ 0.34.

For the empirical demonstration, we looked at actual GWAS results for human height, selecting a number of significantly associated loci at random. For each locus we selected the sentinel (most significantly associated) variant and then considered all variants in LD *r*^2^ *>* 0.6 with the sentinel variant. For these variants we plot variance explained against trait-increasing allele frequency. As one can see in the sample loci in Figure S1, the plots for each locus cluster tightly along both axes; in particular we find no “shadow cluster” at the complementary allele frequency. Note that this demonstration recapitulates none of the simplifying assumptions from the analytic argument (covariates, causal architecture).

This analysis shows that tagging should not flip the trait- or risk-increasing allele frequency from one side of 0.5 to the other. The broader question of whether lead variants adequately approximate causal variants for model fitting is addressed separately through graphld simulations and fine-mapping analyses (Supplement C.3; Figures S39–S41) and through fine-mapping analyses that sample variants according to posterior inclusion probabilities.

#### B.2 Observables and notation

The pipeline described in section A yields a set of approximately independent loci. Each locus has the accompanying summary statistics of allele frequency, estimated effect size, standard error, and *p*-value. These are given for the variant at that locus with the smallest *p*-value (sentinel variant). The allele frequency available for each locus is that of the trait-increasing or risk-increasing allele (RAF) (section B.1), which we denote using *y*. We treat the derived allele frequency, denoted by *x*, as a latent variable. The effect size estimated by the GWAS is denoted by 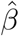, the posterior mean after shrinkage by 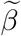, and the true latent effect size by *β*. Standard errors and *p*-values are denoted by SE and *p*.

The variance contribution of a locus is defined as

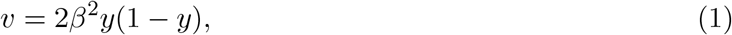

where we may also substitute *x* for *y*. Similarly, let 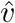 be the GWAS-estimated variance contribution. Ascertainment in GWAS based on a *p*-value threshold can be approximated as a variance-contribution threshold. If *v*^∗^ is the threshold for a particular study and *p*-value, variants face an allele frequency threshold conditional on their effect size

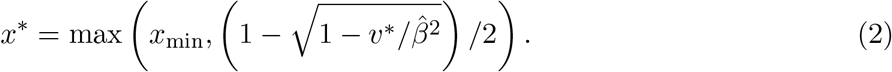

Even if the GWAS is powered to discover some rare variants, we apply the minor allele frequency threshold *x*_min_.

#### B.3 Choosing the variance contribution threshold

In order to truncate the frequency distribution and ultimately calculate the likelihood contribution of each locus, we need to compute the variance contribution threshold *v*^∗^. The relationship between variance contributions and *p*-values is approximately

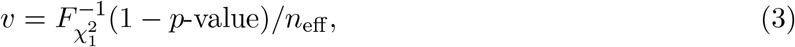

where 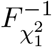 is the inverse CDF of a chi-squared distribution with one degree of freedom and *n*_eff_ is the effective sample size of the GWAS. In order to find a *v*^∗^ corresponding to a particular *p*-value threshold, we first calculated the median *n*_eff_ for the GWAS by using the observed set of {*v*_*i*_, *p*-value_*i*_} value to calculate a local *n*_eff,*i*_ for each genome-wide significant variant in the processed summary statistics: 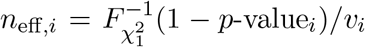 Using the median *n*_eff_, we compute *v*^∗^ for various *p*-value thresholds using equation (3).

The relationship between *p*-values and effective sample sizes for each trait is shown in Figure S6. A single effective sample size is not truly sufficient because the GWAS regressions used to estimate effect sizes and compute *p*-values include confounding variables as well as genetic structure (e.g. principal components). Variants that are correlated with these covariates will require a large sample for their effects to be detected and therefore have a lower effective sample size. We use this procedure to generate a *v*^∗^ that roughly corresponds to a *p*-value threshold. In addition to the variance contribution threshold, we also removed variants that do no satisfy the corresponding *p*-value threshold. Applying the *p*-value threshold removes variants with large standard errors relative to their estimated effect sizes, while making sure that they fall within the frequency range defined by *v*^∗^.

#### B.4 The allele frequency distribution under different models of selection

We first describe the structure of the model in the absence of any uncertainty associated with measurement through GWAS. This provides intuition and is also used in simulations to demonstrate the theoretical limitations of our approach. Our inferences are based on the RAF distribution, conditional on effect size

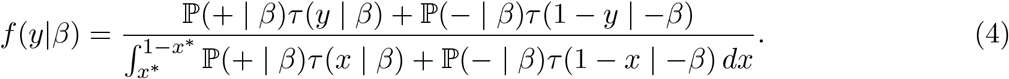

This distribution accounts for the fact that the ancestral/derived status of the trait-increasing allele is unknown. ℙ(+ | *β*) is the probability that a new mutation with effect size magnitude |*β*| is trait-increasing, while ℙ(− | *β*) = 1 − ℙ(+ | *β*) is the probability this effect is trait-decreasing. *τ* (*x* | *β*) is the site-frequency spectrum (SFS) of derived allele frequencies with (signed) effect size *β*. In an equilibrium population this is proportional to the sojourn time a new mutation spends at frequency *x*, while in more general scenarios it gives us the expected number of variants at this frequency. How the SFS depends on *β* varies between different models of phenotypic selection. The denominator normalizes by the expected number of variants in the discoverable range defined by *v*^∗^ to yield a probability density on the RAF. Normalization also removes any dependency on mutation rate under an infinite-sites model.

Describing a model of selection amounts to specifying ℙ(+ | *β*) and *τ* (*x* | *β*). We do this for five models which we hope capture a broad range of possible phenotypic selection scenarios. The SFS, *τ* (*x* | *β*), is determined by the first two moments of the change in allele frequency. Directional selection at the phenotypic level generates additive selection at the genic level, while stabilizing selection at the phenotypic level generates underdominant selection at the genic level. We include one selection coefficient for each effect

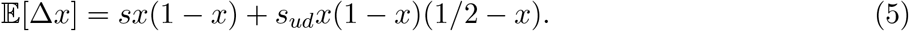

The variance is given by the standard formula for the Wright-Fisher model

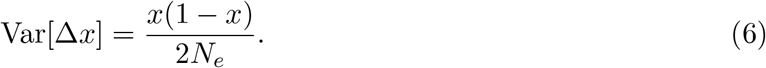

In practice, true effect sizes are not unknown. All inferences here proceed from specifying a value 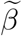 to stand in for the true value of *β* while 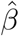 determines the ascertainment threshold 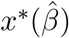:

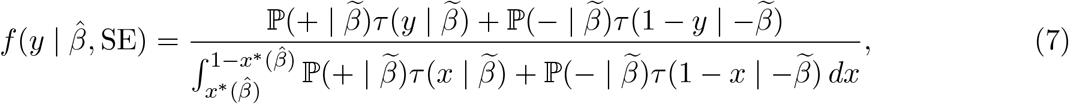

where SE is the standard error reported by the GWAS. 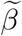 therefore determines the selection acting on the variant. The main text presents results using posterior mean values for 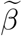, after fitting the distribution of effect sizes using `ashr` [55]. A more detailed discussion of this choice and other alternatives is given in Section D.

##### B.4.1 Neutral

For a neutral model with a total absence of selection on the focal trait, we specify both that there is no relationship between effect sizes and selection coefficients and that all variants discovered by GWAS have selection coefficients of zero. We therefore have *τ* (*x* | *β*) = *τ* (*x*) for any observed *β*, and *τ* (*x*) ∝ 1*/x* at equilibrium. There is a question of how to set the mutational balance between + and − alleles under neutrality. We opt for ℙ(+ | *β*) = ℙ(− | *β*) = 0.5 because strong biases here would lead to rapid changes in the mean trait value over longer evolutionary periods due solely to mutational bias.

##### B.4.2 Directional selection

For a model of directional selection on the focal trait we consider the scenario where the optimum trait value exists outside of the range of phenotypes present in the population. Each mutation experiences selection proportional to its effect size:

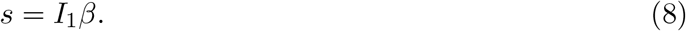

In an equilibrium population the probability a site with effect size *β* is fixed for the risk-decreasing allele is equivalent to the probability that a new mutation with that effect size is risk-increasing. In the diffusion limit this can be obtained by balancing substitution fluxes between the two fixation states. Let *u*(*s*) be the fixation probability of a new mutation with additive selection coefficient *s*, where

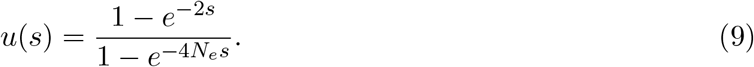

If mutations in the two directions occur at equal rates, then equilibrium requires

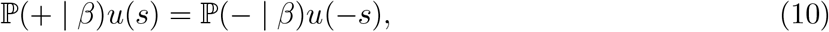

where ℙ(− | *β*) = 1 − ℙ(+ | *β*). Solving gives

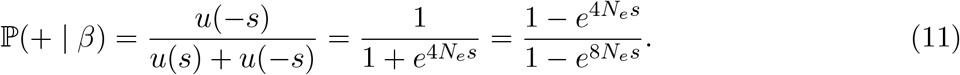

Thus mutational bias arises because sites are more often fixed for the allele favored by directional selection, so new mutations are more likely to point in the disfavored direction [90]. Under a directional model of selection on disease liability the optimal genotype would contain no risk-increasing alleles. The population mean therefore sits off the optimum phenotype because new mutations continually introduce risk-increasing alleles, generating a polygenic mutational bias toward disease. We combine this with the standard expression for the SFS under additive selection or one obtained numerically to account for nonequilibrium demography (section E).

For numerical work it is convenient to rewrite this probability in algebraically equivalent forms that avoid cancellation near *s* = 0 and large exponentials when *s >* 0. In particular,

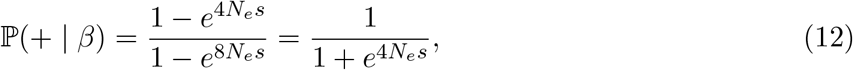

with ℙ(+ | *β*) = 1*/*2 at *s* = 0. We therefore evaluate it using the equivalent piecewise representation

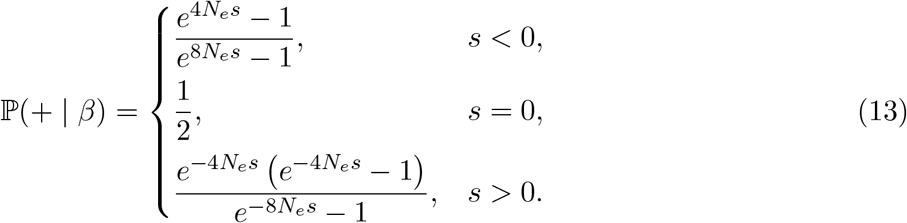

These forms are equivalent to the expression above, but keep the computation well-behaved on both sides of zero.

##### B.4.3 Single-trait stabilizing selection

Stabilizing selection applies to the situation where the population mean for focal trait is very close to the optimal value. Stabilizing selection will be approximately one-dimensional when only the focal trait and those strongly correlated with it are under selection. Each mutation receives a selection coefficient proportional to its squared effect size

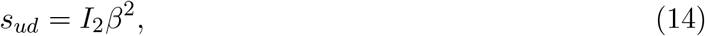

and the form of this selection is underdominant rather than additive. We constrain the sign of *I*_2_ so that selection is never overdominant. Because selection is symmetric with respect to the sign of *β* we have ℙ(+ | *β*) = ℙ(− | *β*) = 0.5. This also agrees with the intuition that if the population mean is close to the optimum there should not be a strong mutational bias

##### B.4.4 Combined directional and single-trait stabilizing selection

We also consider a model where both directional and stabilizing selection at the trait level make a contribution to the selection on individual alleles. Such a model may apply if the population mean is off the optimum value but not far enough that stabilizing selection can be ignored. It is less well-justified than the other models considered but could be useful in cases where selection is relatively strong while mutational asymmetry is mild. We simply allow alleles to have both an additive and underdominant selection coefficient. The stabilizing component of selection remains symmetric in *β*, so P(+ | *β*) is determined only by the additive selection coefficient.

##### B.4.5 Pleiotropic stabilizing selection

The single-trait model of stabilizing selection arises when the variants influencing the focal trait, and those traits highly correlated with it, meaningfully affect no other traits under stabilizing selection. Simons *et al*. [37] developed a model where variants affect many other traits under stabilizing selection that are mutationally uncorrelated with the focal trait. We use the terms single-trait and pleiotropic to refer to the one-dimensional and high-dimensional limits of this model. In the limit as the number of uncorrelated traits becomes large, the effect size on the focal trait conditional on the selection coefficient is

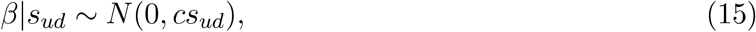

where *c* depends on the number of traits and the strength of selection on each. Because our likelihood is formulated conditional on the effect size, to use the high-dimensional model of pleiotropy we must make assumptions about the distribution of fitness effects of new mutations (DFE). Rather than attempting to also fit this distribution using the data, we put a flat prior on log(*s*_*ud*_)

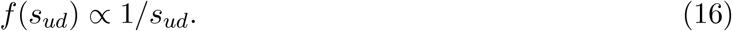

This yields a Lévy distribution on *s*_*ud*_ conditional on *β*

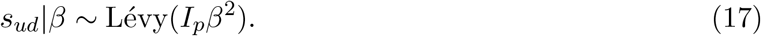

The parameter *I*_*p*_ is analogous to the selection intensities *I*_1_ and *I*_2_ defined above for directional selection and single-trait stabilizing selection. We can connect this parameter back to the average fitness effects of mutations with particular trait effects through

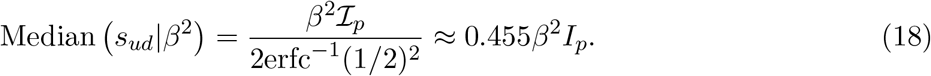

To obtain the SFS we integrate over the distribution of selection coefficients conditional on the effect size

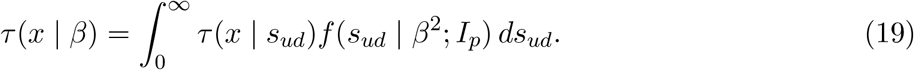

By the same logic given for one-dimensional stabilizing selection, we also assume equal proportions of trait-increasing and trait-decreasing mutations.

#### B.5 Likelihoods and model comparison

For each trait, the data consist of a set of RAFs and effect sizes 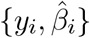 after filtering based on a variance contribution, and possibly also *p*-value, threshold. Depending on the model, the parameters Θ are either empty (neutral), *I*_1_ (directional), *I*_2_ (single-trait / one-dimensional stabilizing), (*I*_1_, *I*_2_) (directional plus stabilizing), or *I*_*p*_ (pleiotropic / high-dimensional stabilizing). Assuming independence among loci, conditional on model parameters, the likelihood is

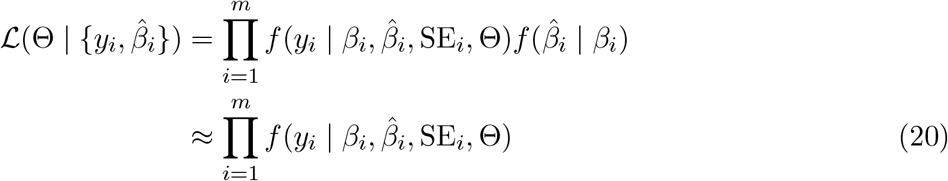

where *m* is the number of loci passing the threshold. This expression results because only frequencies directly depend on selection parameters after conditioning on variant effects, and because we assume that *β*_*i*_ is actually known, either as 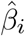 or a shrunken value. *f* here is implicitly truncated to the range to the discoverable range.

Since the models only contain either one or two parameters, we fit them by direct numerical maximization of the likelihood. For each model, likelihoods were evaluated using precomputed site frequency spectrum grids, with interpolation over allele frequency and selection coefficients as described below. We optimized the negative log-likelihood using the Nelder-Mead algorithm as implemented in `scipy.optimize.minimize` [91]. For the single-trait stabilizing model, we optimized log_10_(*I*_2_) over the interval [−8, 2]. For the pleiotropic stabilizing model, we optimized log_10_(*I*_*p*_) over 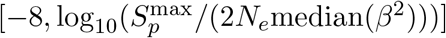, where 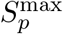 is the largest scaled selection coefficient represented in the precomputed pleiotropic grid. For the directional model, we optimized *I*_1_ directly over [−10, 10], allowing both positive and negative values. For the combined directional and single-trait stabilizing model, we optimized *I*_1_ and log_10_(*I*_2_) over [−10, 10] and [−8, 2], respectively.

All models considered here converge to neutrality as the selection intensity parameters become small. We therefore use likelihood ratio tests to compute *p*-values for each model relative to the neutral model. To compare models of selection we directly compare likelihoods by computing Akaike information criteria.

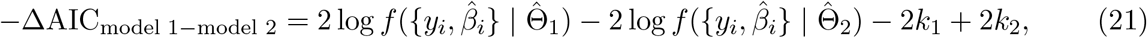

where 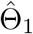 and 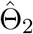 are the maximum likelihood parameter values and *k*_1_ and *k*_2_ are the numbers of free parameters. ΔAIC values presented in figures are on the scale of twice the log-likelihood differences.

#### B.6 Uncertainty and comparison of selection intensity parameter estimates

We quantified uncertainty in the fitted selection intensity parameters for the original trait set in two ways (Figure S14). First, we calculated Wald confidence intervals from the curvature of the log-likelihood surface around the maximum-likelihood estimate. For *I*_2_ and *I*_*p*_, likelihoods were evaluated on a local grid of parameter values. Resulting grids were then smoothed, and used to estimate the Fisher information. Confidence intervals were then calculated on the log scale and transformed back to the original parameter scale. Second, we generated 1,000 bootstrap datasets for each trait by resampling retained loci with replacement and re-fitting the selection model. Bootstrap intervals were taken as the 2.5th and 97.5th percentiles of the resulting parameter estimates.

Estimates of *I*_2_ and *I*_*p*_ are only partially comparable to each other and across traits. The median selection coefficient under the pleiotropic model is equivalent selection under the one-dimensional model when *I*_*p*_ ≈ 0.455*I*_2_. However, predicted site frequency spectra show other qualitative deviations not captured by this comparison (Figure S45). Parameter estimates between traits depend on the scale of the estimated effect sizes, which are not guaranteed to be consistent. Model comparisons were implemented so as not to be sensitive to the scale of GWAS estimates.

#### B.7 Variant-level selection coefficients

To visualize the strength of stabilizing selection implied by the fitted models, we computed a variant-level scaled underdominant selection coefficient, *S*_*ud*_ = 2*N*_*e*_*s*_*ud*_, for each retained locus. Under the single-trait stabilizing model, this quantity is determined directly by the fitted selection intensity and the variant effect size:

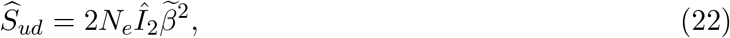

where 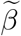 is the shrunk effect estimate used for inference.

Under pleiotropic stabilizing selection, the fitted parameter *Î*_*p*_ implies a distribution of selection coefficients rather than a single value,

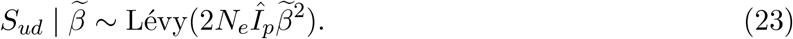

For visualization, we combined this distribution with the underdominant site-frequency spectrum at the observed risk-allele frequency *y*, so that

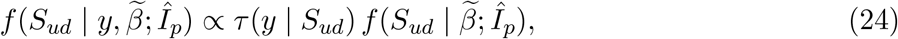

where *τ* (*y* | *S*_*ud*_) is the site-frequency spectrum under underdominant selection in the demographic model used for the corresponding dataset. The value displayed for each variant in the pleiotropic panels was the median of this conditional distribution,

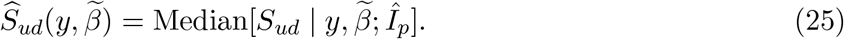

Thus, for the single-trait model each plotted value is the direct selection coefficient implied by the fitted model, whereas for the pleiotropic model each plotted value is a frequency-informed summary of the posterior distribution of selection coefficients for that variant.

We plotted estimated variant-level selection coefficients for all stabilizing selection models fit in Figure 3 (Figures S21–S28). Selection coefficients were scaled to a hypothetical ancestral effective population size of *N*_*e*_ = 10000 in order to provide a comparison point from very weak 2*N*_*e*_*s <* 1 to strong 2*N*_*e*_*s >* 100 selection. Selection coefficients implied by the models of stabilizing selection we fit spanned this entire range. For most traits, the highest-frequency, lowest-effect variants identified were inferred to be under very weak selection (*S*¡2). However, for some traits in the original data pleiotropic models estimated very strong selection for the weakest, most frequent variants. SCZ and BMI were the most extreme in this regard. Pleiotropic stabilizing models of selection also inferred stronger median selection coefficients than point estimates under the single trait model. We suspect that pleiotropic model estimates are unrealistically high because of the assumption of a flat distribution of fitness effects on log *s* used in the likelihood. This choice simplified inference, and did not appear to have a major effect on model comparison. However, it is likely that assuming strong selection occurs as frequently as weak inflates posterior estimates of selection per-variant.

Per-variant estimates of selection were also considerably smaller for fine-mapping data sets for both the single-trait and pleiotropic stabilizing models. As noted previously, fine-mapping data sets tended to not report variants near the ascertainment boundary, making this unsurprising. The truth likely lies somewhere between the raw-GWAS pleiotropic fits and those from fine-mapping, and methods which specifically model the underlying distribution of fitness effects are better suited to producing individual estimates of selection coefficients [38]. We ultimately chose not to take this approach due to the complicating effect on model comparison.

For visualization of the combined directional and stabilizing model, we used the maximum-likelihood estimates (*Î* _1_, *Î* _2_) together with the shrunk effect estimate 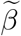 for each retained variant to compute both the scaled additive and underdominant selection coefficients,

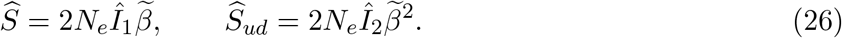

We then plotted variants in the same RAF-versus-effect-size layout as in the stabilizing-selection panels, with point color indicating the inferred stabilizing component 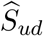 and the secondary y-axis indicating the corresponding directional component 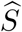 (Figure S29). Only traits for which the combined model had a lower AIC than the single-trait stabilizing model were shown. The sign of 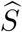 indicates the direction of the directional component, with negative values corresponding to selection against trait-increasing or risk alleles and positive values corresponding to selection against trait-decreasing or protective alleles.

### C Simulations

#### C.1 Trait simulations

To evaluate the ability of these statistics to distinguish between different models of selection we performed simulations designed to provide approximately realistic distributions of effect sizes and allele frequencies. Our simulation strategy used the prior distribution of effect sizes, ℙ(*β*), fit using ASH for a given trait. We fix the number of ascertained variants at some number *m* and simulate allele frequencies and effect sizes until it is reached. We first sample a set of effect sizes {*β*_*i*_} from ℙ(*β*) and compute the allele frequency distribution *f* (*y*|*β*_*i*_) (equation (4)) for the selection intensity parameters (*I*_1_, *I*_2_, *I*_*p*_) corresponding to the simulation. Rather than using the MAF cutoff *x*^∗^ corresponding to a variance contribution threshold *v*^∗^, we first apply a global MAF threshold of *x*^∗^ = 0.01. This is equivalent to the assumption that ℙ(*β*) represents the effect size distribution of all alleles with MAF greater than *x*^∗^. Using *f* (*y*|*β*_*i*_), we sample trait-increasing allele frequencies from the range (*x*^∗^, 1 − *x*^∗^), yielding a large set of {*y*_*i*_, *β*_*i*_}.

We take variants from the set of allele frequencies and effect sizes to approximate GWAS ascertainment using a variance-contribution threshold. For simulations without ascertainment bias, we simply apply *v*^∗^ to 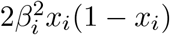, where *v*^∗^ corresponds to the given trait and a particular *p*-value threshold. Otherwise, we add GWAS noise to *β*_*i*_ using

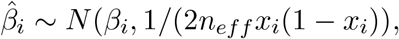

and we assume that the correct standard error 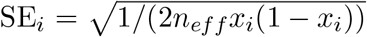 is available as a summary statistic. Variants are then added to the ascertainment set when 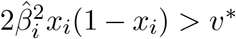. We add variants until *m* is reached, and draw new *β*_*i*_ from the prior and repeat the process if not enough variants from the current set pass ascertainment.

When calculating likelihoods using simulated data we face the choice of what to treat as the true effect size. In simulations without ascertainment bias this is always the true effect size *β*_*i*_. In simulations with ascertainment bias we calculate likelihoods using *β*_*i*_, 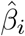, and the posterior mean calculated using the known ℙ(*β*) and SE_*i*_ for that simulation. Making calculations for each of these allow us to investigate the effects of ascertainment bias on allele frequencies and uncertainty in effect sizes somewhat separately. Simulation results were qualitatively similar for different traits and we present results only for IBD.

#### C.2 Simulations under Simons *et al*. (2025) DFE

Simons *et al*. [38] found that estimated DFEs for different quantitative traits have similar shapes. They presented a single shared distribution (SSD) that fit the data from all analyzed traits reasonably well (Figure S33). We performed simulations using this DFE to evaluate our model’s ability to discriminate between one-dimensional and high-dimensional stabilizing selection. We sampled s_ud_ from f_SSD_ then sampled a derived allele frequency using either a constant effective population size or a demographic model. Under the high-dimensional pleiotropy model [37], the effect size (*β*) conditional on s_ud_ is

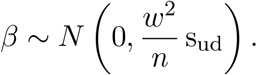

We transformed *β* so that variance contributions would be in units of scaled selection coefficients

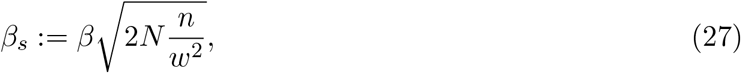

and

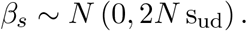

The variance contribution of an allele with effect size *β*_*s*_ is 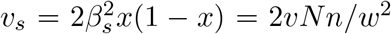. Again following Simons *et al*. [37], the expected variance contribution in trait units of a pleiotropic locus under strong selection is

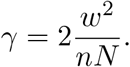

We model ascertainment by keeping alleles with a variance contribution exceeding some threshold *v*^∗^. To aid with interpretation, we choose this threshold relative to the expected contributions from strongly selected alleles:

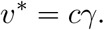

On the selection-transformed scale this threshold is

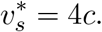

To simulate GWAS noise and ascertainment bias in the sample, we also need to simulate 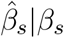. GWAS noise can be modeled as being approximately normally distributed so that

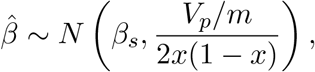

where *V*_*p*_ is the phenotypic variance in the population and *m* is the sample size. For transformed effect sizes this is

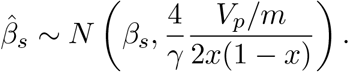

The approximate variance contribution threshold corresponding to a particular p-value *p*^∗^ is therefore

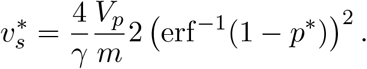

Since we have already specified that 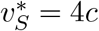, we can define a scaled effective sample size 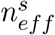 such that

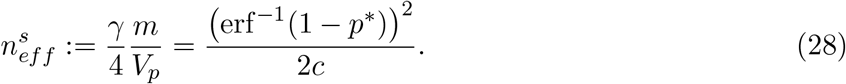

GWAS noise was then simulated as 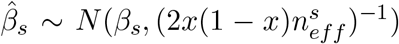 and alleles with 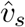 exceeding 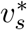 were retained as part of the sample. All of the above holds for the case of single trait (one-dimensional) stabilizing selection, 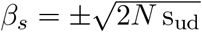, with trait-increasing and trait-decreasing alleles equally likely. For the demographic model of non-Finnish European history we let *N* be the ancestral population size.

To create a sample of {*y*_*i*_, *β*_*i*_} we used a similar approach to the trait-based simulations but instead first sampled selection coefficients from f_SSD_(s_ud_). This was weighted by the probability of observing an allele with that selection coefficient above the MAF threshold *x*^∗^ = 0.01: 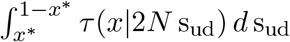. We then sampled a frequency from the truncated SFS, *β*_*i*_ from *N* (0, 2*N* s_ud_) and 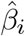 from 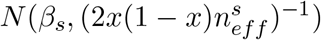. We computed likelihoods without ascertainment, and with ascertainment bias only treating 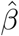 as the true effect size. We used a *p*-value threshold of 5 × 10^−8^, 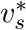 at (1, 3, 5, 7), and 500 ascertained variants in each simulation. Increasing 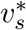 while holding the *p*-value threshold and number of ascertained variants constant corresponds to decreasing the power of the GWAS while increasing the mutational target size to ensure that a large number of strongly selected variants are ascertained. In simulations with ascertainment bias this also increased the effect of winner’s curse by lowering *n*_*eff*_.

DFE-based simulations were used to evaluate the power to distinguish the low- and high-dimensional models of stabilizing selection. All simulations showed strong evidence (−ΔAIC *>* 50) to reject a neutral model of selection on GWAS variants (Figure S35). The evidence for selection and power to detect the pleiotropy of stabilizing selection were higher in simulations using a constant population size (*N*_*e*_ = 10,000), likely due to the effect of the out-of-Africa bottleneck allowing alleles under moderate selection with moderate effect sizes to drift to higher frequencies (Figure S36). However, even under a constant population size a substantial fraction of simulations would fail to show strong evidence for the true pleiotropy model. We also used trait-based simulations to evaluate the power to detect the true pleiotropy model and found that it was limited until the overall evidence for selection became strong (*I*_2_ or *I*_*p*_ large, −ΔAIC *>* 100) (Figure S38).

The distribution *p*(s_ud_ | *β*) used to compute the likelihood in the pleiotropic (high-dimensional) model of stabilizing selection assumed a flat (improper) prior on log s_ud_. We checked whether results were robust to this choice by instead using the SSD as a prior and calculating likelihoods for DFE-based simulations. Changing the prior had little effect on the likelihood even though this was the prior used to generate the data (Figure S37).

#### C.3 Simulated tagging variants using graphld

Variants were simulated using the linkage disequilibrium graphical model (LDGM) framework implemented in python [92, 93]. The genome was divided into blocks based on linkage disequilibrium. Within a block, each variant was assigned a prior variance that was based on the alpha model: 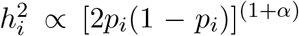 · weight. The weight is assigned based the relevant annotations for the variant in question, and their per-variant heritability enrichment [94]. This accounts for both the distribution of genomic annotations and the frequency-dependent architecture of complex traits. The simulated effect size of a variant was drawn from the normal distribution *β*_*i*_ ~ *N* (0, *h*^2^)*/m*, where *m* is the number of variants with simulated non-zero effects. The marginal effects were calculated using all regional *β* values and the linkage data contained in the LDGM.

For these simulations, we selected parameters based on LDL cholesterol, using published estimates of *h*^2^ [95, 96] and *α* [23]. For LDL cholesterol’s underlying polygenicity, which is difficult to estimate from data, we used a value that produced a simulated genetic architecture similar to the observed architecture of the trait. We produced 10 simulations of LDL under selection, and 10 simulations of a neutral LDL (*α* = 0).

The output of these simulations includes both the frequency and effect of the true causative variant, as well as the frequencies and marginal effects of lead variants in the simulated GWAS (Figure S39). We fit models to both of these sets and used this comparison to investigate the effects of uncertainty in both the frequency and effect sizes of variants on inferences of selection. In the lead-variant analysis, we used the genome-wide significant lead variants selected from each LD clump. In the causal-variant analysis, we used simulated causative variants passing the same filters. Because the number of retained lead and causative variants could differ, it was treated as a consequence of ascertainment bias to be tested.

### D GWAS and ascertainment bias

The truncated RAF distribution in equation (4) assumes that if a variant with true effect size *β* passes the minor allele frequency threshold *x*^∗^ defined by *v*^∗^ it will be observed in the GWAS sample. In reality, variants that pass either a *p*-value or variance contribution threshold will be subject to winner’s curse and estimates of their effect sizes 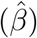 will be biased upwards. We developed strategies to partially account for the effect of winner’s curse while avoiding fitting the distribution of effect sizes and its dependence on allele frequency. We also performed simulations to investigate some of the impacts of ignoring this ascertainment bias.

The first strategy simply ignores winner’s curse and treats 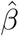 as if it were the true effect size. This strategy uses overestimates of effect sizes, and more so for low frequency variants with larger standard errors where winner’s curse will have a stronger effect (Figure S10). We simply substitute 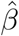 for 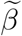 in equation (7).

#### D.1 Posterior effect size shrinkage

The second strategy treats the ASH posterior mean from the adaptive shrinkage (ASH) method implemented in the *ashr* package [55], as the true effect size, while still using 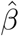 to calculate the frequency cutoff *x*^∗^. Ascertainment of GWAS variants can cause traits evolving neutrally to display signals of selection. As described in the main text, the concept of winner’s curse reflects that some variants have a perceived effect in a GWAS sample that is higher than their causal effect. These variants will be inherently more likely to be discovered in GWAS, and because most variants in a population are relatively rare, most variants identified through winner’s curse will also be rare, potentially causing or exaggerating a U-shaped curve.

This Bayesian method first constructs a model of GWAS variant effect sizes and standard errors—to balance computational tractability with sufficient information across a range of effect sizes, we construct the model using every fifth variant tested in each GWAS. The Pan-UK Biobank and Finngen meta-analysis reported only significant associations, so our ASH model was based on the Pan-UK Biobank data alone. Because these standard errors are generally larger than those of the meta-analysis, this strategy is conservative and will lead to a greater reduction in the effect size estimates of genome-wide significant associations. Our model was a mixture of normal distributions with an additional point mass at 0. A separate model was calculated for each GWAS, and then used to re-estimate effect sizes. *ashr* produces a posterior distribution of effect sizes for each variant. In our analysis we used the median of the distribution, but the results did not change when we tested randomly sampling from each distribution instead.

The degree of shrinkage depends on the standard error of a variant, which is influenced by frequency and local effective sample size. If ascertained variants are under selection, then we would expect that higher frequency variants have lower effect sizes on average. Prior distributions were fit to pruned variants with minor allele frequencies greater than 0.01. The posterior distribution would therefore be overestimated for high frequency variants and underestimated for low frequency variants. This should have a flattening effect on the effect size–frequency distribution and lead to conservative inferences of selection. Ultimately, because we do not make inferences about either the underlying distribution of effect sizes or the DFE, our approach must make a guess about the true effect size at each locus.

##### D.1.1 Maximum likelihood shrinkage

A third strategy uses a maximum likelihood estimate of effect sizes conditional on ascertainment based on *v*^∗^. This estimate follows from existing statistical genetics approaches to correct for ascertainment bias [54, 97, 98]. Similar to the definition of *x*^∗^, a variant is ascertained approximately when 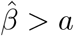, where

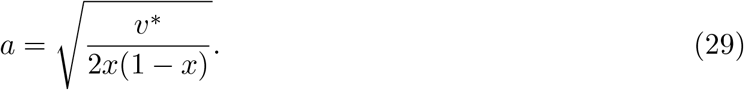

The maximum likelihood *β*, corrected for winner’s curse is therefore that which maximizes

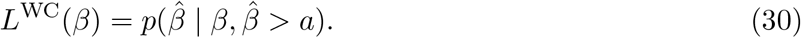

Conditioning on ascertainment gives

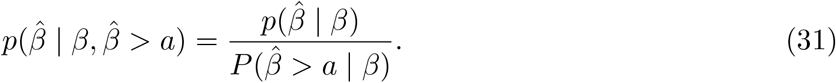

Under the normal approximation 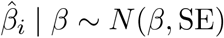,

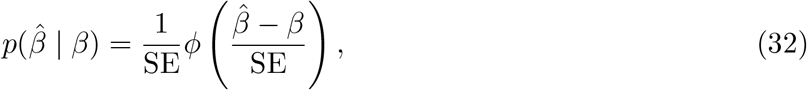

and

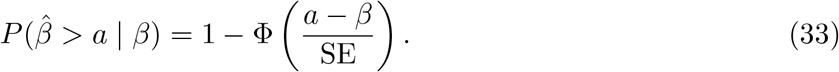

Thus

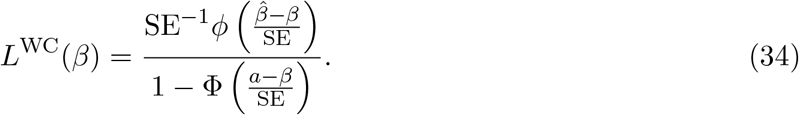

Dropping terms that do not depend on *β*, the conditional log likelihood is

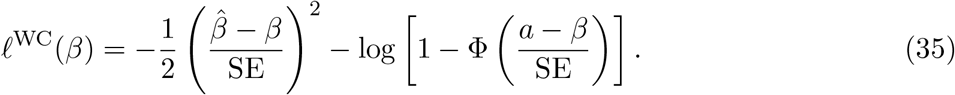

The maximum-likelihood winner’s-curse-corrected effect size is

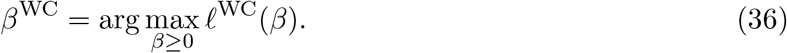

This is equivalent to finding the effect size that satisfies

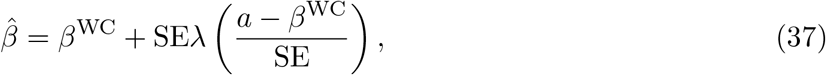

where 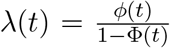 is the inverse Mills ratio. This is solved with Brent’s root finding method as implemented in scipy [91], where we know the winner’s curse corrected estimate falls in 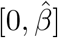. If the maximum occurs at the boundary of the parameter space, then *β*^WC^ = 0. The estimated 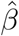 is therefore modeled as a true effect plus a winner’s curse contribution that increases with the standard error of the GWAS as well as how close the true effect size is to the ascertainment boundary.

The above strategies describe different ways to account for uncertainty in *β*. However, even if we knew the true value of *β* there would still be a distortion to the frequency distribution if the variance contribution threshold is based on 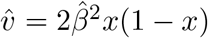. We can see this by also conditioning on this event

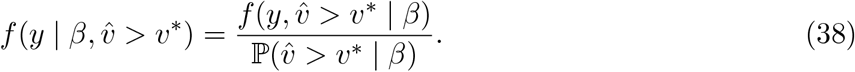

The RAF density should in theory be adjusted by the probability that the estimated effect size exceeds a threshold dependent on *y*

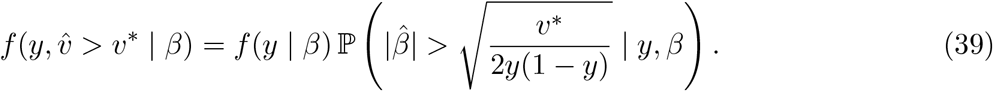

GWAS effect sizes should be approximately normally distributed conditional on the true effect size, frequency, and effective sample size.

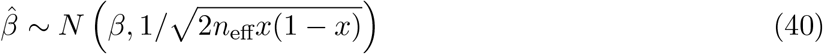

In practice, we do not make use of this adjustment and rely on simulations to evaluate the extent of the bias that results from ignoring it.

#### D.2 Empirical consequences of winner’s curse strategies

In the main text we present results treating ashr-adjusted estimates as true variant effects. The maximum-likelihood winner’s curse (MLWC) estimates shrink aggressively for variants near the ascertainment boundary, even setting *β*^WC=0^ for variants where 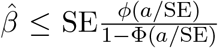. This is almost certainly an over-correction given the high rate at which genome-wide significant loci replicate in independent cohorts [99]. In contrast to results for trait simulations in section C.1, `graphld` neutral simulations did show spurious signals of selection when lead variants with their associated effect size estimates were used in inference (Figure S40). This is possibly due to the fact that these simulations used a single normal distribution of effect sizes. Without a tail of possible large effect variants, there are fewer opportunities for large effect, high frequency variants to provide evidence against selection, and a greater opportunity for winner’s curse to confound selection estimation. The application of both the posterior mean and maximum likelihood (MLWC) approaches to adjust for winner’s curse reduced the rate of these false positives (Supplements D,D.2, Figure S40). In simulations with selection, the use of lead variants also inflated the evidence for stabilizing selection models, while both shrinkage procedures reduced test statistics to the range observed when using causal variants.

However, to evaluate the robustness of our conclusions, we also re-fit all models using MLWC estimates. Despite the conservatism of this approach, doing so did not affect our conclusions with respect to the model of selection in most cases (Figure S11). For pleiotropic stabilizing models, type 2 diabetes and diverticulosis/diverticulitis, which were Bonferroni-significant in the MVP fine-mapping dataset, were only nominally supported under maximum likelihood shrinkage, as were diverticulitis, malignant neoplasms, and BBJ BMI, while BBJ CAD and IBD became unsupported. For single-trait stabilizing models, eight traits lost Bonferroni-significance (IBD, diverticulitis, malignant neoplasms, varicose veins, UKBB fine-mapping LDL, MVP diverticulosis/diverticulitis, MVP type 2 diabetes, and BBJ BMI) but remained nominal, while six became unsupported at any level (type 2 diabetes, BBJ CAD, BBJ DBP, BBJ SBP, hypothyroidism, and MVP Glaucoma).

We did not find strong systematic differences between the likelihood of stabilizing selection models in the known-causal versus lead-variant analyses, or either shrinkage procedure (Figure S41). The only notable difference was that the MLWC approach favored single-trait models. The fact this is opposite to what we saw in analyses of GWAS data may provide additional support for the pleiotropic model (Figure S11). However, `graphld` uses an alpha model of selection, making these results difficult to interpret too strongly.

The analyses of underdominance and *I*_*r*_ effect size to selection scaling were also repeated using the MLWC estimates. The evidence for the underdominant form of selection versus additive was attenuated but in no cases flipped to favor additive selection. In contrast, in *I*_*r*_ analyses estimates of *r* were consistently higher than when using posterior means (Figure S30). Using posterior mean effect sizes, we showed that a cluster of lipid traits have evidence for *r* significantly lower than the stabilizing value of 2. Using MLWC estimates, these traits had *r* estimates *>* 1, were only nominally significantly less than 2 in one instance (LDL), and all increased to ~ 2 as outliers were removed. That observation may therefore have been due to a residual effect of winner’s curse in effect size estimates. However, an overall increase of *r* values relative to 2 in the MLWC fits may indicate that estimated effect sizes were too aggressively adjusted.

### E Non-equilibrium demography

The GWAS from which we obtained summary statistics for our analyses were performed primarily in populations of European ancestry. The exception was analyses of Biobank Japan GWAS summary statistics, which were originally conducted in a population of predominantly Japanese ancestry [58]. The demographic histories shaping the genealogical relationships among these individuals and their genomes are complex and not well described by an equilibrium population model. In particular, bottlenecks and recent population growth can substantially alter the distribution of allele frequencies even in the absence of changes to the form of selection. To account for these effects, we computed likelihoods using precomputed site frequency spectrum grids under fitted demographic histories rather than relying only on equilibrium expressions.

For analyses of the original trait set, UK Biobank SuSiE-X traits, and MVP traits, we used a precomputed site frequency spectrum grid based on the demographic model of non-Finnish European history fit by Tennessen *et al*. [57]. For BBJ traits, we used an analogous grid based on the model of Japanese population history fit by Jouganous *et al*. [59] (Figure S32). These grids were generated numerically using the software `dadi` over a dense set of allele frequencies and a two-dimensional grid of additive and underdominant selection coefficients [100]. To calculate likelihoods from these grids, we obtained the site frequency spectrum corresponding to the relevant selection coefficients by linear interpolation on the precomputed grid of additive and underdominant selection coefficients. For the pleiotropic stabilizing selection model, we additionally numerically integrated over the underdominant selection grid to construct a second grid indexed by *S*_*p*_, and calculated likelihoods by linear interpolation on the resulting *S*_*p*_ grid.

We noticed that, for very strong underdominant selection, frequency spectra truncated to common MAF ranges could be nearly flat. This behavior occurred in both demographic models and was not present for the additive models of selection. While flat, the overall probability of variants falling in this frequency range was minuscule. This pathological behavior results from our conditioning on all variants being in the observable range regardless of selection coefficient. To avoid numerical contributions from extremely high-frequency selected variants that are effectively absent under these models, we truncated each precomputed spectrum in its far upper tail. Specifically, for each combination of additive and underdominant selection coefficients in the precomputed grid, we compared the remaining tail mass of the selected spectrum to that of the corresponding neutral spectrum on the stored allele-frequency grid and identified the first frequency above which the ratio of these remaining masses was at most 10^−8^. We then set the selected spectrum to zero above that frequency. Under the rough assumption that there are on the order of 1.5 × 10^7^ variants with frequency above 0.01 in humans, this corresponds to less than one selected variant expected in the truncated portion of the spectrum. This truncation has little effect on the frequency range relevant to common variation, but improves numerical stability in model fitting.

In some instances, this truncation procedure did not fully remove the impact of flattening at large selection coefficients. This led to two modes in the likelihood surfaces for *I*_2_ and *I*_*P*_, the second of which corresponded to flattening of the normalized spectra under very strong selection. To account for this, we added a final layer on top of maximum-likelihood inference that searched for a first mode at smaller parameter values even when the global maximum occurred at the higher mode. Among the traits considered in the main analysis, this changed the reported fit only for schizophrenia under pleiotropic stabilizing selection (Figure S50).

### F Variant matching across studies

Comparisons of effect sizes across GWAS require identifying the same variant in both studies. We treated this as a sensitivity analysis rather than as a full trans-ancestry replication analysis. For each trait, we started from the loci retained in the original model fitting pipeline after LD filtering and after applying the variance-contribution threshold. These loci define the denominator for matching counts. We then intersected this set with the corresponding processed summary statistics from the second study by SNP identifier. No proxy variants were substituted for unmatched loci.

This exact-matching strategy is conservative because the same association may be tagged by a different variant in another study or ancestry. It therefore separates two possible effects on inference: loss of loci caused by the requirement that the original variant be present in the second study, and changes in model fit caused by replacing effect-size estimates at the loci that remain. To isolate the first effect, we fit a matched-original analysis in which only exactly matched loci were retained, but the original effect-size estimates were used. To isolate the second effect, we used the same matched set of loci but replaced original effect-size estimates with those from the second study.

### G BBJ variant matching

Data from Biobank Japan (BBJ) provide an opportunity to control for winner’s curse by using independently estimated summary statistics. This analysis requires a set of loci that can be compared across studies. To select these loci, we took traits from our analysis that had also been measured in well-powered BBJ GWAS (BMI, CAD, DBP, HDL, height, LDL, RBC, SBP, SCZ, T2D, triglycerides, and WBC). We used variants that were present in both studies. Variants were first filtered by applying the genome-wide significance threshold and variance-contribution threshold to the original GWAS. Variants were then filtered for nominal significance (*P <* 0.05) in the BBJ GWAS. This nominal significance filter was selected to balance three goals: 1) to allow for the lower power of some BBJ GWAS, 2) to avoid winner’s curse in our use of BBJ data, and 3) to drop loci that are not trait-associated in BBJ, possibly because the causative variant is absent or rare in the BBJ population.

after filtering variants, we followed our previous clumping procedure, prioritizing variants based on their P-values in the original GWAS. After this ascertainment procedure—based primarily on our original summary statistics—we used BBJ-estimated effect sizes to test our model likelihoods.

Because BBJ effect estimates could be noisy for many of the original loci, due to differences in sample size and variant frequencies between studies, we tested two strategies for effect-size replacement: replacement for all matched loci, or only for loci expected to be sufficiently powered in BBJ. To approximate the expected power in BBJ we calculated

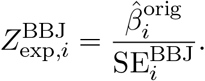

where 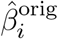 is the original effect-size estimate and 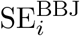 is the BBJ standard error. This quantity reflects the Z-score we might expect from the BBJ GWAS if the original effects were estimated without error. It will therefore be inflated by winner’s curse, but we find it useful as an approximate metric. In effect replacement analyses we replaced all matched variants and then used thresholds of *Z*_exp_ ≥ 2 and *Z*_exp_ ≥ 5 (Figure S17). Comparisons between matched-original and BBJ-replacement fits therefore test the effect of replacing original effect estimates among loci that could be exactly matched, while comparisons between full-original and matched-original fits show the effect of exact matching itself.

For traits we were able to analyze with effect-matching, the likelihoods of stabilizing models were attenuated primarily due to the drop-out of variants for which we could not match to BBJ in the first place. This is due to differences in sample size and because these variants tended to be rare (Figure S18). Effect size matching did not have a strong or consistent effect on the evidence for stabilizing selection on top of dropping these loci (Figure S19). Effect size replacement also failed to have a strong effect on the likelihood of the pleiotropic versus one-dimensional stabilizing models (Figure S20). While replacement effects were noisy, these result support the notion that winner’s curse did not substantially create signatures of selection or preferences for pleiotropic models.

## H The distribution of variance contributions

Simons *et al*. [37] derived the distribution of variance contributions exceeding the threshold *v*^∗^ under the single-trait and pleiotropic models of stabilizing selection. These results apply to the case when selection is strong and the population is at equilibrium. They argued that, because strongly selected variants should be discovered first by GWAS, the likelihoods of the observed genome-wide significant *v* values under the two models could provide a test for whether selection is highly pleiotropic or not. We implemented this test and applied it to simulated GWAS data from both trait- and DFE-based simulations as well as to genome-wide significant variants from all traits. We also derived the limiting form of the tail distribution for both models as the variance contribution threshold becomes large relative to the average variance contribution of a strongly selected site.

Following Simons *et al*. [37], the distribution of variance contributions for strongly selected sites under the single-trait stabilizing selection model is

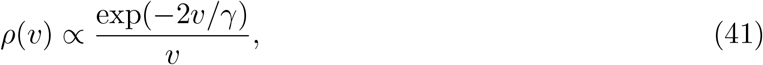

where *γ* is the expected variance contribution of a site under strong selection. Because only variants with variance contribution exceeding *v*^∗^ are ascertained, it makes sense to work with the distribution of 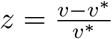 so that

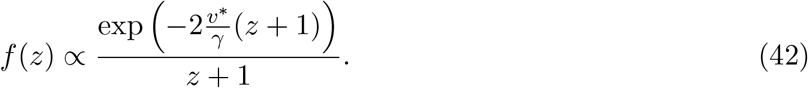

Also define 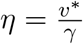:

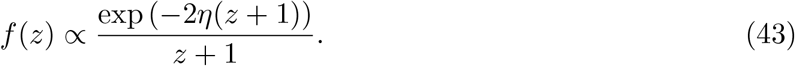

Applying the same approach to the pleiotropic variance distribution

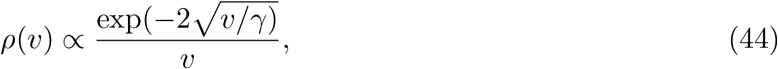

we get

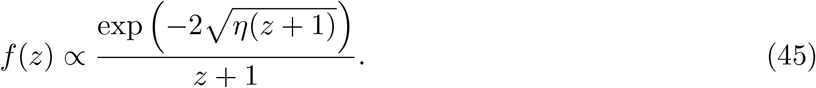

### H.1 Simulation results

The inclusion of variants under weaker selection or at higher frequencies violates the assumptions leading to the distributions in equations (43) and (45). The assumption of a constant population size is also not consistent with allele frequencies in humans. We applied the Simons *et al*. (2018) test to simulated data from trait- and DFE-based simulations in order to determine its robustness to these violations. To focus only on the effects of changes to the population genetics model, we only used simulations free of ascertainment bias. For each simulation, we calculated the maximum likelihood value of *η* for both the one- and high-dimensional distributions and compared likelihoods.

At the neutral end of trait-based simulations using the IBD-derived distribution of effect sizes, the high-dimensional pleiotropy model always had a significantly higher likelihood (Figure S43). For strong selection in one-dimensional simulations, the log-likelihood difference between the models decreased towards zero. In high-dimensional simulations the evidence for the high-dimensional model also decreased as selection became strong. In DFE-based simulations a comparable bias towards the high-dimensional model was also observed (Figure S44). Overall, the inclusion of high-frequency variants strongly favored the model of high-dimensional stabilizing selection. These appear in our simulation data due to a combination of including alleles under weak/moderate selection and the out-of-Africa bottleneck allowing some strongly selected variants to drift to high frequencies. That the inclusion of high-frequency variants would favor the high-dimensional model makes intuitive sense, as this model allows the possibility of occasional large effect variants experiencing weaker selection when their effects on pleiotropic traits are small by chance.

### H.2 Application to trait data

We observed similar results when applying the Simons *et al*. [37] model to estimated variance contributions from our set of traits using the *v*^∗^ corresponding to p–value = 5 × 10^−8^ as well as the p–value cutoff itself. The high-dimensional model had a substantially higher likelihood for all traits, even those like breast cancer and IBD where we detected no significant deviation from neutrality (Figure S42). For some traits, like schizophrenia and systolic/diastolic blood pressure, the variance contribution distribution fit the data reasonably well, but for most traits we observed a fatter tail of large variance contributions than in the best-fit one- and high dimensional models.

### H.3 Limiting distributions

We derived limiting forms for the distributions of variance contributions as the variance contribution threshold becomes large relative to the average variance contribution of a strongly selected site. That is, as *η* becomes large. GWAS will likely only uncover the tail of strongly selected mutations affecting a given trait, so it makes sense to ask how the distributions corresponding to the two models of pleiotropy differ in their tail behavior.

Starting with the one-dimensional model, calculating the normalizing constant for equation (43) yields an incomplete gamma function Γ(0, 2*η*). When *η* is large, this is approximately

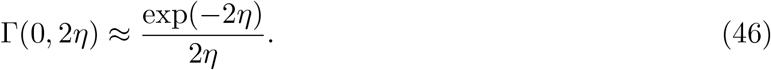

The density of *z* can then be approximated as

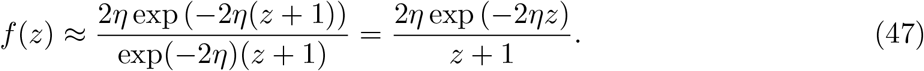

If *η* is large enough, the density will be concentrated at low values of *z* as larger variance contributions will be increasingly suppressed by selection. We can ignore *z* in the denominator so that

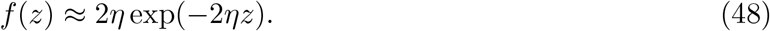

This is an exponential distribution with rate 2*η*.

For the high-dimensional model, the normalizing constant is also an incomplete gamma function, 2Γ(0, 2 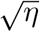). We can apply the same approximation as in equation (46), though in this case it must be that 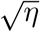 is large. Doing this for equation (45) gives

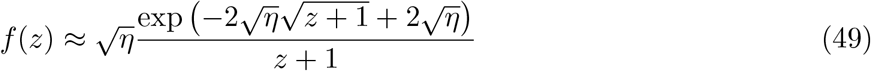

Assuming that the density is concentrated at small *z*, we can use 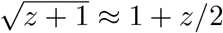, giving

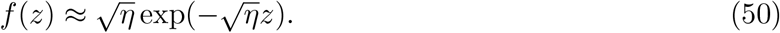

This is again an exponential distribution, although the rate is proportional to 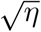 in this case. The two models are not distinguishable when *v*^∗^ is sufficiently high because both models have exponential tails and *γ* is unknown. The one-dimensional model will approach the exponential limit faster than the high-dimensional model. This implies an asymmetry in the ability to distinguish the two where it is more likely for traits under one-dimensional stabilizing selection to look identical to a high-dimensional model with a lower *γ*.

## I Effect-size to selection scaling

To test the prediction, made by models of stabilizing selection, that the strength of selection scales with *β*^2^, we fit modified models where this scaling was allowed to vary. For single-trait stabilizing selection, the *β*^2^ scaling appears directly in the relation *s* = *I*_2_*β*^2^. In the pleiotropic model, the same scaling appears through the conditional relationship E[*β*^2^ | *s*] ∝ *s*. We therefore introduced a free scaling exponent *r* and fit models in which the strength of selection scales with |*β*|^*r*^.

For single-trait stabilizing selection, this gives

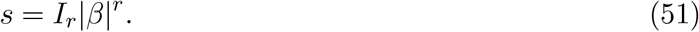

For pleiotropic stabilizing selection, the corresponding modification is less direct. We used

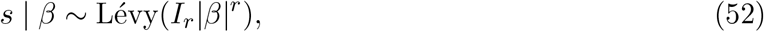

which provides a simple test of departures from the stabilizing expectation because the median selection coefficient remains proportional to *I*_*r*_|*β*|^*r*^, even though the original Lévy distribution is motivated by the quadratic relationship *β* | *s* ~ *N* (0, *cs*).

We applied these models to the original trait set using the same processing steps and using posterior mean shrunk GWAS effect-size estimates as stand-ins for *β* and also tested robustness to the maximum likelihood winner’s curse adjustment. For each trait we fit both the stabilizing models and their *r*-scaling extensions by maximum likelihood, using the same numerical optimization procedure described above. In the single-trait model we optimized log_10_(*I*_*r*_) and *r*, initialized at the maximum-likelihood estimate of *I*_2_ and at *r* = 2, with bounds log_10_(*I*_*r*_) ∈ [−8, 2] and *r* ∈ [0, 10]. In the pleiotropic model we optimized the analogous *I*_*r*_ parameter and *r*, initialized at the maximum-likelihood estimate of *I*_*p*_ and at *r* = 2. The upper bound for the pleiotropic intensity was set by the largest value represented in the precomputed *S*_*p*_ grid, as in the standard pleiotropic fit. The standard *I*_2_ and *I*_*p*_ models were re-fit on the same variants, allowing likelihoods from the free-scaling and *r* = 2 models to be compared directly.

### I.1 Outlier detection

To assess whether estimates of the scaling exponent were driven by a small number of influential loci, we performed a case-deletion analysis within each trait [101]. For the primary analysis, influence was measured under the pleiotropic stabilizing selection model. We refit the pleiotropic stabilizing selection model after removing each variant in turn. Let 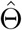 denote the maximum-likelihood estimate from the full set of variants and 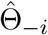 the estimate obtained after removing variant *i*. We assigned variant *i* a case-deletion deviation

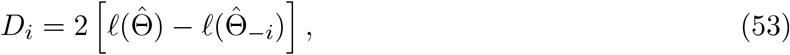

where both likelihoods were evaluated on the full dataset. Variants with larger values of *D*_*i*_ were those for which deleting the variant led to parameter estimates that fit the full data substantially worse, and were therefore treated as more influential.

We then ranked variants within each trait by this deviation and repeated the standard and *r*-scaling fits after removing the top 0, 1, 2, or 5 variants. When variants were removed, ties in case-deletion deviation were broken by the estimated variance contribution. As a sensitivity analysis, we also repeated the outlier-ranking procedure using a model with the scaling exponent fixed at *r* = 0, and then performed the same refits after removing the top-ranked variants from that alternative ranking.

### I.2 Underdominance

To test an additional prediction of stabilizing selection, we also compared the “underdominant” form of the frequency spectrum that arises under stabilizing selection to the standard additive form. We fit models of the effect-size to selection relationships in equations (14) and (17) that were preserved, but variants instead experienced additive negative selection. This preserves the approximate behavior of the models at low allele frequencies while removing the frequency dependence that weakens and reverses selection around intermediate frequencies under the underdominant model (Supplementary Figure S45). We fit these additive models to the original trait set using the same procedures as described above and compared to the underdominant form using AIC.

## J Code and data availability

Processed GWAS data and code for analyses, simulations, and figures can be accessed at https://github.com/emkoch/smilenfer.

## K Figures

**Figure S1:**
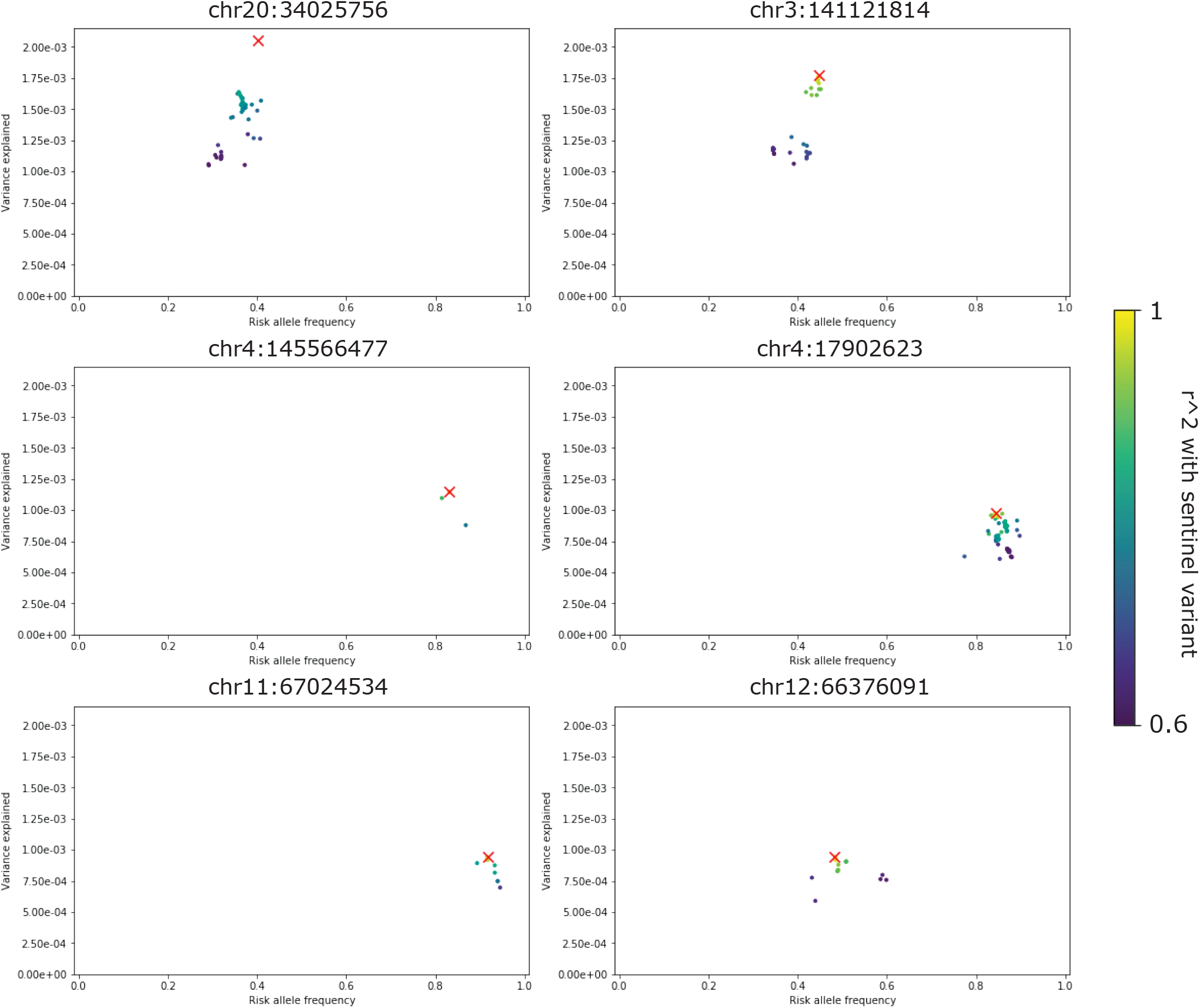
Sentinel variants cluster tightly with linked variants in high LD. A sample of six genome-wide significant loci for standing height. The sentinel variant (lowest *p*-value) is shown alongside nearby variants in high LD. Only variants with similar trait-increasing allele frequencies have a high *r*^2^ with the sentinel variant.

**Figure S2:**
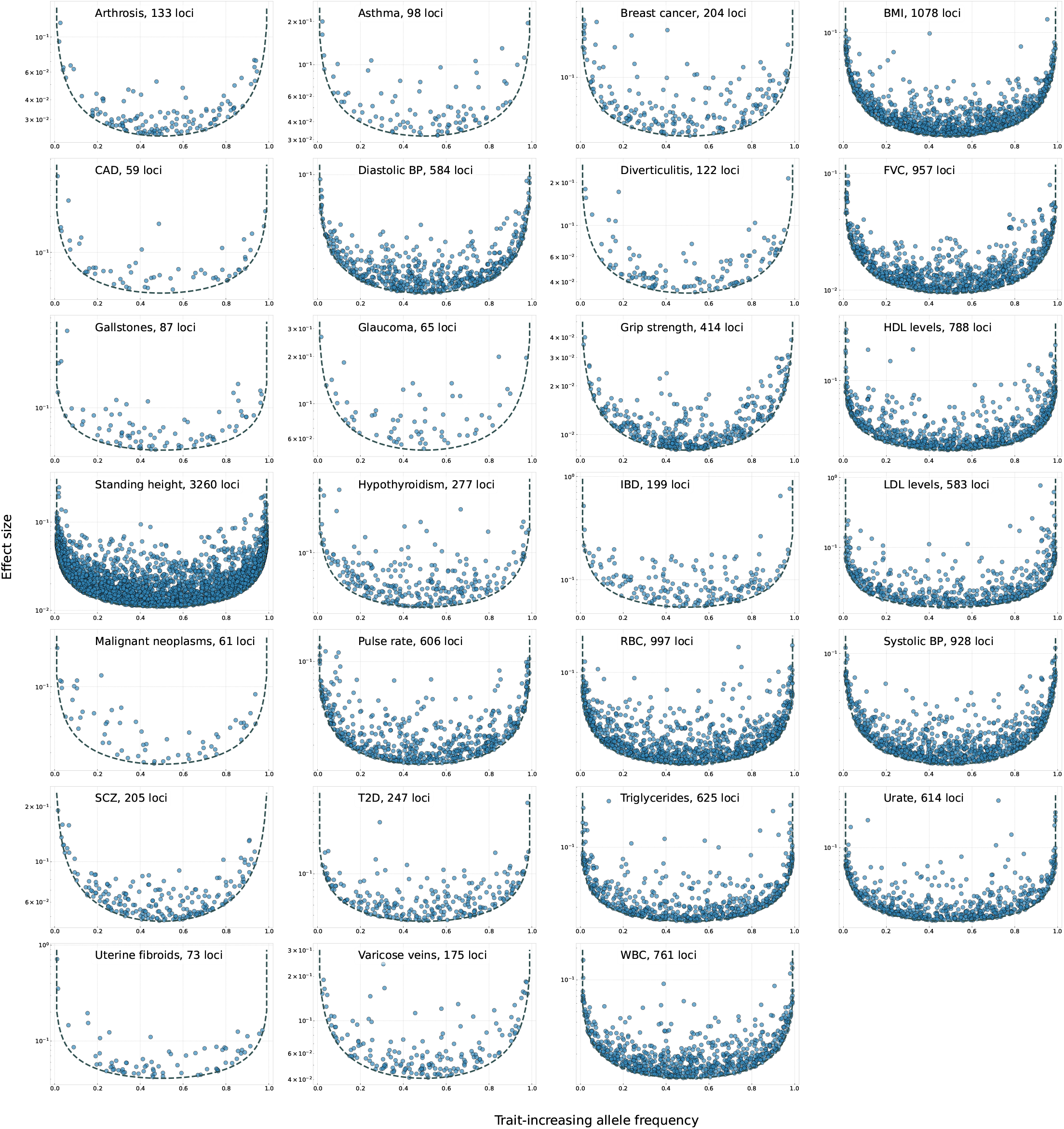
U-shaped distributions of trait or disease risk-increasing allele frequencies and effect sizes for all analyzed traits. For disease traits, the effect size is calculated as the odds-ratio minus one. Raw GWAS estimates are shown here without shrinkage. We include variants that are both genome-wide significant and have an estimated variance contribution exceeding a threshold variance associated with 5 × 10^−8^. The dashed line shows the effect size cutoff imposed by the variance contribution threshold.

**Figure S3:**
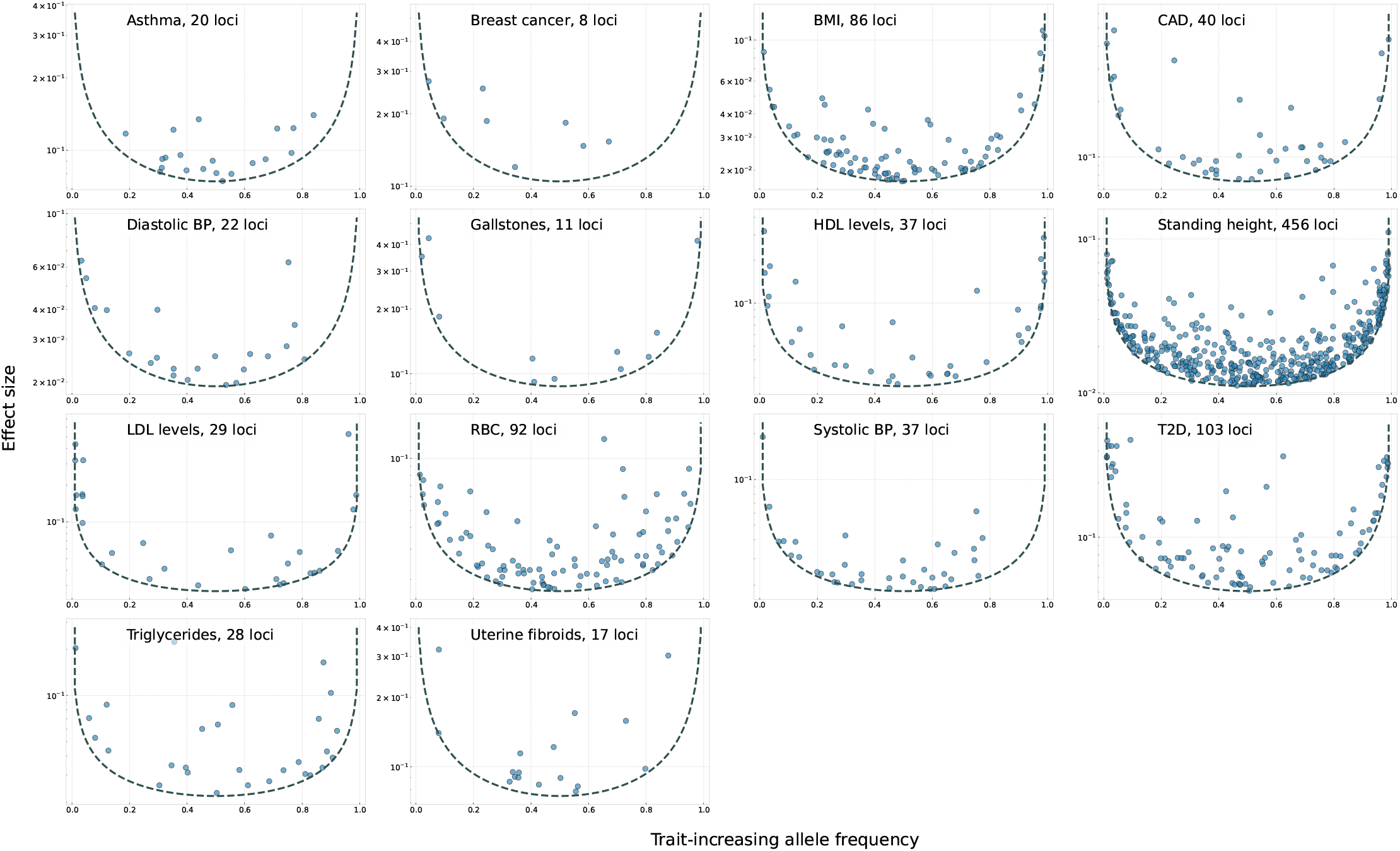
U-shaped distributions of trait or disease risk-increasing allele frequencies and effect sizes for BBJ traits after high-effect clumping. For disease traits, the effect size is calculated as the odds-ratio minus one. Raw GWAS estimates are shown here without shrinkage. Starting from the processed BBJ trait tables, loci were additionally clumped using a 500 kb procedure that retained the variant with the largest effect size in each region, matching the version of the data used for the main BBJ model fits. We include variants that are both genome-wide significant and have an estimated variance contribution exceeding a threshold variance associated with 5 × 10^−8^. The dashed line shows the effect size cutoff imposed by the variance contribution threshold. Panel labels also indicate the number of retained loci for each trait.

**Figure S4:**
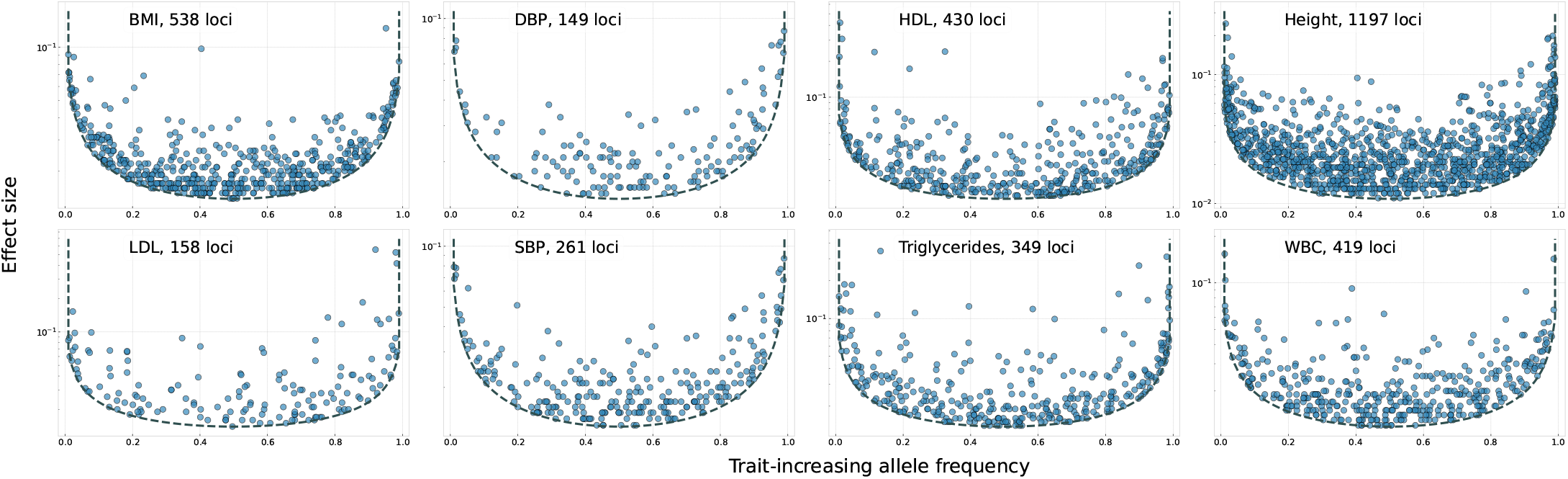
U-shaped distributions of trait-increasing allele frequencies and effect sizes for UKBB SuSiE-X traits. Raw effect size estimates are shown here without shrinkage. For each trait, one variant was sampled from each fine-mapped credible set, and we include sampled variants whose estimated variance contribution exceeds the threshold associated with 5 × 10^−8^. The dashed line shows the effect size cutoff imposed by the variance contribution threshold. Panel labels also indicate the number of retained loci for each trait.

**Figure S5:**
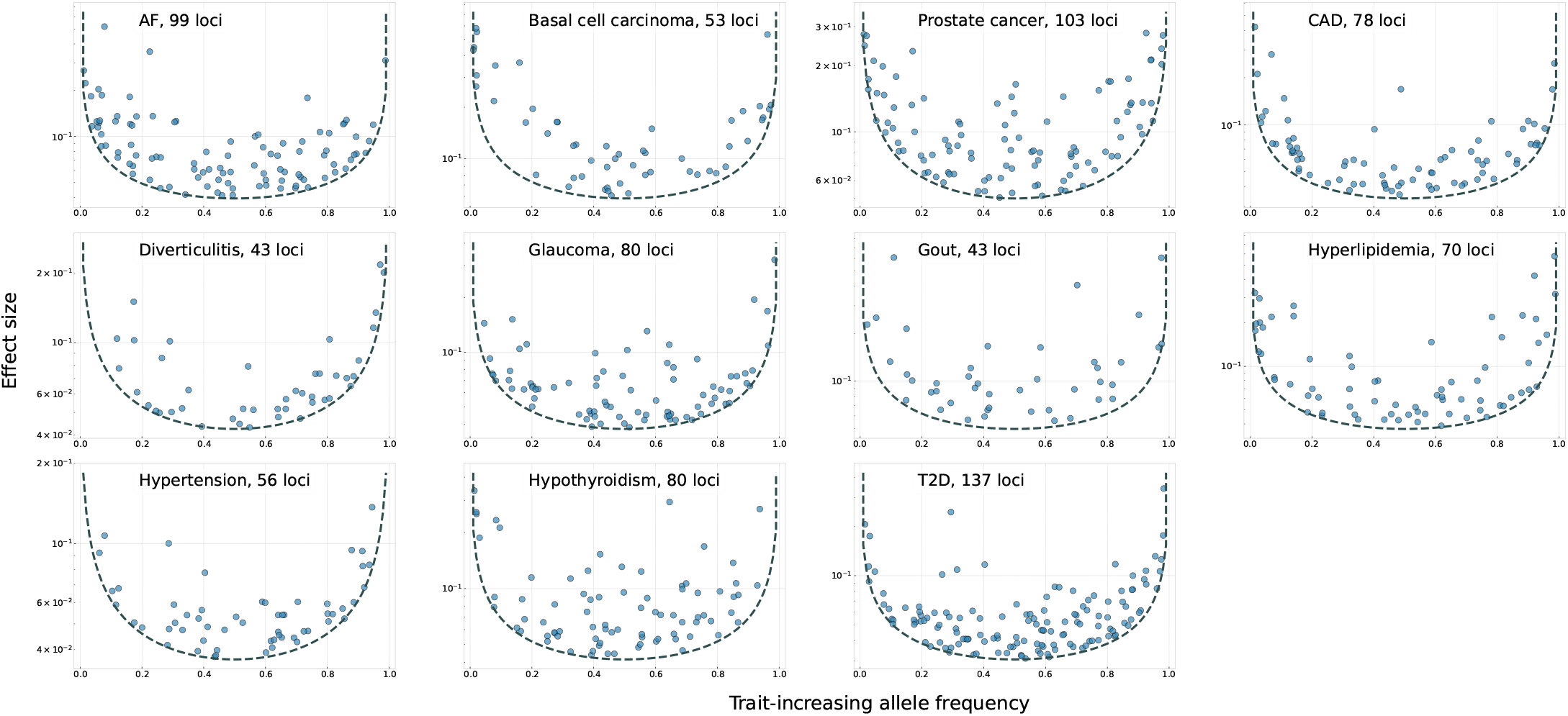
U-shaped distributions of trait or disease risk-increasing allele frequencies and effect sizes for MVP traits. For disease traits, the effect size is calculated as the odds-ratio minus one. Raw effect size estimates are shown here without shrinkage. For each trait, one variant was sampled from each fine-mapped credible set, and we include sampled variants whose estimated variance contribution exceeds the threshold associated with 4.6 × 10^−11^. The dashed line shows the effect size cutoff imposed by the variance contribution threshold. Panel labels also indicate the number of retained loci for each trait.

**Figure S6:**
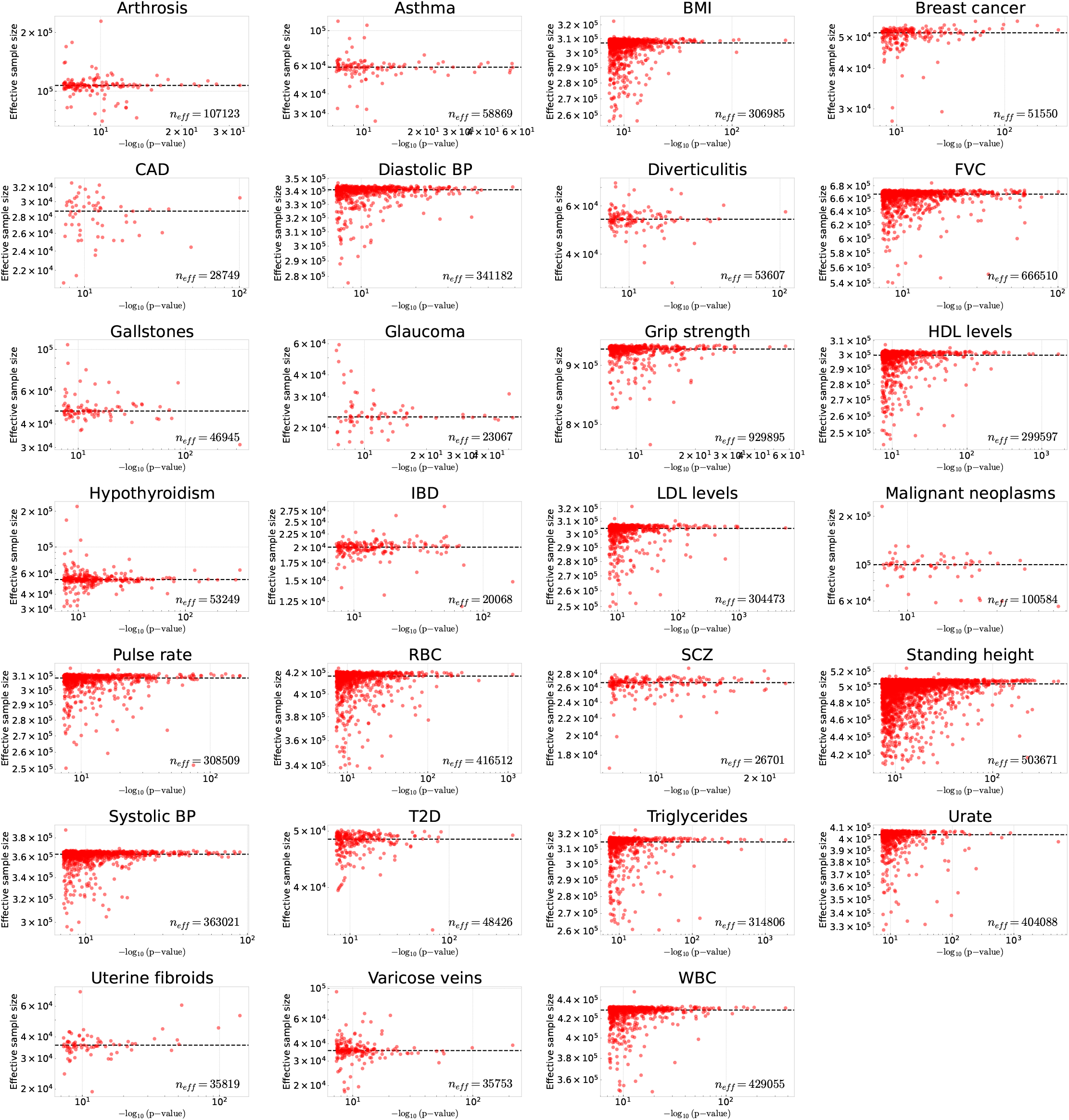
Effective sample size and *p*-value relationship for all traits. For each trait under consideration we plot the relationship between effective sample sizes calculated using GWAS effect sizes 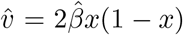 and *p*-values 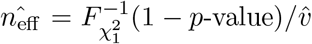. Only genome-wide significant variants (*p*-value*<* 5 × 10^−8^) are shown. Dashed lines and values in the bottom right correspond to the median effective sample size among genome-wide significant variants.

**Figure S7:**
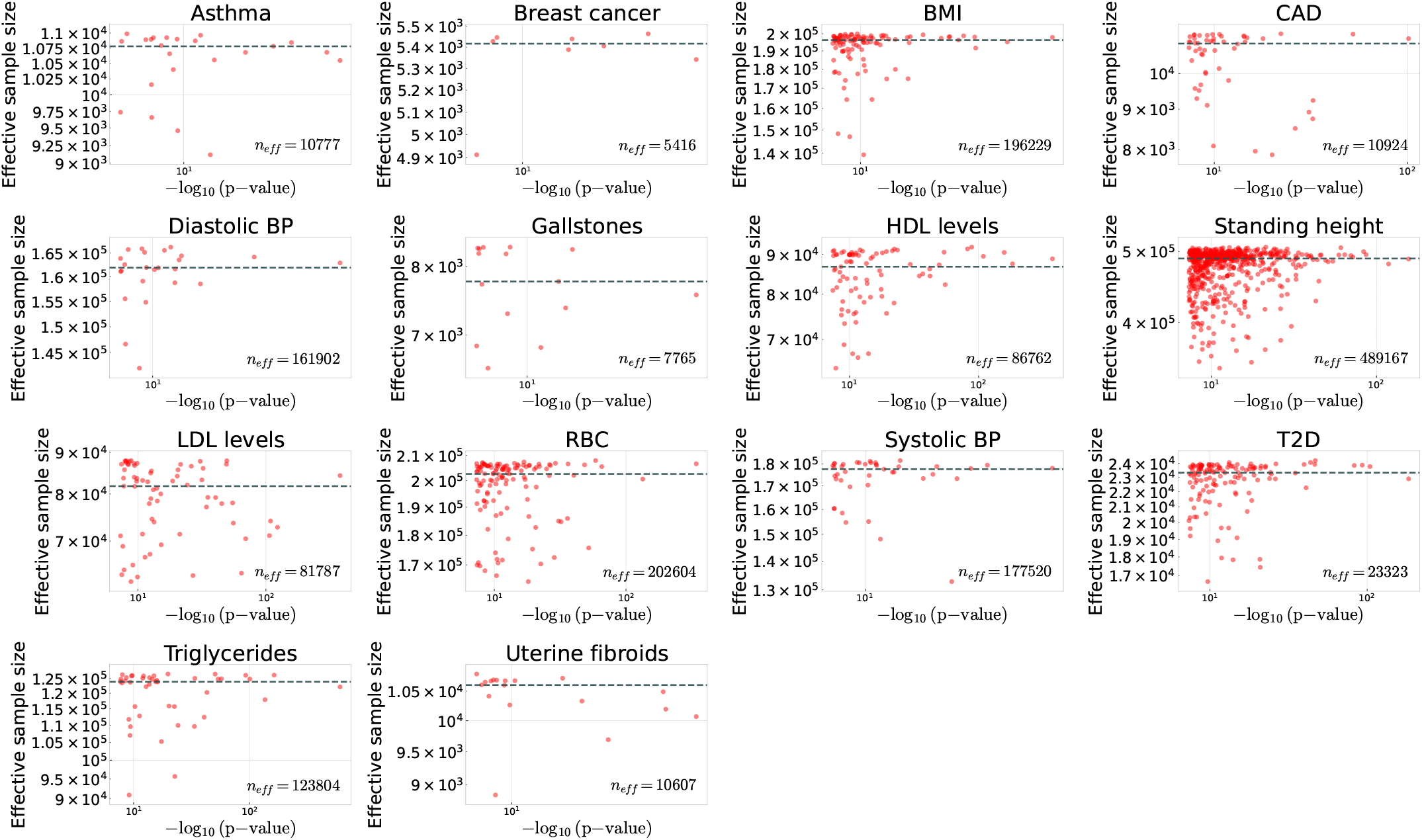
Effective sample size and *p*-value relationship for BBJ traits. For each BBJ trait under consideration we plot the relationship between effective sample sizes calculated using GWAS effect sizes 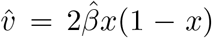 and *p*-values 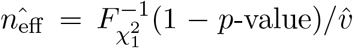. Only genome-wide significant variants (*p*-value*<* 5 × 10^−8^) are shown. Dashed lines and values in the bottom right correspond to the median effective sample size among genome-wide significant variants.

**Figure S8:**
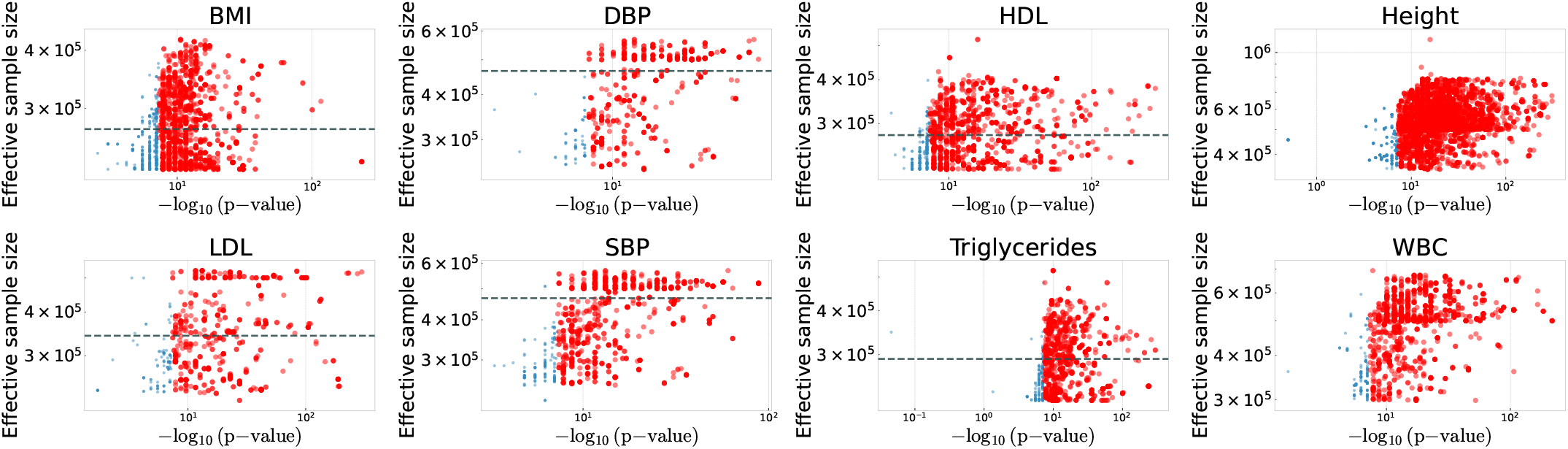
Effective sample size and *p*-value relationship for UKBB SuSiE-X traits. For each trait under consideration we plot the relationship between effective sample sizes calculated using fine-mapped effect sizes 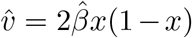 and *p*-values 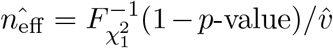. Only variants corresponding to genome-wide significant GWAS loci (*p*-value*<* 5 × 10^−8^) are shown. Dashed lines and values in the bottom right correspond to the median effective sample size among genome-wide significant variants.

**Figure S9:**
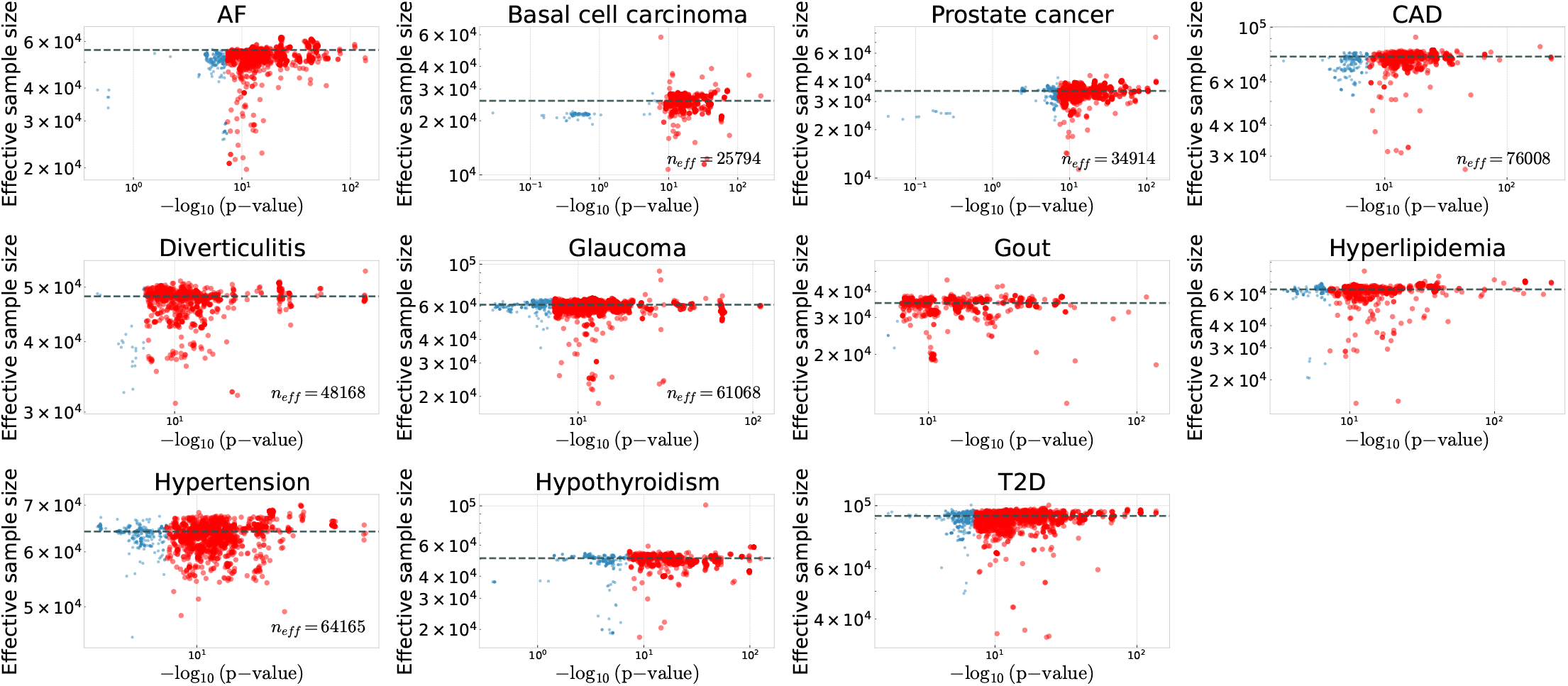
Effective sample size and *p*-value relationship for MVP fine-mapped traits. For each trait under consideration we plot the relationship between effective sample sizes calculated using fine-mapped effect sizes 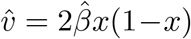 and *p*-values 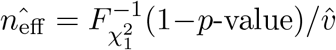. Only phenome-wide significant variants (*p*-value*<* 4.6 × 10^−11^) are shown. Dashed lines and values in the bottom right correspond to the median effective sample size among phenome-wide significant variants.

**Figure S10:**
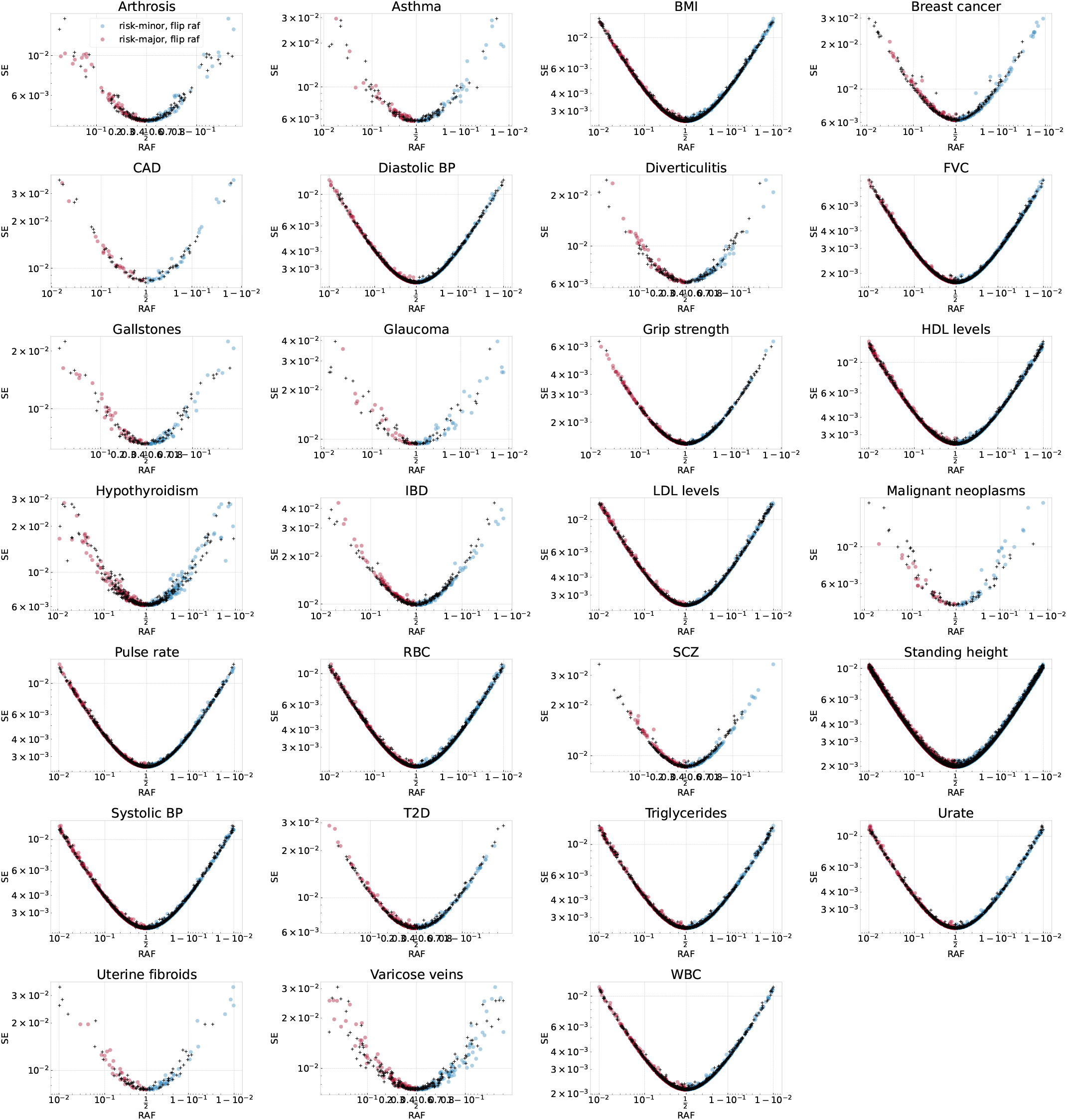
Relationship between standard errors and trait or disease risk-increasing allele frequencies. Black points show standard errors obtained from GWAS summary statistics plotted as a function of the trait or risk increasing allele frequency (RAF). To visualize asymmetries in between variants where the minor or major allele is correlated with increased risk, we also flip the plot along the RAF= 0.5 line (blue and red points). Equation (40) predicts that standard errors should depend only on minor allele frequencies and the effective sample size of the study. In practice, noise results from correlations between the variants and other genotypes and covariates. Asymmetry is also apparent for disease traits where there is greater power to detect low-frequency risk alleles than low-frequency protective alleles. RAFs are plotted on a logit scale.

**Figure S11:**
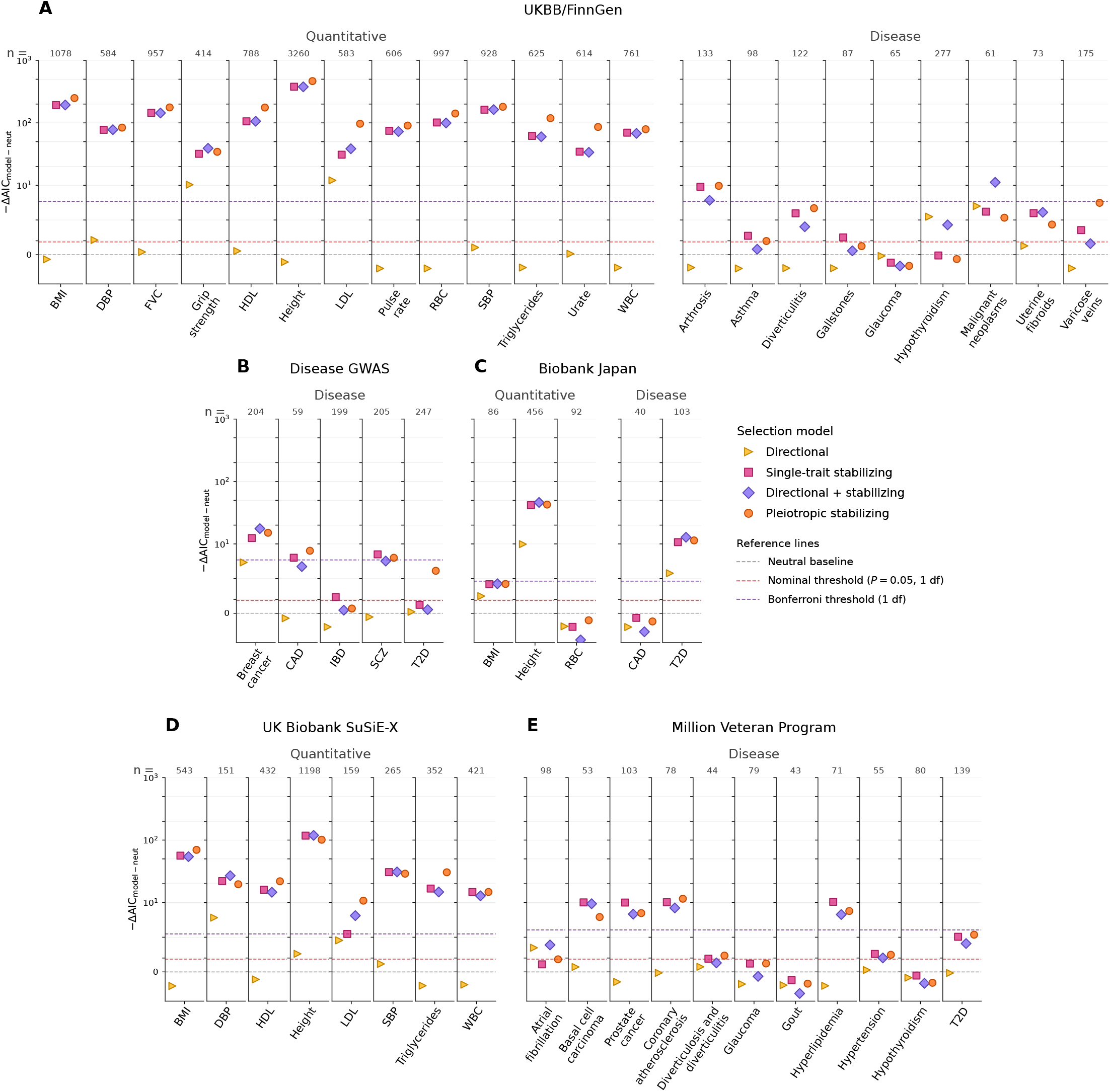
Model comparison favors stabilizing selection across categories of traits when using maximum likelihood shrinkage of effect sizes. We repeated the Figure 3 analysis using variant effects adjusted with the maximum likelihood winner’s curse procedure instead of posterior means. BBJ analyses only clumped by highest *p*-value and analyses of fine-mapped datasets only used single samples from credible set probabilities.

**Figure S12:**
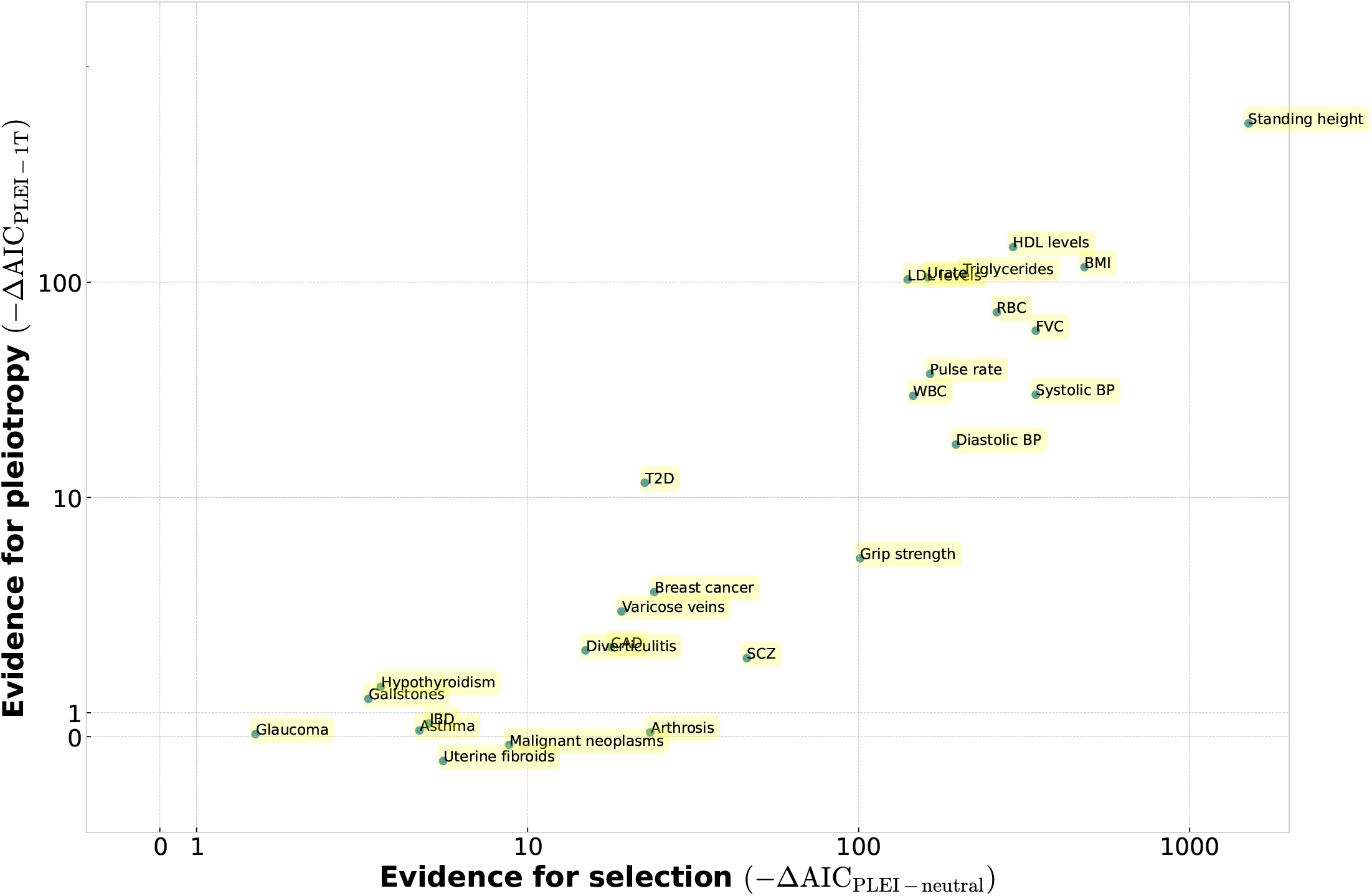
Likelihood increases for pleiotropic versus single-trait stabilizing selection. Evidence for selection was defined as the likelihood increase of pleiotropic stabilizing selection versus neutrality, while evidence for pleiotropy was defined as the likelihood increase of pleiotropic versus single-trait stabilizing selection. Log-likelihoods were put on the AIC scale to facilitate comparison. Similarly to simulations, significant likelihood differences between models appear for −ΔAIC *>* 10 (Figure S38).

**Figure S13:**
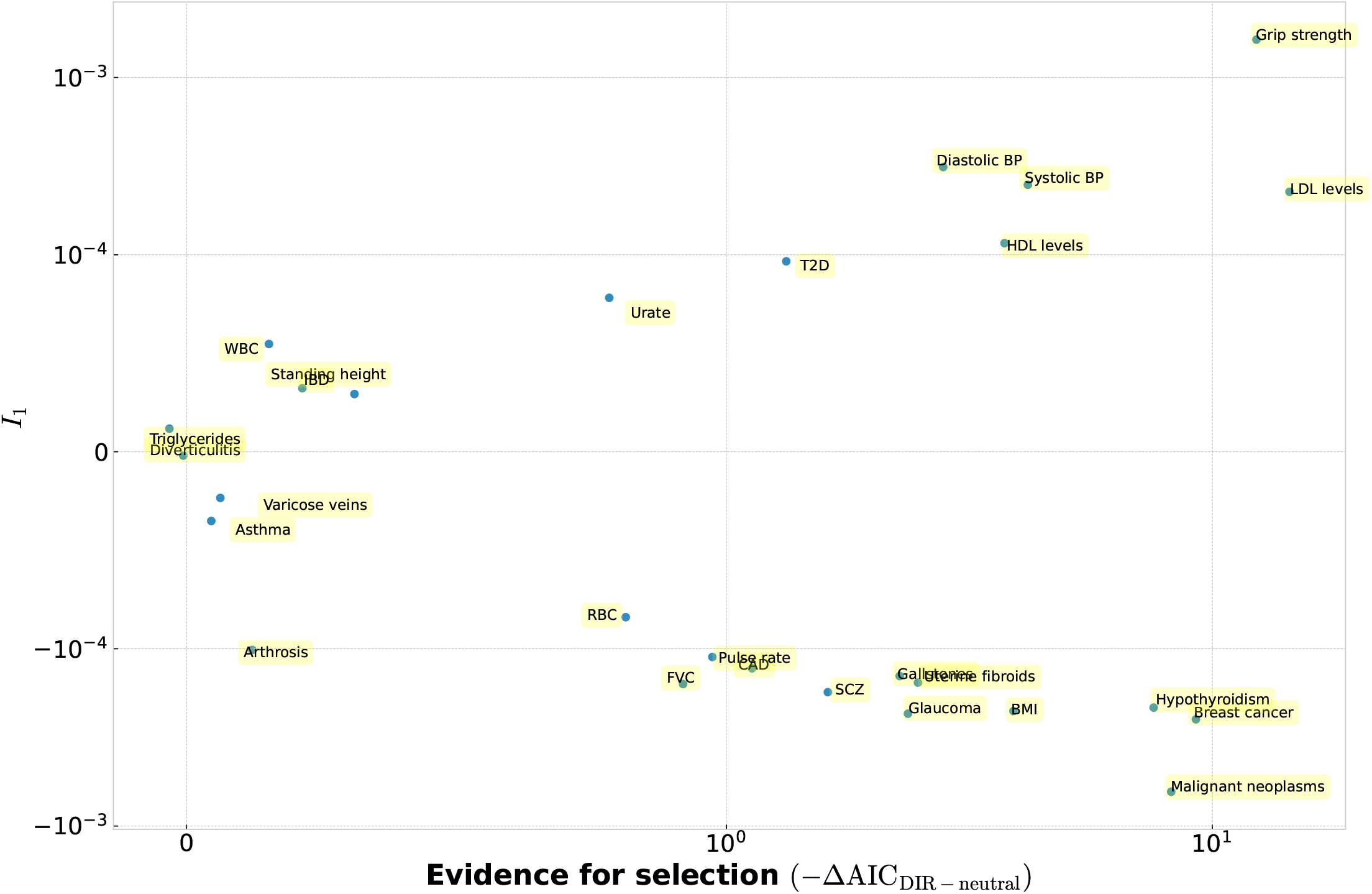
Directional selection estimates indicate selection against risk alleles in disease traits and selection against trait-decreasing alleles for quantitative traits. The y-axis displays estimated directional selection parameters for all 27 traits. Negative *I*_1_ values indicate negative selection against trait-increasing or risk alleles. The x-axis displays the improvement in likelihood versus the neutral model obtained by fitting the directional selection model for each trait.

**Figure S14:**
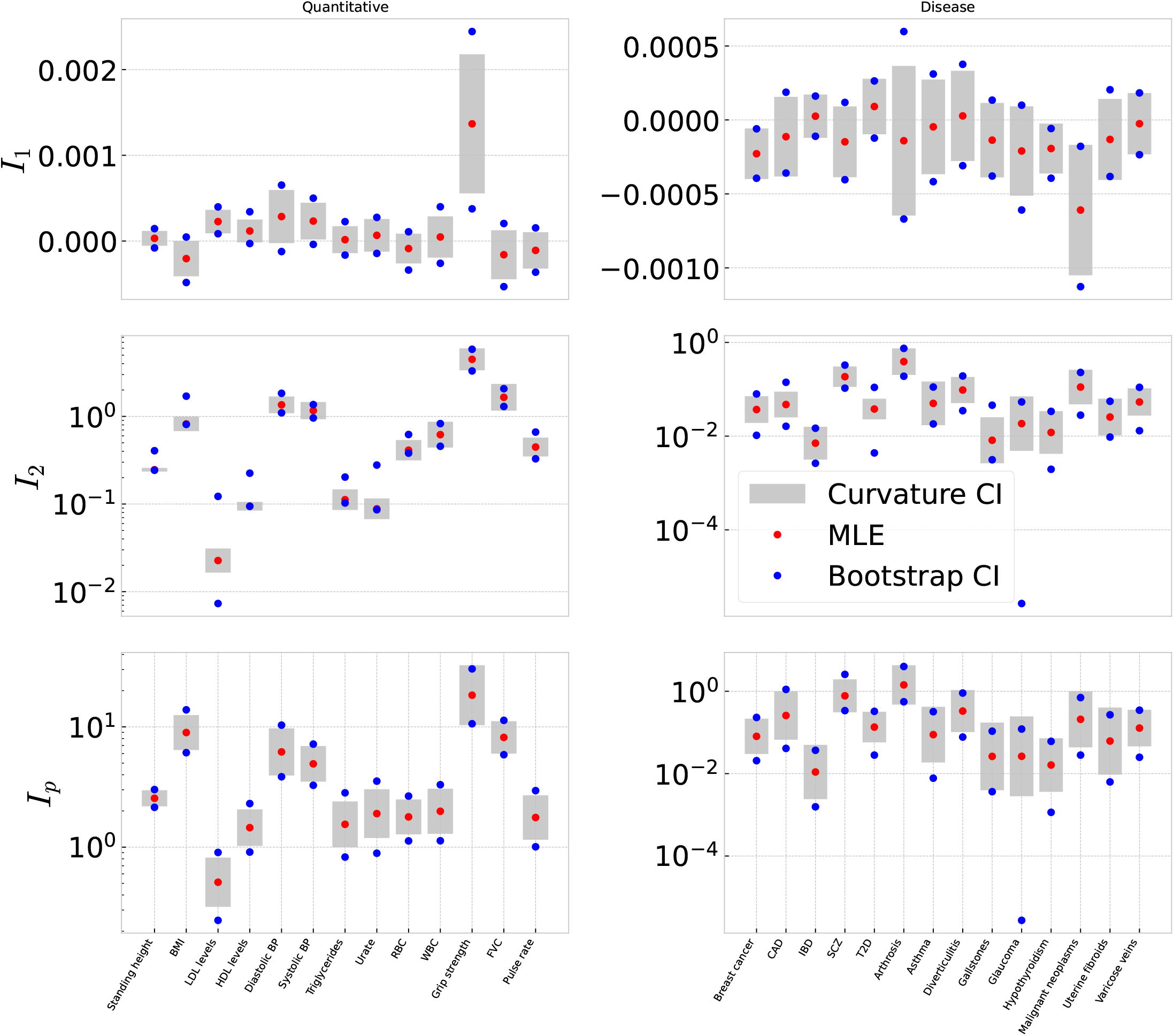
Confidence intervals for selection parameter estimates in the original trait fits. For each original trait, red points show maximum-likelihood estimates of the directional selection parameter *I*_1_, the single-trait stabilizing selection parameter *I*_2_, and the pleiotropic stabilizing selection parameter *I*_*p*_. Gray intervals show 95% Wald confidence intervals calculated from the curvature of the log-likelihood surface around the maximum-likelihood estimate. To calculate this curvature, we evaluated the likelihood across a local grid of log_10_(*I*_2_) or log_10_(*I*_*p*_) values, smoothed the log-likelihood values, and used the inverse Fisher information to compute the standard error. Blue points show bootstrap confidence intervals calculated as the 2.5th and 97.5th percentiles across 1,000 bootstrap datasets generated by resampling retained loci with replacement and re-fitting the selection model. Quantitative traits and disease traits are shown separately. Fits used raw GWAS effect-size estimates.

**Figure S15:**
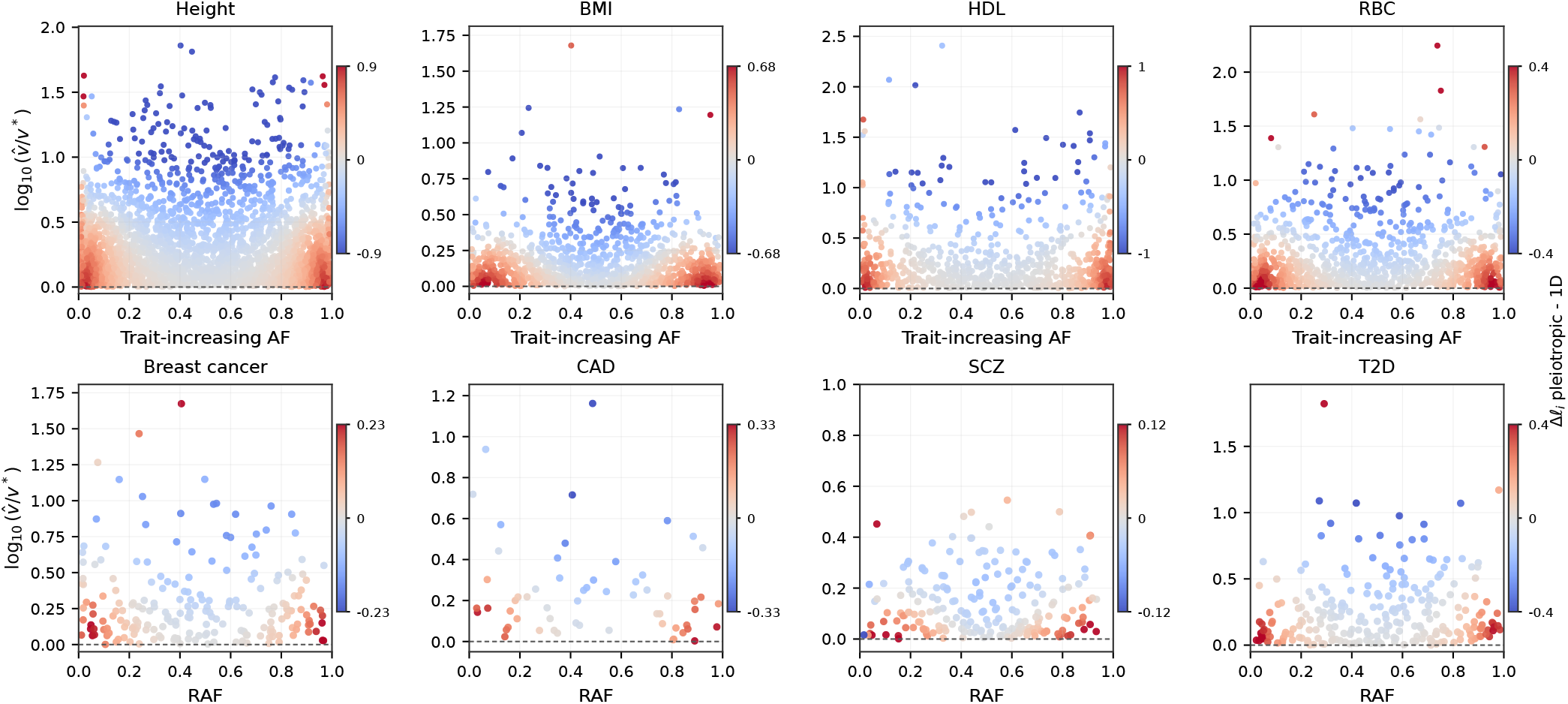
Locus-level likelihood contributions for pleiotropic versus single-trait stabilizing selection. For each displayed trait, points show retained loci from the original analysis. The x-axis shows the trait-increasing allele frequency for quantitative traits and the risk allele frequency (RAF) for disease traits. The y-axis shows the variance contribution of each locus relative to the ascertainment threshold, 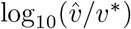. Point color shows the difference in per-locus log-likelihood contribution between the fitted pleiotropic and single-trait stabilizing models, Δ*ℓ*_*i*_ = *ℓ*_*i*,plei_ − *ℓ*_*i*,1D_.

**Figure S16:**
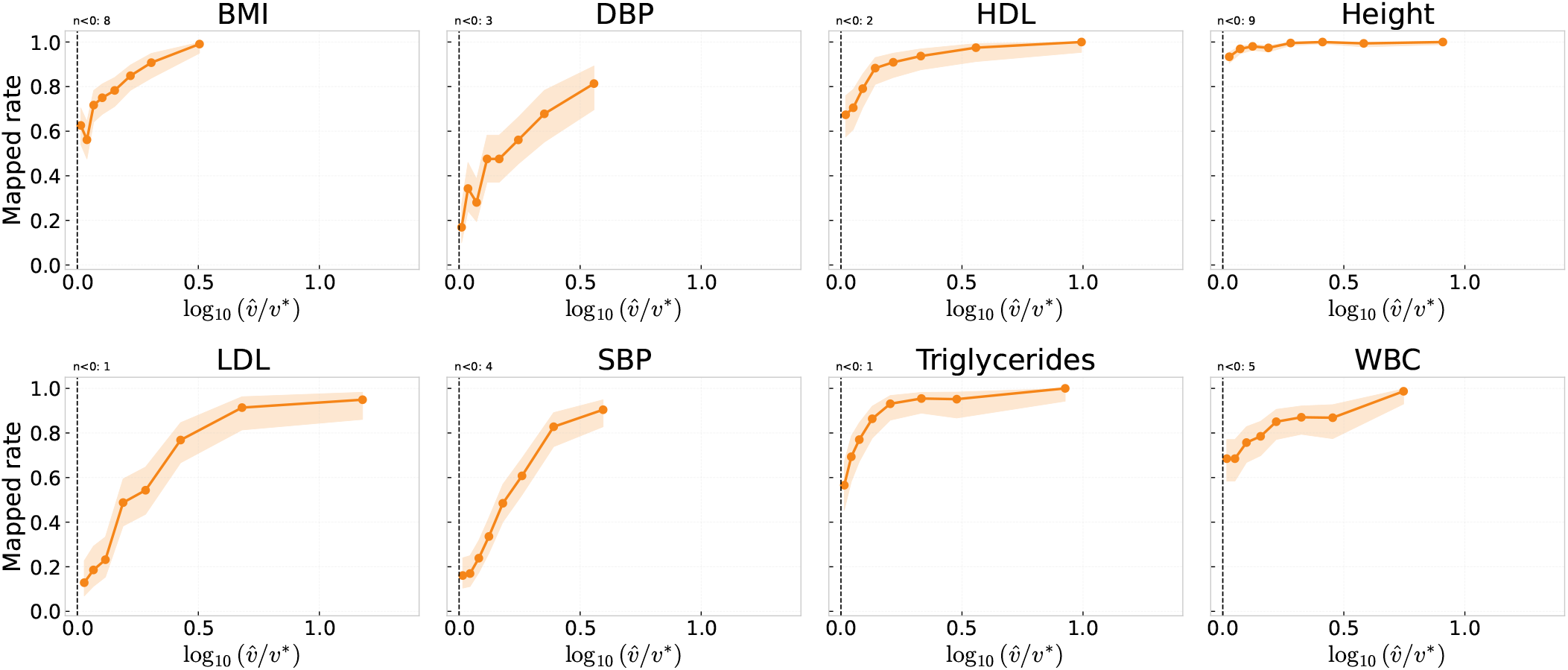
Mapping of original UKBB loci to SuSiE-X credible sets as a function of distance above the ascertainment boundary. For each UKBB trait re-analyzed with SuSiE-X, we plot the fraction of original loci that map to a reported SuSiE-X credible set against 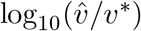, where 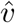 is the estimated variance contribution of the original locus and *v*^∗^ is the variance-contribution threshold corresponding to 5 × 10^−8^. Points show bin-wise mapping rates, and shaded regions show binomial proportion confidence intervals. The vertical dashed line marks the original GWAS ascertainment boundary. Values labeled *n <* 0 indicate the number of original loci falling below that boundary.

**Figure S17:**
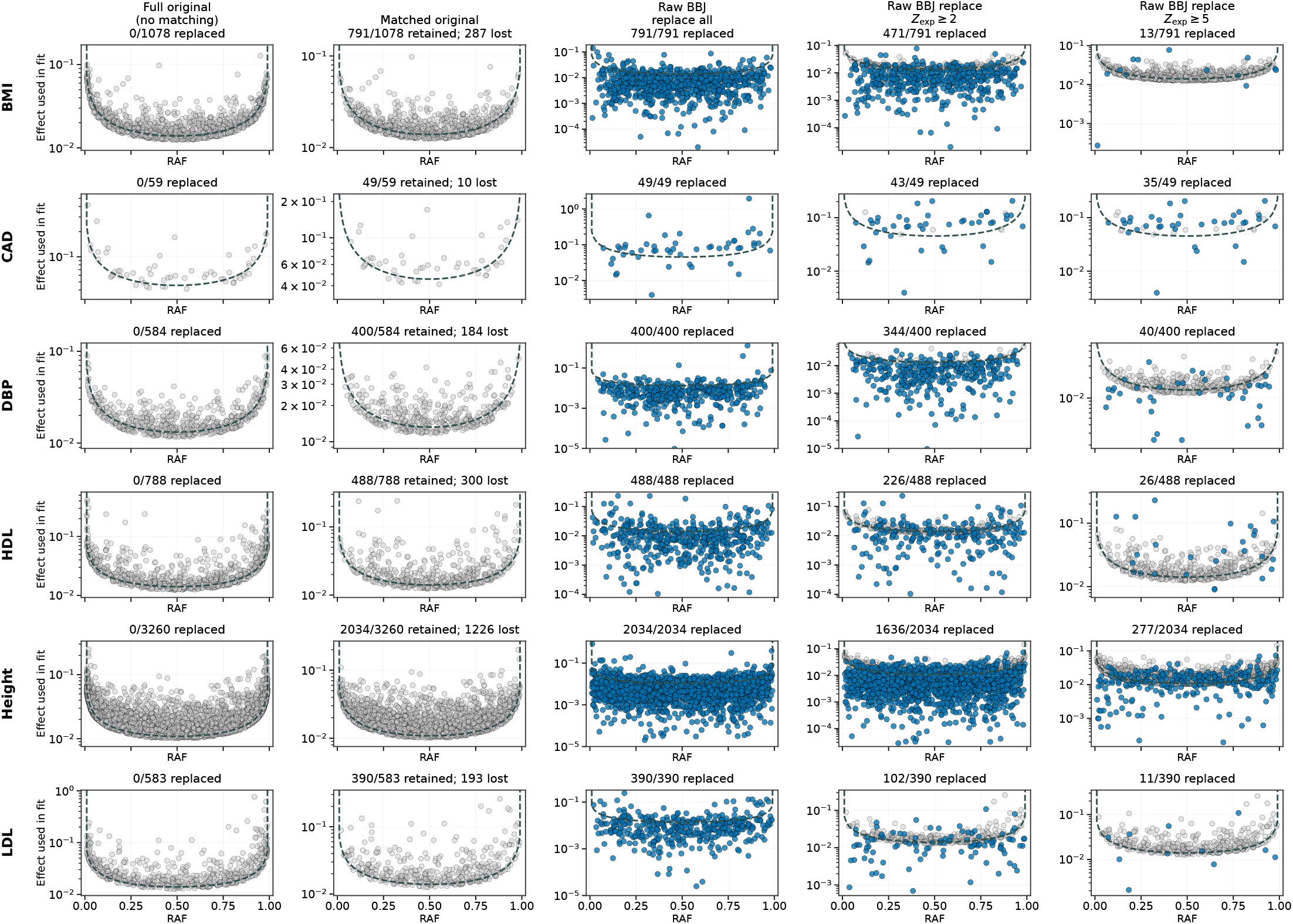
Effects of BBJ matching and BBJ effect-size replacement on smile plots. For each trait, we compare the original European-ancestry smile plot, the subset of original loci that could be matched to BBJ, and the same matched loci after replacing original effect-size estimates with BBJ effect-size estimates. Columns show the full original loci, matched original loci, BBJ replacement for all matched variants, BBJ replacement for variants with expected BBJ *Z* ≥ 2, and BBJ replacement for variants with expected BBJ *Z* ≥ 5. The matched original panels indicate how many original loci were retained after matching to BBJ and how many were lost. For disease traits, effect sizes are shown as odds-ratio minus one.

**Figure S17:**
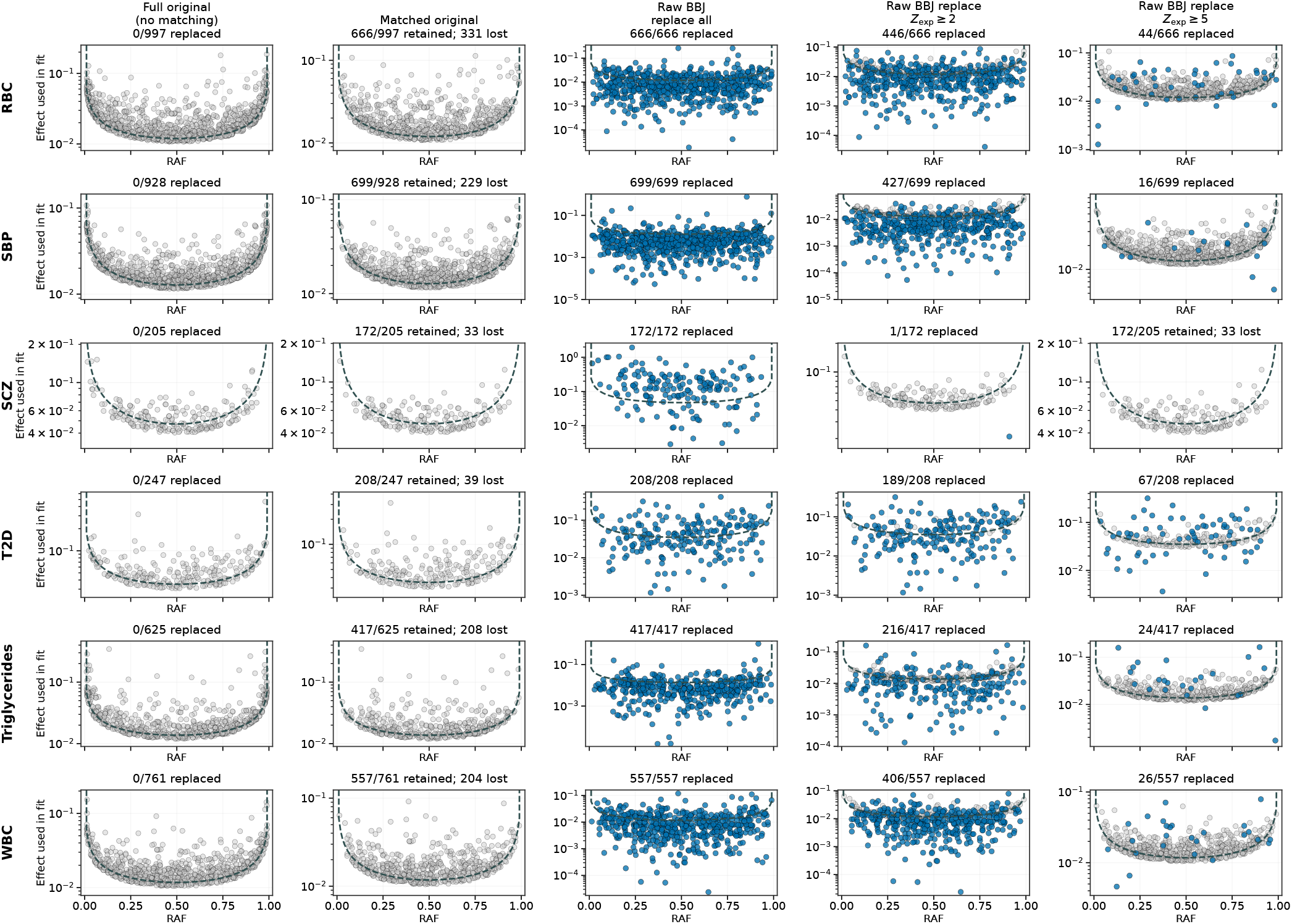
Effects of exact BBJ matching and BBJ effect-size replacement on smile plots. Continued.

**Figure S18:**
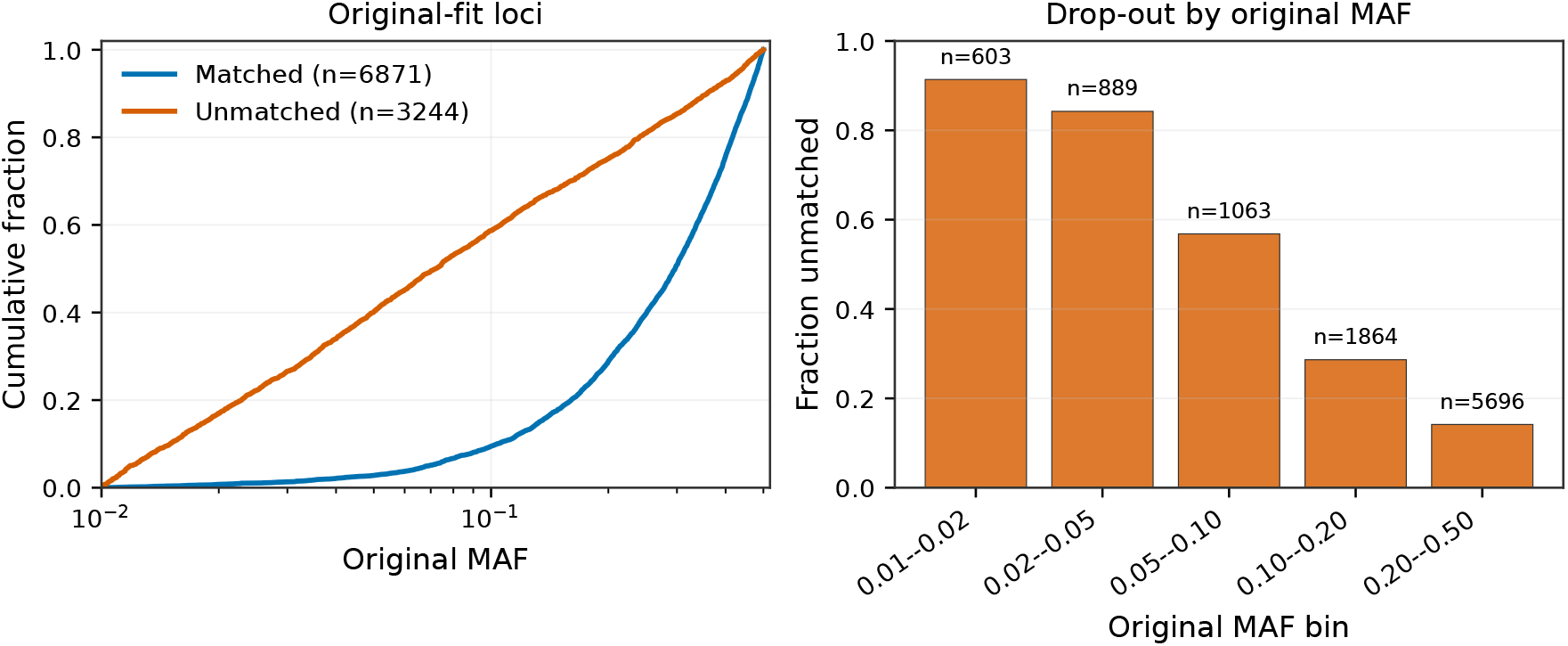
Original loci that could not be exactly matched to BBJ were enriched for rare variants. For each trait in the BBJ effect-size replacement analysis, we started with the loci included in the original model fits and determined whether the same SNP identifier had BBJ frequency and effect-size estimates. No proxy variants were used. Left panel shows the cumulative distribution of original minor allele frequencies for loci that were exactly matched to BBJ and loci that were not matched. Right panel shows the fraction of original-fit loci that were unmatched within bins of original minor allele frequency.

**Figure S19:**
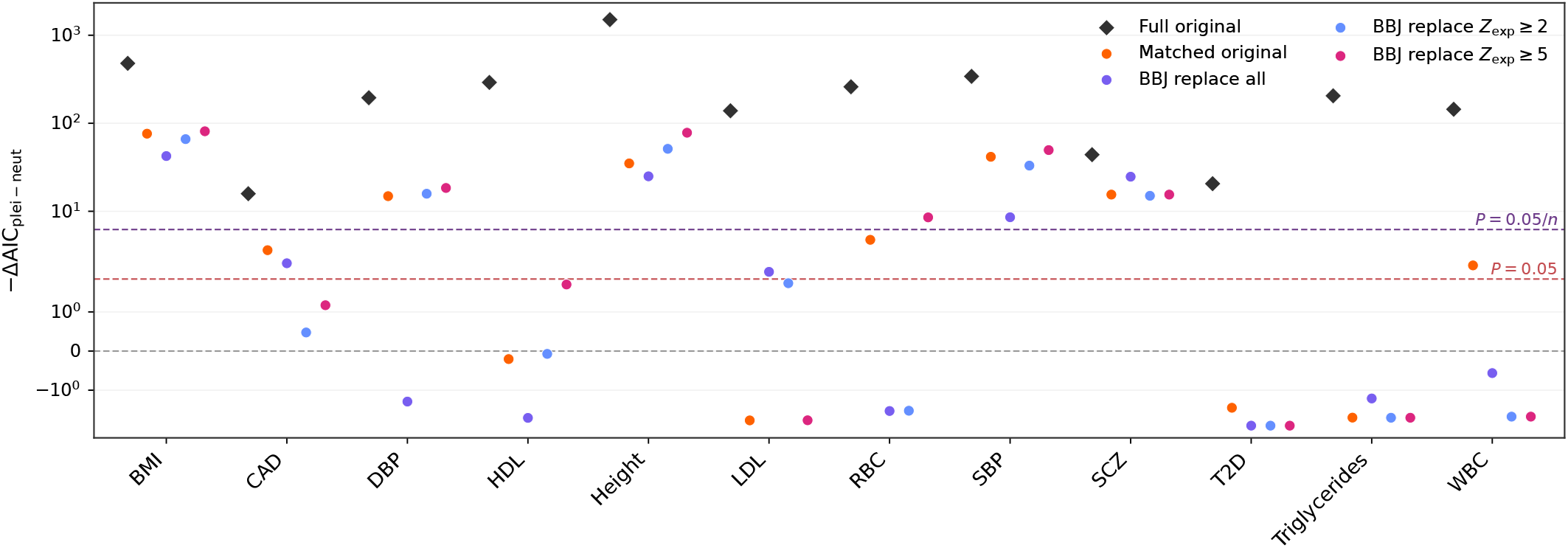
Effects of BBJ effect-size replacement on evidence for selection. We compared AIC-scale likelihood differences between selection models and neutrality for original European-ancestry loci, BBJ-matchable loci from among the original loci, and BBJ-matchable loci with effect-size replacement. Full original shows the original fits before restricting to variants that could be matched to BBJ (same as Figure 3). Matched original uses only original loci that could be matched to BBJ, while retaining the original effect estimates. BBJ replacement analyses use the same matched loci, but replace original effect sizes with BBJ effect-size estimates for all matched variants, for variants with expected BBJ *Z* ≥ 2, or for variants with expected BBJ *Z* ≥ 5. Reference lines show the corresponding one-parameter likelihood ratio test thresholds for *P* = 0.05 and Bonferroni-adjusted *P* = 0.05*/n*.

**Figure S20:**
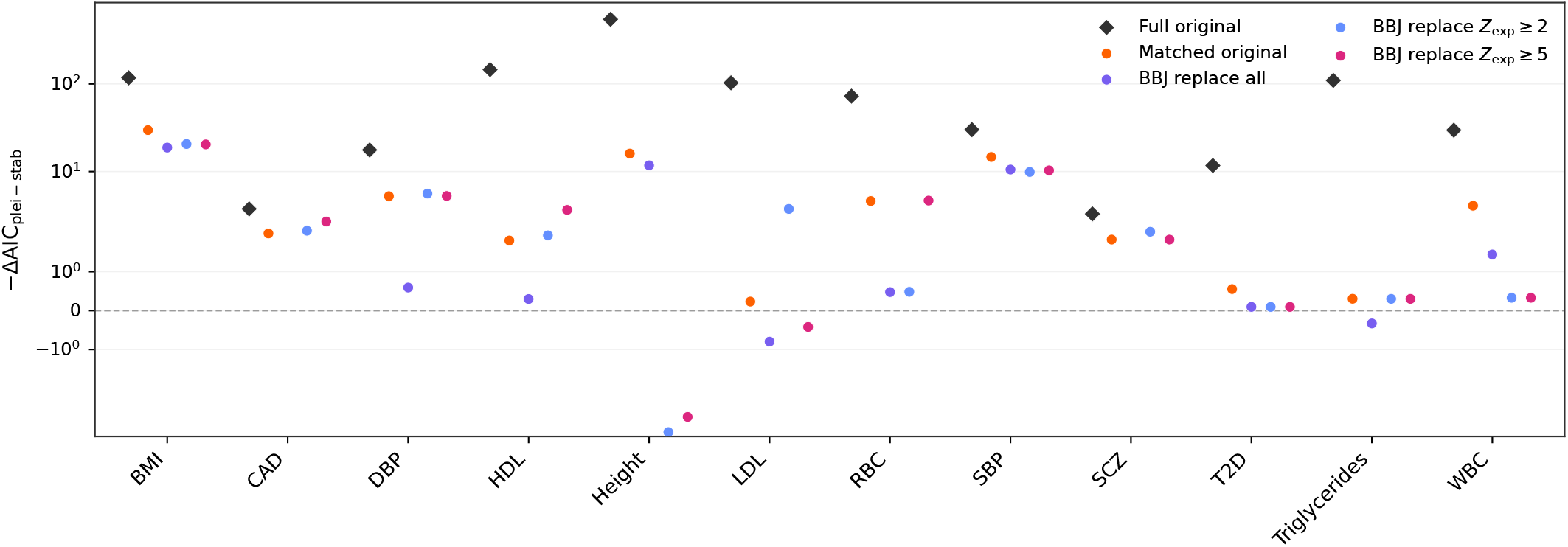
Effects of BBJ effect-size replacement on support for pleiotropic versus 1D stabilizing selection. We compared AIC-scale likelihood differences between pleiotropic and single-trait stabilizing selection models for original European-ancestry loci, BBJ-matchable loci from among the original loci, and BBJ-matchable loci with effect-size replacement. Positive values indicate support for the pleiotropic model relative to the single-trait model. Full original shows the original fits before restricting to variants that could be matched to BBJ. Matched original uses only original loci that could be matched to BBJ, while retaining the original effect estimates. BBJ replacement analyses use the same matched loci, but replace original effect sizes with BBJ effect-size estimates for all matched variants, for variants with expected BBJ *Z* ≥ 2, or for variants with expected BBJ *Z* ≥ 5.

**Figure S21:**
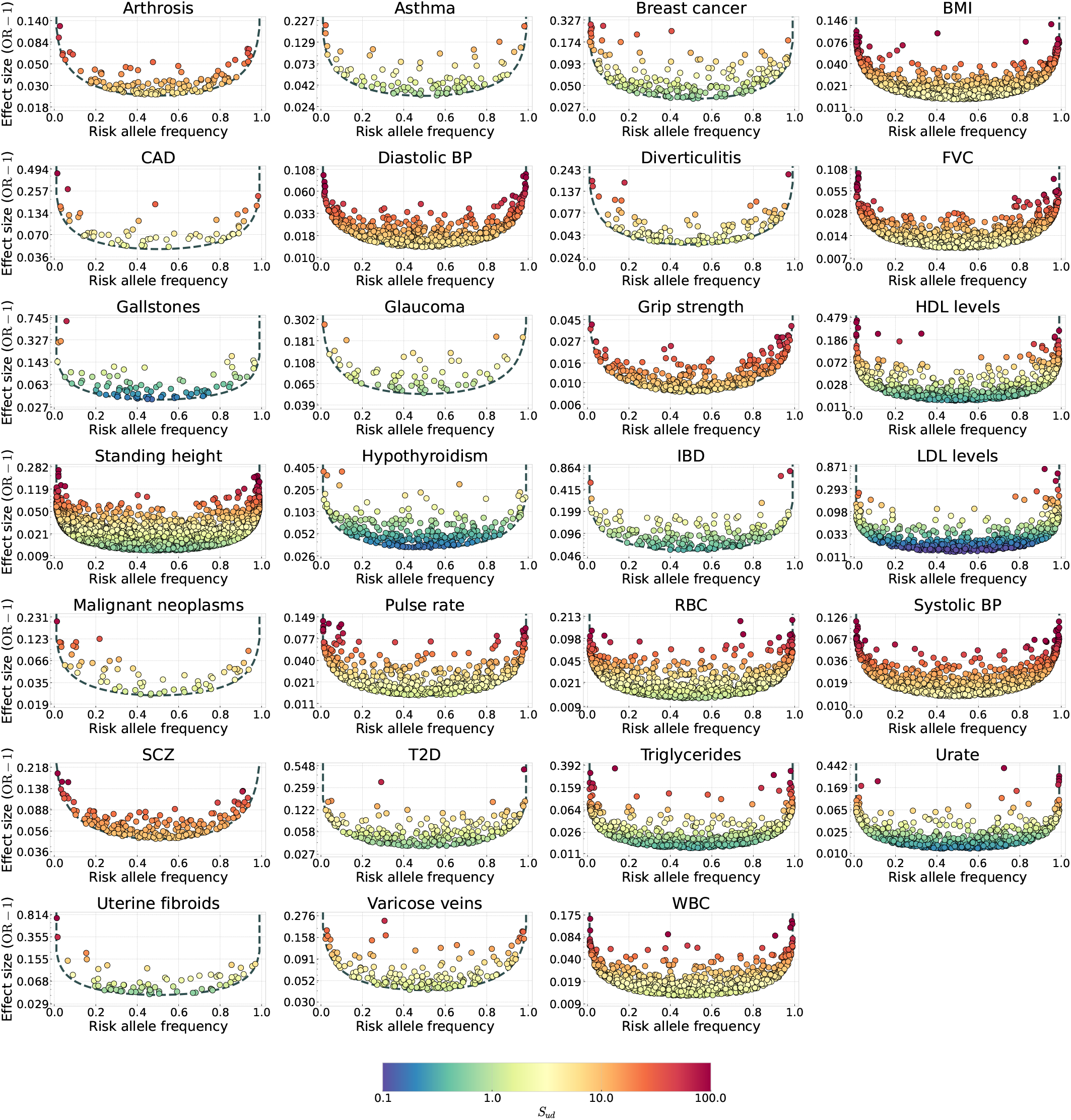
Estimated single-trait stabilizing selection coefficients on all disease and quantitative trait variants. We used maximum-likelihood estimates of selection intensities to predict the strength of selection on observed GWAS variants. Estimates represent selection coefficients under the single-trait model of stabilizing selection. Selection estimates were calculated using shrunk effect sizes, while the y-axis shows GWAS effect sizes. For disease traits, GWAS effect sizes are shown as the odds-ratio minus one. The dashed line shows the frequency cutoff at each effect size. Frequency thresholds are computed based on raw effect sizes, while the strength of selection is fit using shrunken estimates. The scaled strength of underdominant selection is calculated as *S*_*ud*_ = 2*N*_*e*_*s*_*ud*_, with *N*_*e*_ = 10,000 used to roughly represent scaled selection coefficients in an ancestral human population.

**Figure S22:**
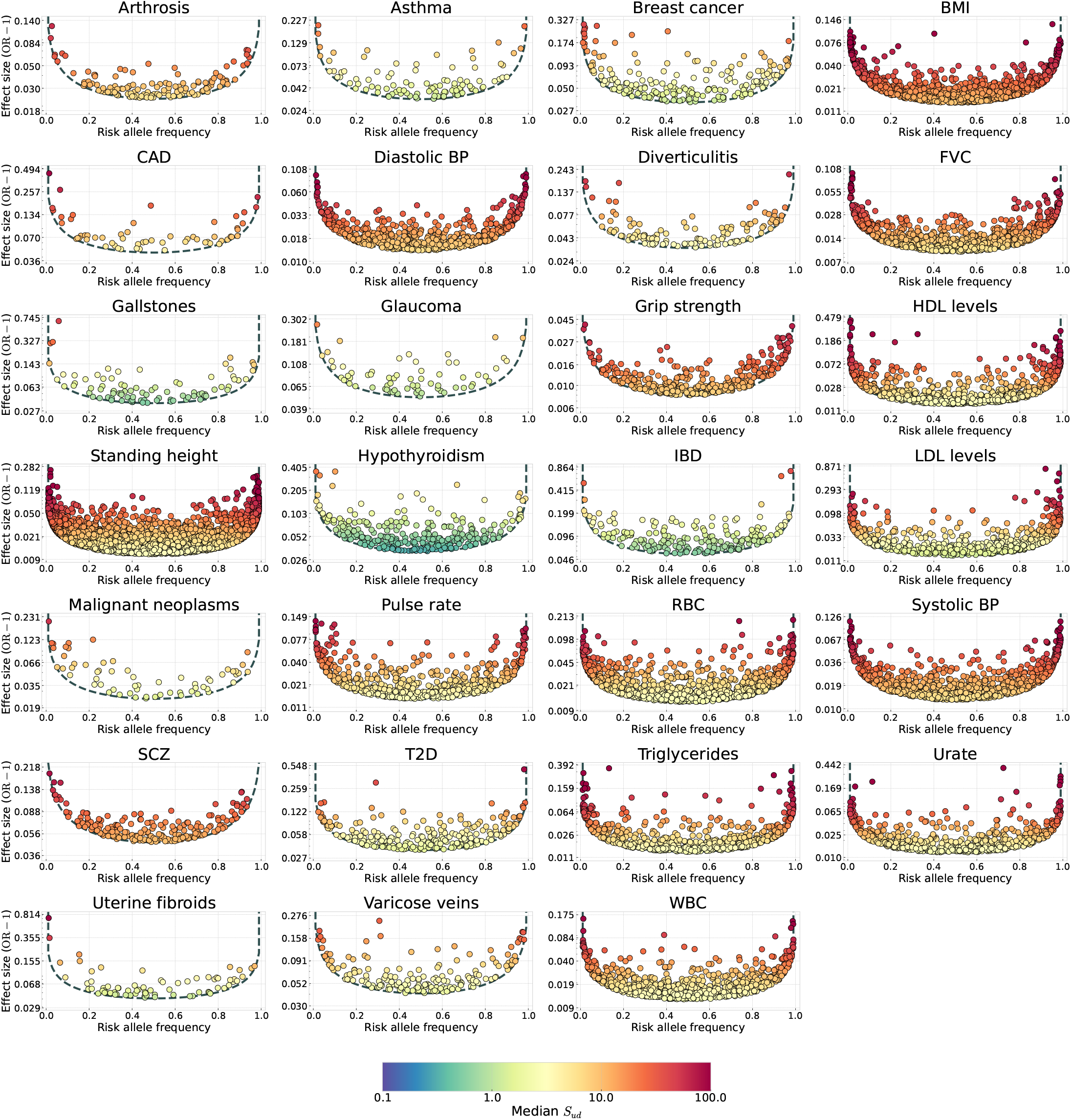
Estimated median pleiotropic stabilizing selection coefficients on all disease and quantitative trait variants. Median selection coefficients are shown given the estimated *I*_*p*_ for each trait, conditional on trait-increasing allele frequencies. The dashed line shows the frequency cutoff at each effect size. Frequency thresholds are computed based on raw effect sizes, while the strength of selection is fit using shrunken estimates. For disease traits, GWAS effect sizes are shown as the odds-ratio minus one. The scaled strength of selection is calculated as *S*_*ud*_ = 2*N*_*e*_*s*_*ud*_, with *N*_*e*_ = 10,000 used to roughly represent scaled selection coefficients in an ancestral human population.

**Figure S23:**
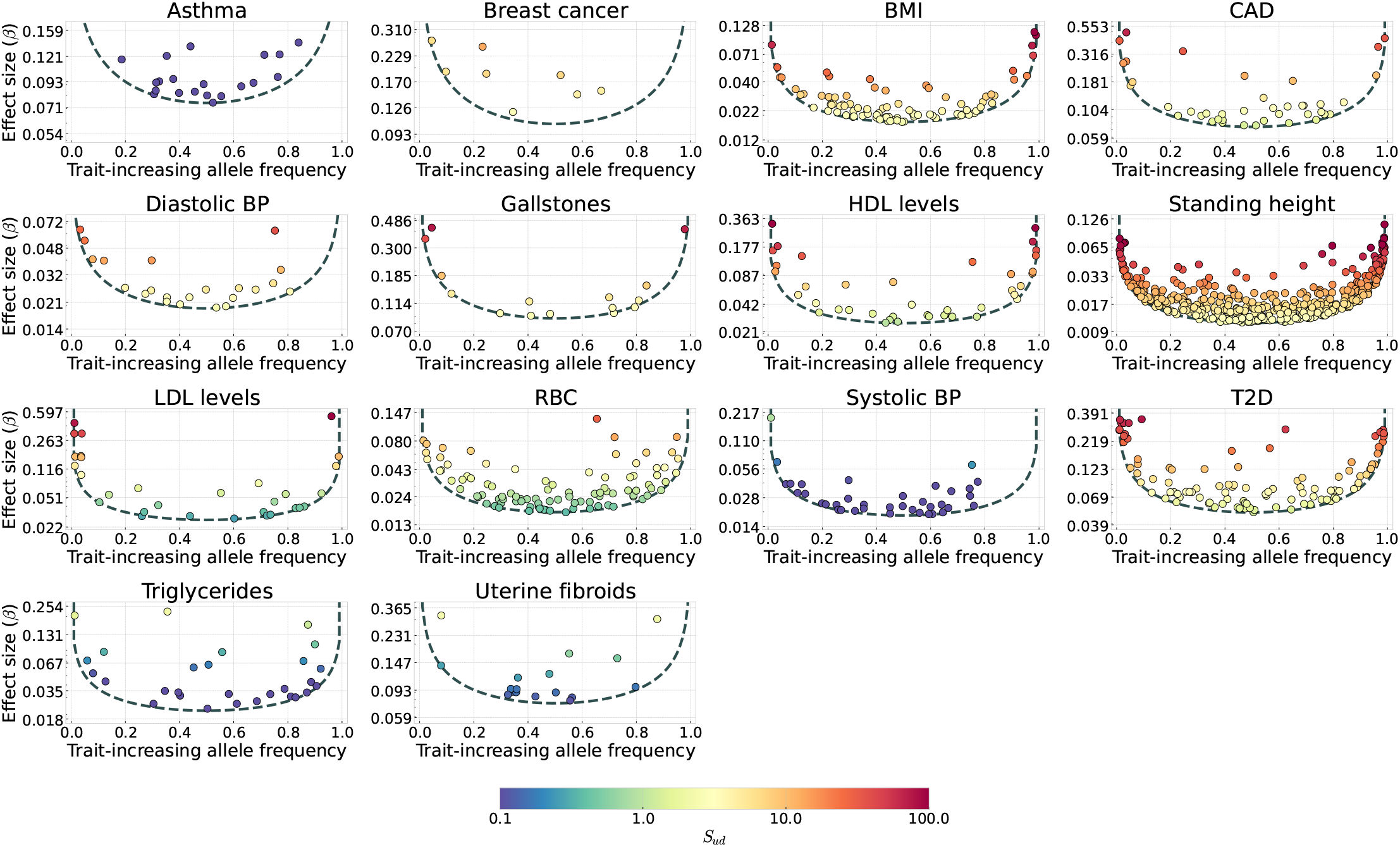
Estimated single-trait stabilizing selection coefficients on BBJ disease and quantitative trait variants. We used maximum-likelihood estimates of selection intensities to predict the strength of selection on observed GWAS variants. Estimates represent selection coefficients under the single-trait model of stabilizing selection. Selection estimates were calculated using shrunk effect sizes, while the y-axis shows GWAS effect sizes. For disease traits, GWAS effect sizes are shown as the odds-ratio minus one. BBJ loci were high-clumped using a 500 kb procedure, and likelihoods were calculated using the Japanese demographic model. The dashed line shows the frequency cutoff at each effect size. Frequency thresholds are computed based on raw effect sizes, while the strength of selection is fit using shrunken estimates. The scaled strength of underdominant selection is calculated as *S*_*ud*_ = 2*N*_*e*_*s*_*ud*_, with *N*_*e*_ = 10,000 used to roughly represent scaled selection coefficients in an ancestral human population.

**Figure S24:**
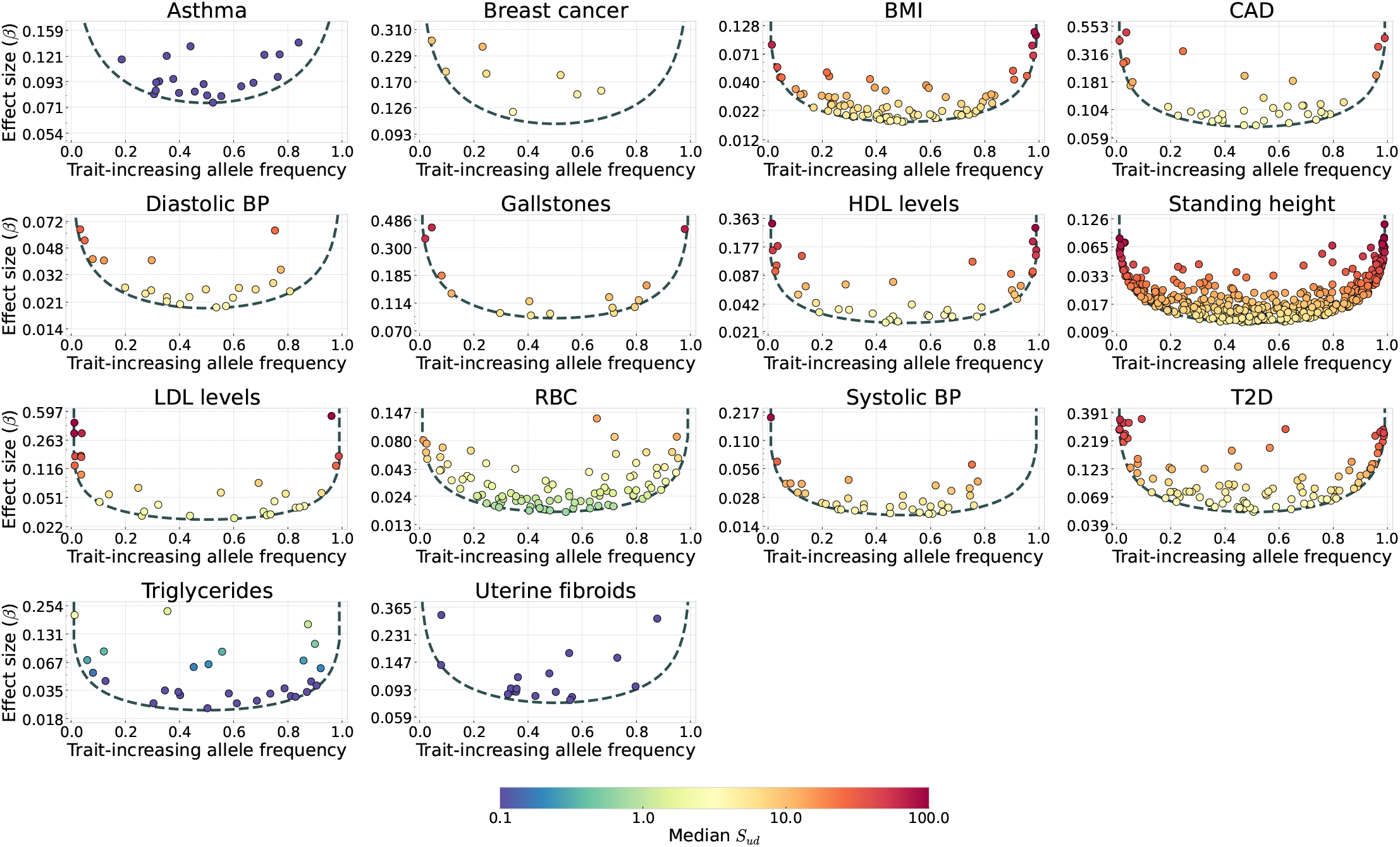
Estimated median pleiotropic stabilizing selection coefficients on BBJ disease and quantitative trait variants. Median selection coefficients are shown given the estimated *I*_*p*_ for each trait, conditional on trait-increasing allele frequencies. For disease traits, GWAS effect sizes are shown as the odds-ratio minus one. BBJ loci were high-clumped using a 500 kb procedure, and likelihoods were calculated using the Japanese demographic model. The dashed line shows the frequency cutoff at each effect size. Frequency thresholds are computed based on raw effect sizes, while the strength of selection is fit using shrunken estimates. The scaled strength of stabilizing selection is calculated as *S*_*ud*_ = 2*N*_*e*_*s*_*ud*_, with *N*_*e*_ = 10,000 used to roughly represent scaled selection coefficients in an ancestral human population.

**Figure S25:**
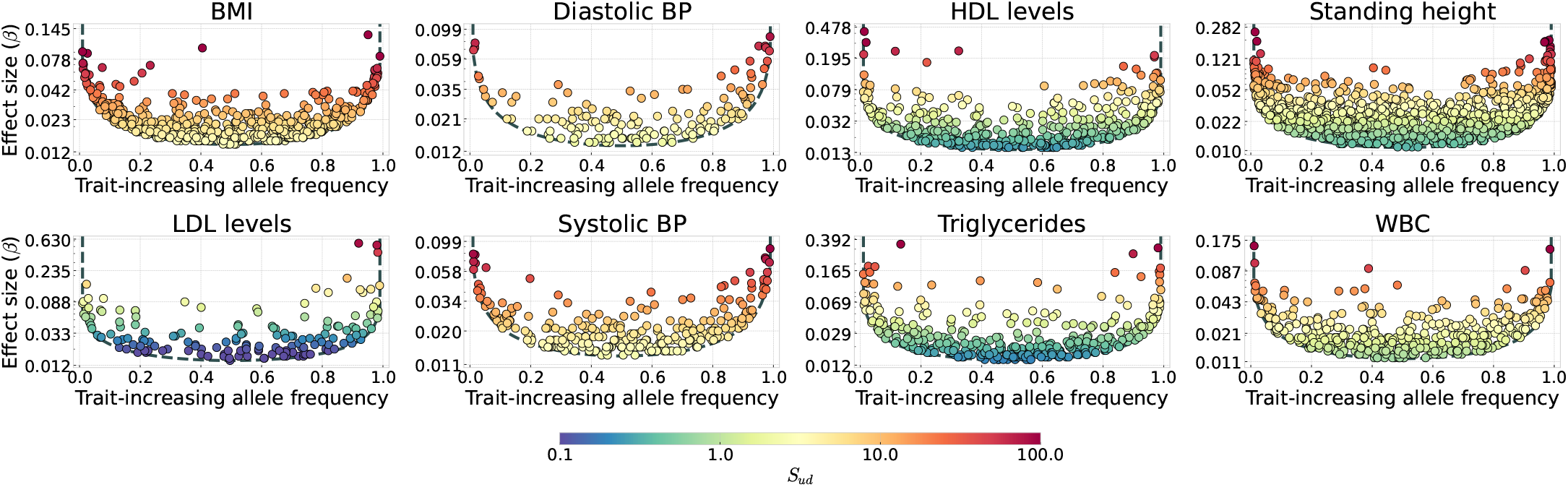
Estimated single-trait stabilizing selection coefficients on UKBB SuSiE-X quantitative trait variants. We used maximum-likelihood estimates of selection intensities to predict the strength of selection on observed GWAS variants. Estimates represent selection coefficients under the single-trait model of stabilizing selection. Selection estimates were calculated using fine-mapped effect-size estimates, while the y-axis shows fine-mapped effect sizes. One variant was sampled from each fine-mapped credible set for each trait. The dashed line shows the frequency cutoff at each effect size. Frequency thresholds are computed based on raw effect sizes, while the strength of selection is fit using fine-mapped effect-size estimates. The scaled strength of underdominant selection is calculated as *S*_*ud*_ = 2*N*_*e*_*s*_*ud*_, with *N*_*e*_ = 10,000 used to roughly represent scaled selection coefficients in an ancestral human population.

**Figure S26:**
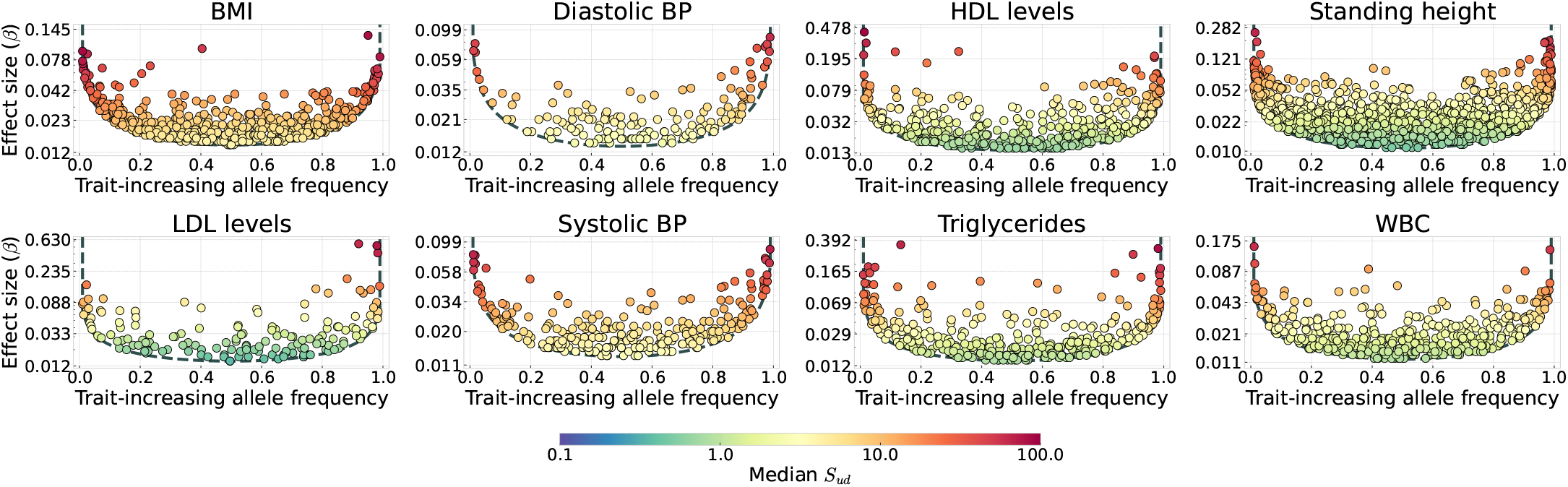
Estimated median pleiotropic stabilizing selection coefficients on UKBB SuSiE-X quantitative trait variants. Median selection coefficients are shown given the estimated *I*_*p*_ for each trait, conditional on trait-increasing allele frequencies. One variant was sampled from each fine-mapped credible set for each trait. The dashed line shows the frequency cutoff at each effect size. Frequency thresholds are computed based on raw effect sizes, while the strength of selection is fit using fine-mapped effect-size estimates. The scaled strength of stabilizing selection is calculated as *S*_*ud*_ = 2*N*_*e*_*s*_*ud*_, with *N*_*e*_ = 10,000 used to roughly represent scaled selection coefficients in an ancestral human population.

**Figure S27:**
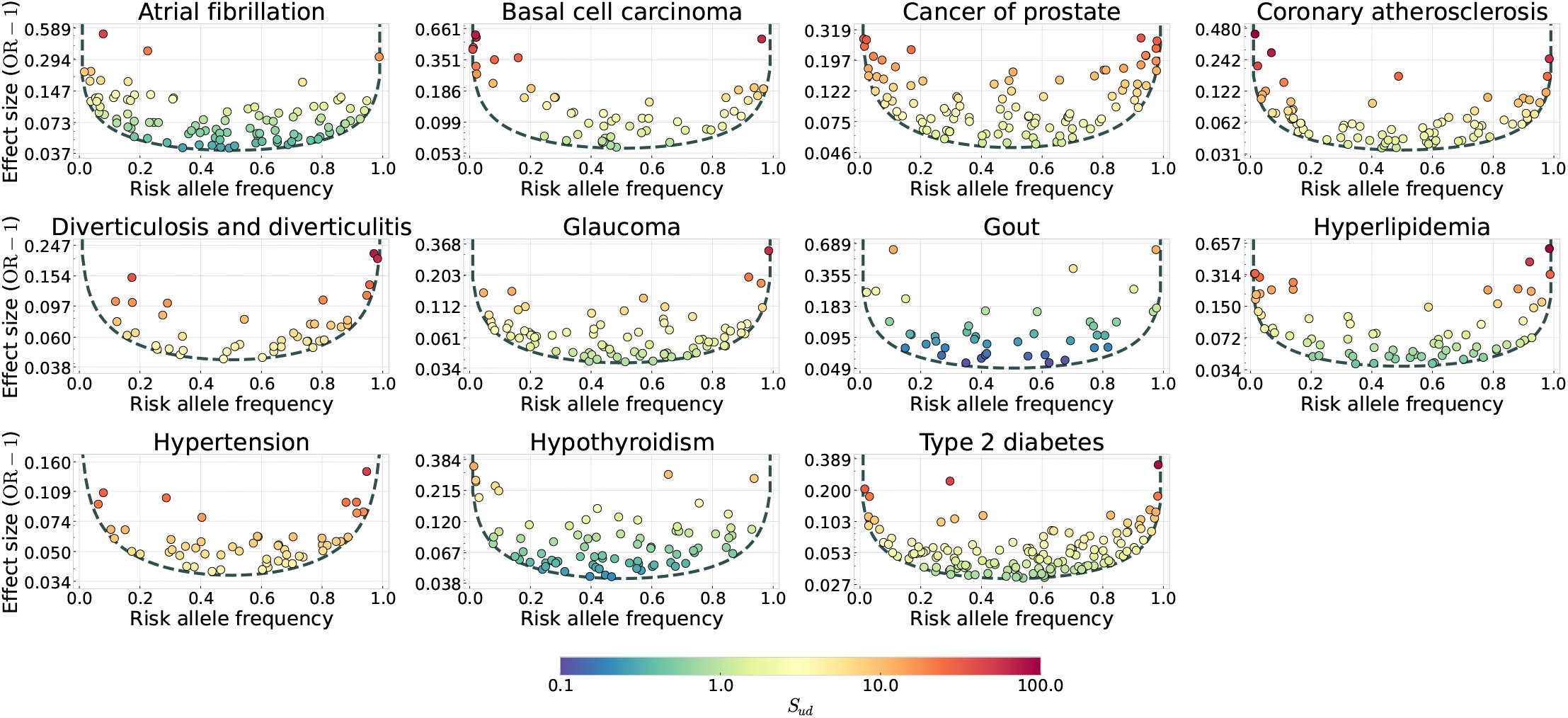
Estimated single-trait stabilizing selection coefficients on MVP disease traits. We used maximum-likelihood estimates of selection intensities to predict the strength of selection on observed GWAS variants. Estimates represent selection coefficients under the single-trait model of stabilizing selection. Selection estimates were calculated using fine-mapped effect-size estimates, while the y-axis shows fine-mapped effect sizes. For disease traits, fine-mapped effect sizes are shown as the odds-ratio minus one. One variant was sampled from each fine-mapped credible set for each trait. The dashed line shows the frequency cutoff at each effect size. Frequency thresholds are computed based on raw effect sizes, while the strength of selection is fit using fine-mapped effect-size estimates. The scaled strength of underdominant selection is calculated as *S*_*ud*_ = 2*N*_*e*_*s*_*ud*_, with *N*_*e*_ = 10,000 used to roughly represent scaled selection coefficients in an ancestral human population.

**Figure S28:**
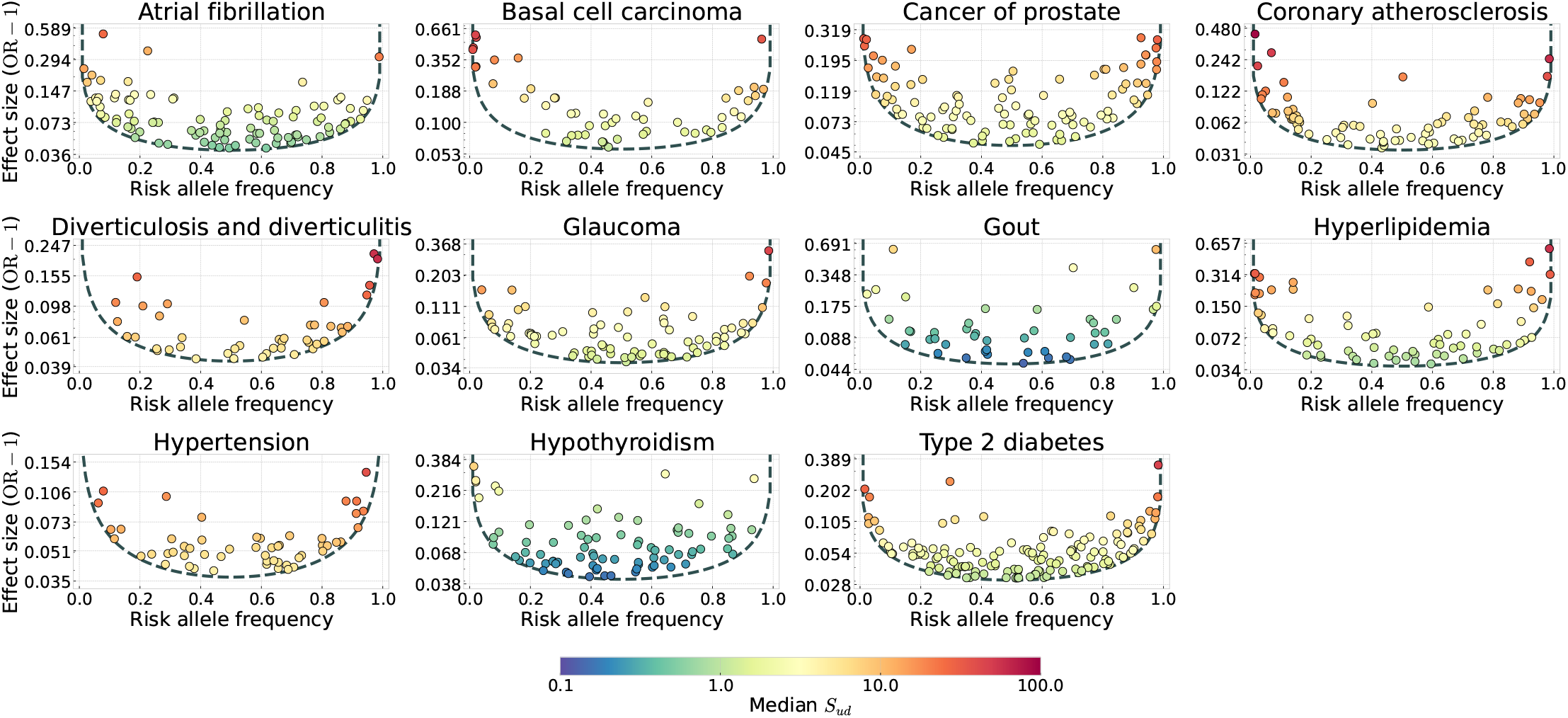
Estimated median pleiotropic stabilizing selection coefficients on MVP disease traits. Median selection coefficients are shown given the estimated *I*_*p*_ for each trait, conditional on risk allele frequencies. One variant was sampled from each fine-mapped credible set for each trait. For disease traits, fine-mapped effect sizes are shown as the odds-ratio minus one. The dashed line shows the frequency cutoff at each effect size. Frequency thresholds are computed based on raw effect sizes, while the strength of selection is fit using fine-mapped effect-size estimates. The scaled strength of stabilizing selection is calculated as *S*_*ud*_ = 2*N*_*e*_*s*_*ud*_, with *N*_*e*_ = 10,000 used to roughly represent scaled selection coefficients in an ancestral human population.

**Figure S29:**
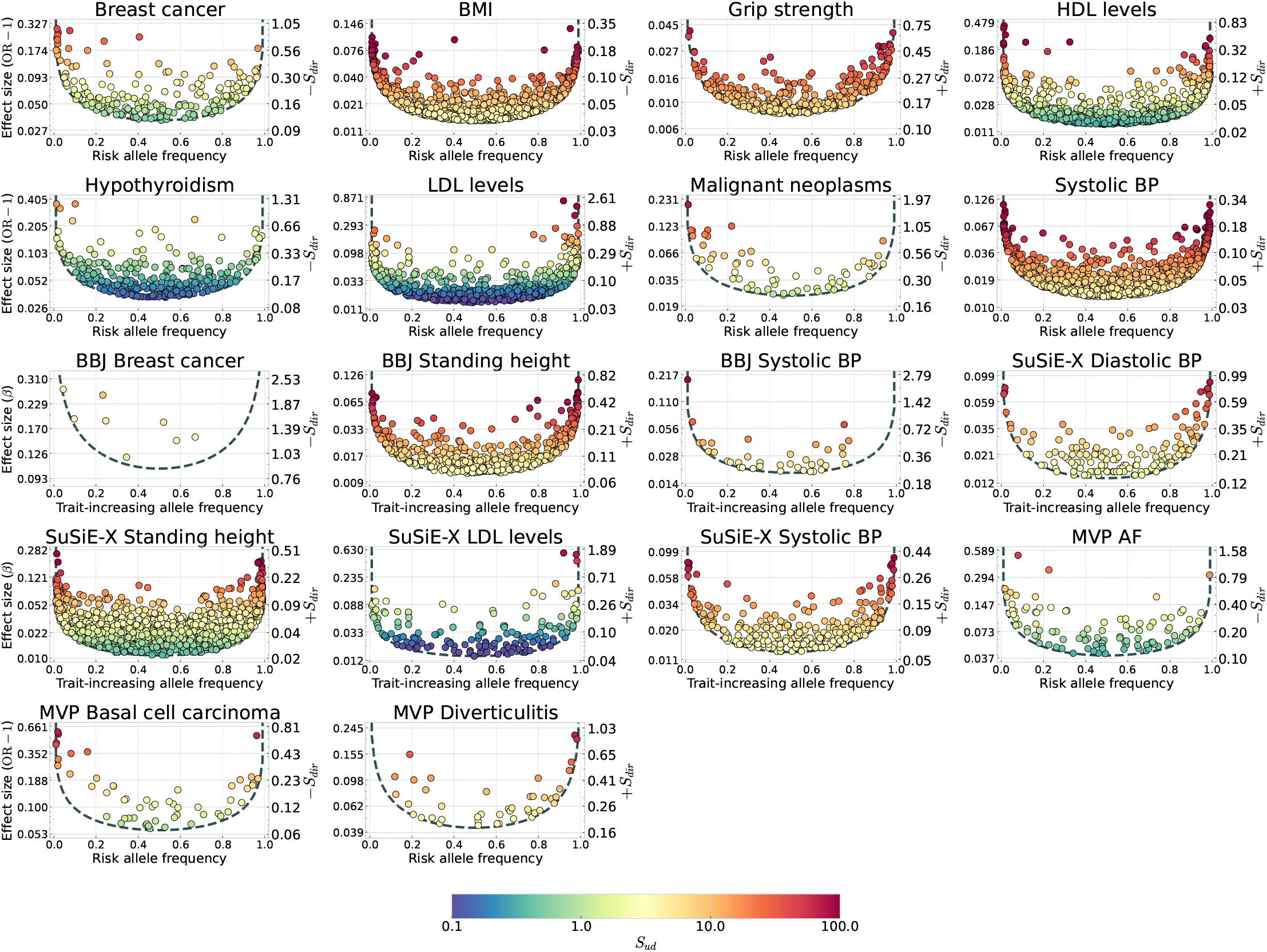
Estimated selection coefficients for the combined directional and stabilizing selection model across datasets. Equivalent structure to Figure S21, except that the right y-axis indicates the strength of directional selection while the color indicates the strength of stabilizing selection. We show traits from the original, BBJ, UKBB SuSiE-X, and MVP analyses for which the combined directional and stabilizing model has a lower AIC than the single-trait stabilizing model. Dataset-specific prefixes in panel titles indicate the source of each trait. + indicates selection against trait-decreasing (protective) alleles while − indicates selection against trait-increasing (risk) alleles.

**Figure S30:**
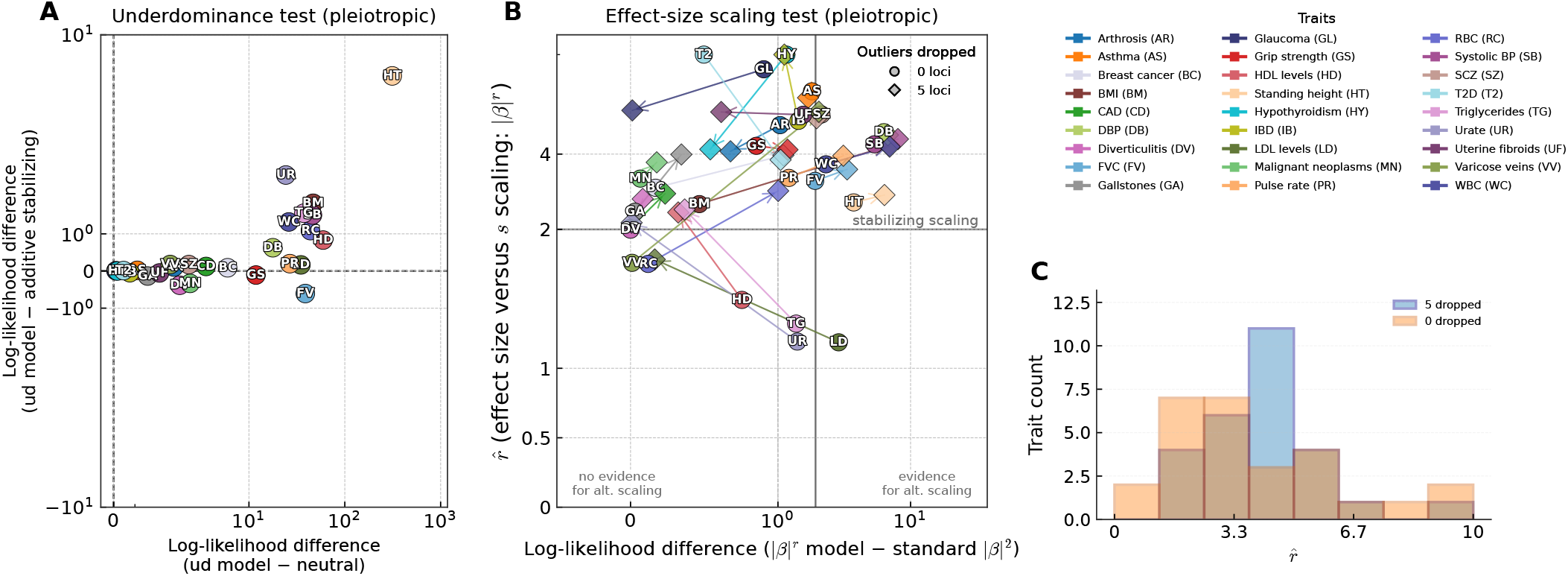
Consistency with underdominant selection and the effect size to selection scaling predicted by stabilizing selection under maximum likelihood shrinkage. We repeated the Figure 4 analysis using variant effects adjusted with maximum likelihood winner’s curse procedure instead of ashr posterior means.

**Figure S31:**
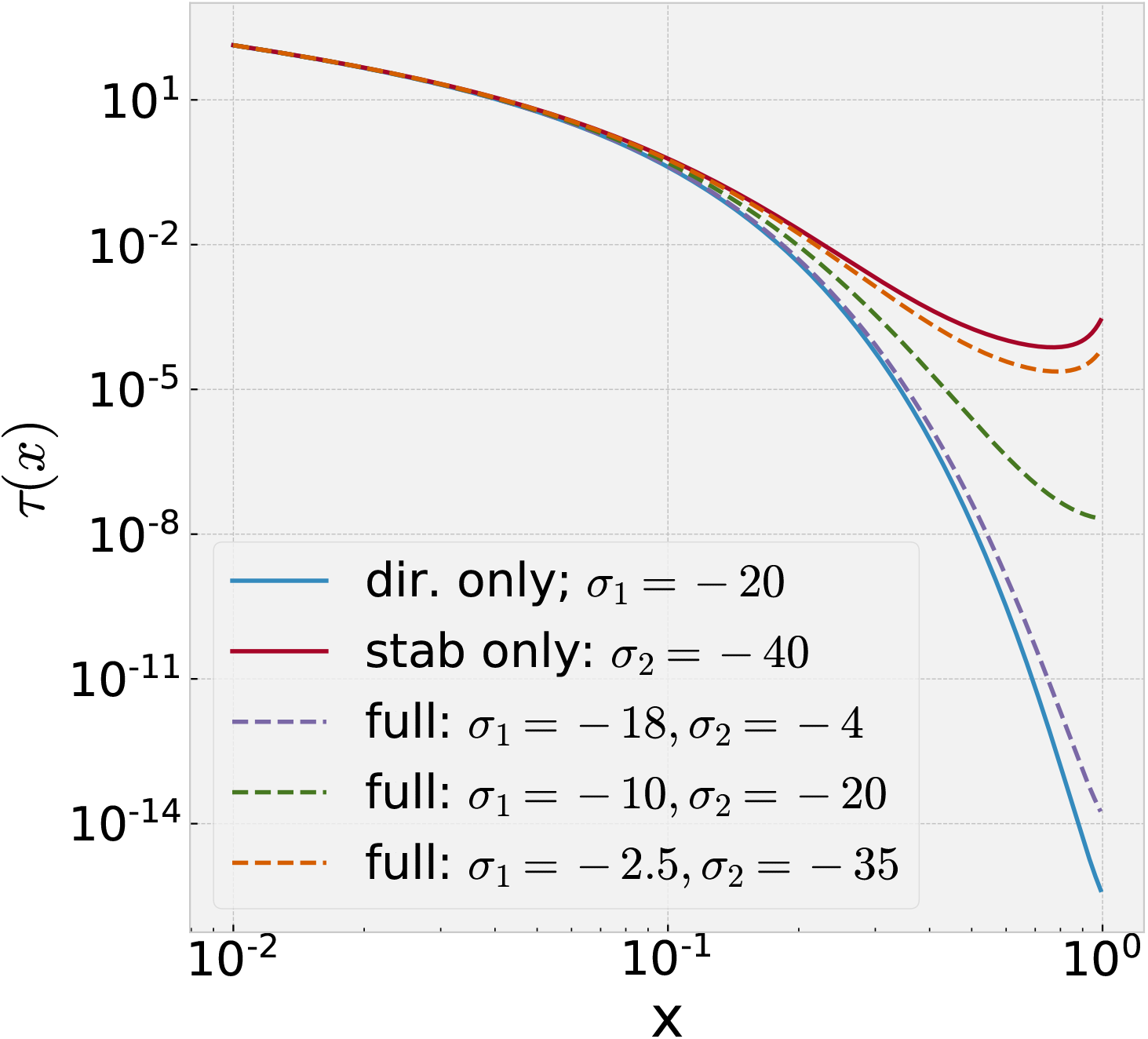
The allele frequency distribution with both additive and underdominant selection interpolates between the two models. At low frequencies the SFS under directional (additive) and stabilizing (underdominant) selection look similar, but more high frequency variants are expected for stabilizing selection. An underdominant scaled selection coefficient twice as large is required to match to additive selection.

**Figure S32:**
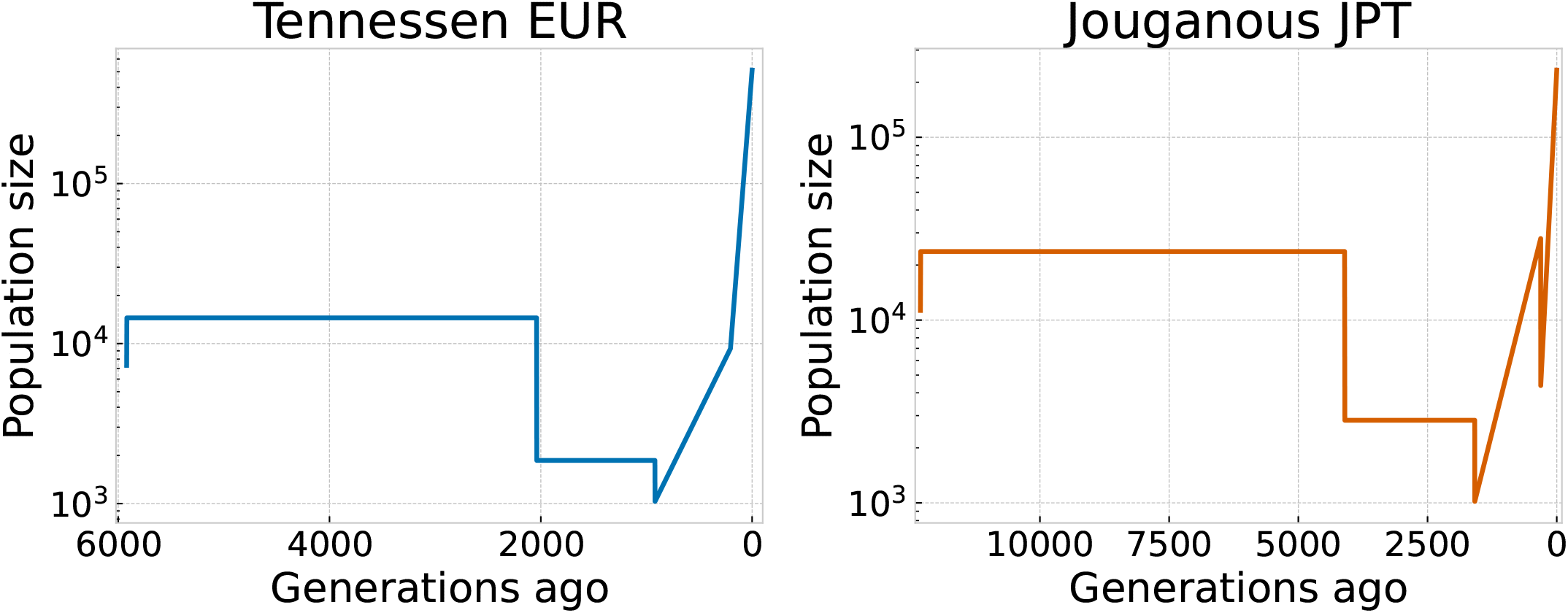
Demographic history models used to generate allele frequency distributions. We implemented the demographic model of non-Finnish European population history fit by [57] and the model of Japanese population history fit by [59]. The original models also included periods of migration between African and between European and Asian populations that we have omitted. Populations sizes were calibrated using a mutation rate of 2.36 × 10^−8^ per site per generation for the EUR model and 1.44 × 10^−8^ for the JPT model.

**Figure S33:**
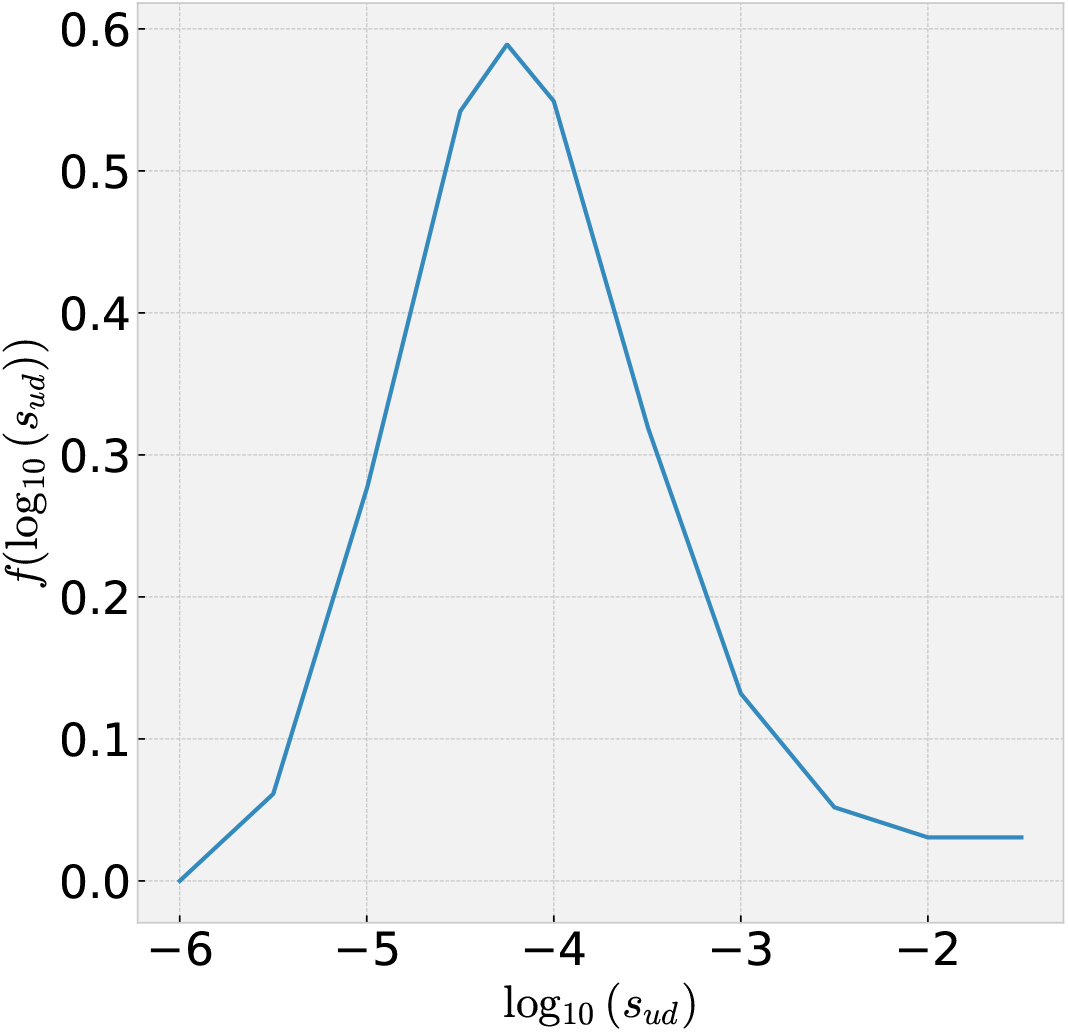
Approximate single shared DFE (SSD). We approximated the DFE inferred by Simons *et al*. (2025) [38] using a piece-wise linear function on their Figure 3b. The density shown is a probability distribution on log_10_(*s*_*ud*_), rather than the density of *s*_*ud*_ plotted on a log scale.

**Figure S34:**
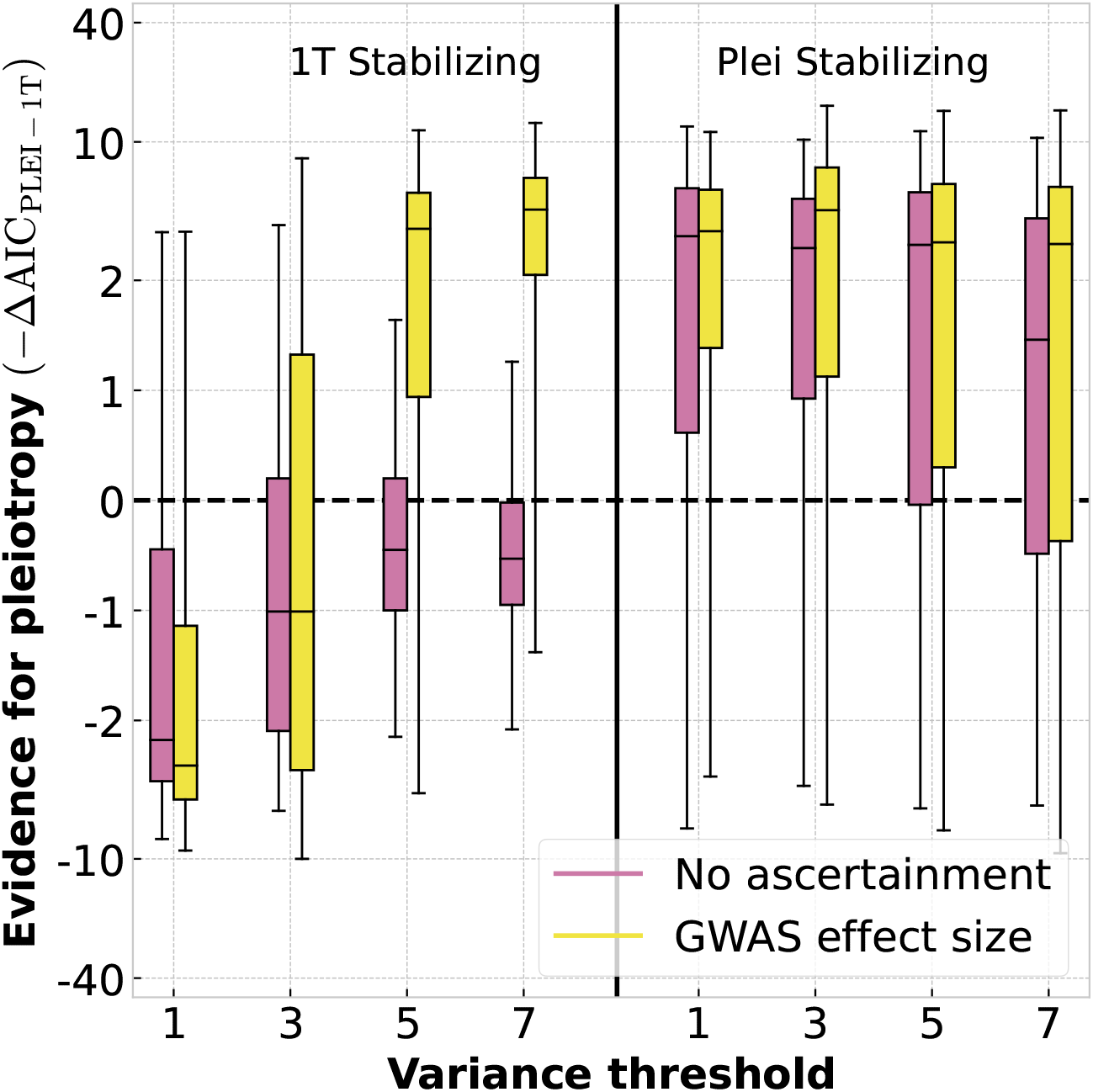
Discerning pleiotropic versus single-trait stabilizing selection in simulations. Simulations performed using the DFE estimated by Simons *et al*. [38] under models of single-trait and pleiotropic stabilizing selection. Positive values indicate support for the pleiotropic model and negative values indicate support for the single-trait model. We compare simulations where effect sizes are known without error to a model of GWAS ascertainment at a *p*-value threshold of 5 × 10^−8^. Increasing the variance threshold corresponds to more strongly selected variants being included and greater winner’s curse when ascertainment bias is modeled. Each simulation was fixed at 500 ascertained variants. In the absence of ascertainment (pink boxes), the true model does have a higher likelihood in our method on average. However, there is substantial variance such that this is not true of every simulation. The power to discriminate between the two stabilizing selection models decreases as the ascertainment threshold is raised. This is because strongly selected variants behave similarly under both models. In simulations that included GWAS noise, when single-trait selection was the true model, it was possible for the pleiotropic model to have a higher likelihood when the variance threshold was high.

**Figure S35:**
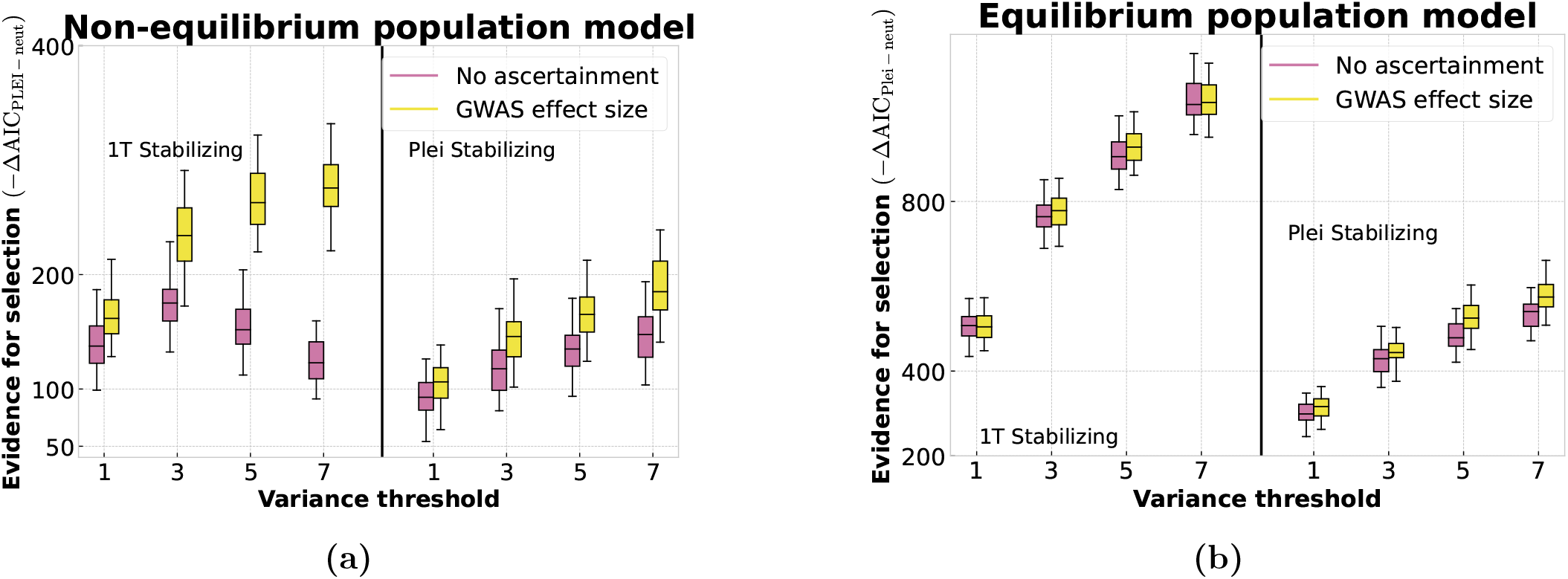
All SSD simulations have strong evidence for selection. Panel (a) shows the increase in likelihood for the stabilizing selection models relative to neutrality for DFE simulations using the non-Finnish European (NFE) demography, while panel (b) shows results for the same DFE and an effective population size of *N*_*e*_ = 10 000. The body of each boxplot represents the 25 and 75 percentiles of 500 simulations, while whiskers represent the 2.5 and 97.5 percentiles. Without modeling ascertainment bias, the evidence for selection initially increases as the variance-contribution threshold increases because alleles with larger effect sizes, and therefore variance contributions will be under stronger selection. However, the average evidence actually declines for one-dimensional selection and the NFE model as ascertained variance concentrate at the bottom of the U-shaped ascertainment curve. Ascertainment bias has a minor effect in equilibrium simulations, but favors the selection model in NFE simulations.

**Figure S36:**
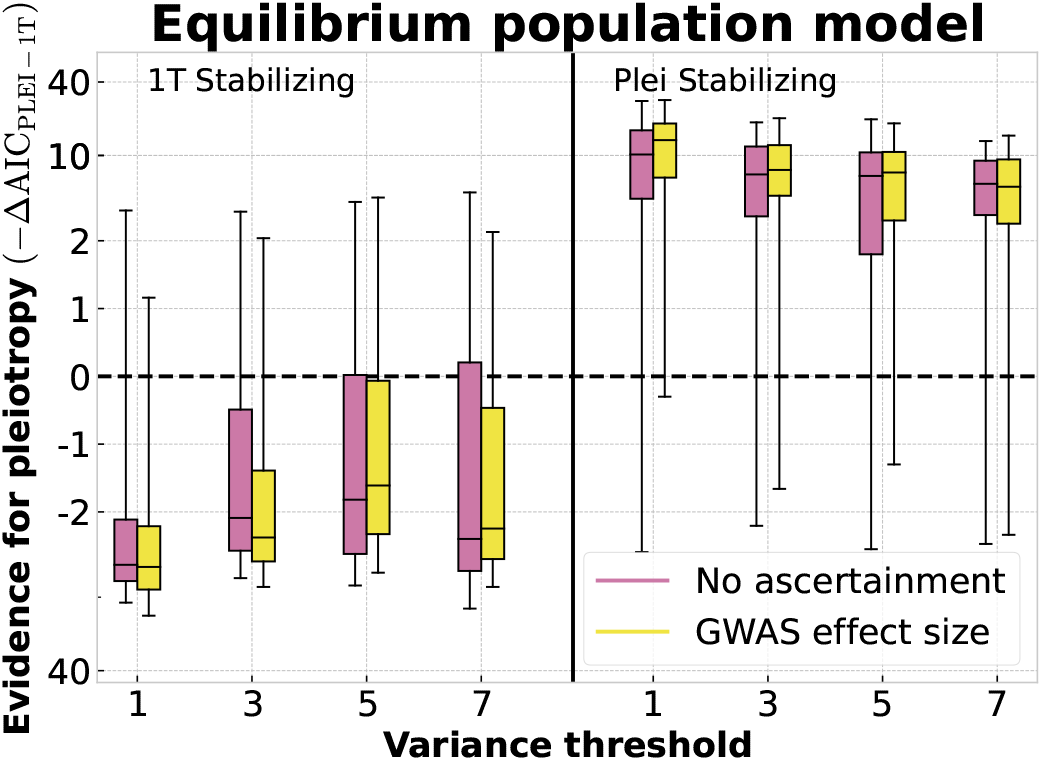
Ability to discriminate single-trait and pleiotropic models of stabilizing selection in an equilibrium population. Log-likelihood differences between the two stabilizing selection models in simulations of a population at equilibrium (*N*_*e*_ = 10 000) using the SSD DFE. The body of each boxplot represents the 25 and 75 percentiles of 500 simulations, while whiskers represent the 2.5 and 97.5 percentiles. There is greater power to discriminate between models than under simulations using the NFE demography, but the probability of false-positives remains non-negligible. In contrast to non-equilibrium simulations, ascertainment bias does not strongly affect results.

**Figure S37:**
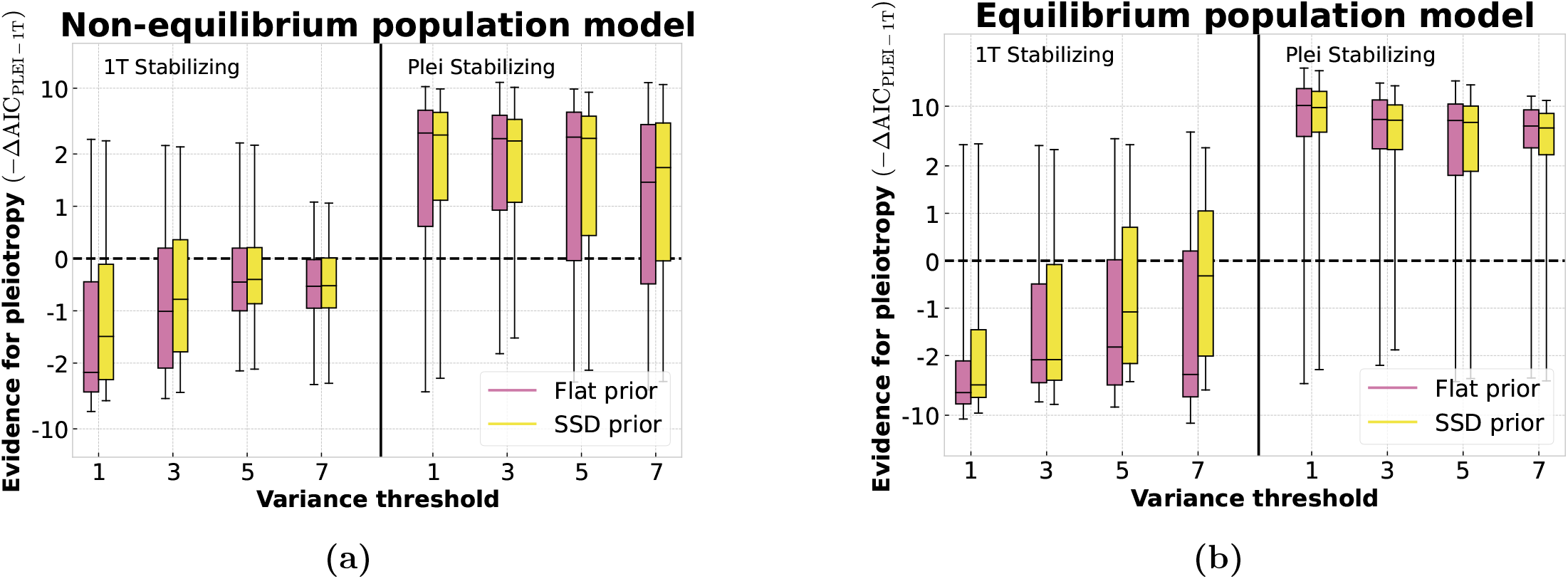
Little effect of using the SSD as a prior distribution during likelihood computations. Log-likelihood differences between the two stabilizing selection models in simulations of a population at equilibrium (*N*_*e*_ = 10 000) using the SSD DFE the simulate variance. The body of each boxplot represents the 25 and 75 percentiles of 500 simulations, while whiskers represent the 2.5 and 97.5 percentiles. During inference, we test the effect of using the SSD as a prior when calculating *p*(*s*_*ud*_ | *β*) ∝ *p*(*β* | *s*_*ud*_)*p*(*s*_*ud*_) rather than a flat prior on log *s*_*ud*_.

**Figure S38:**
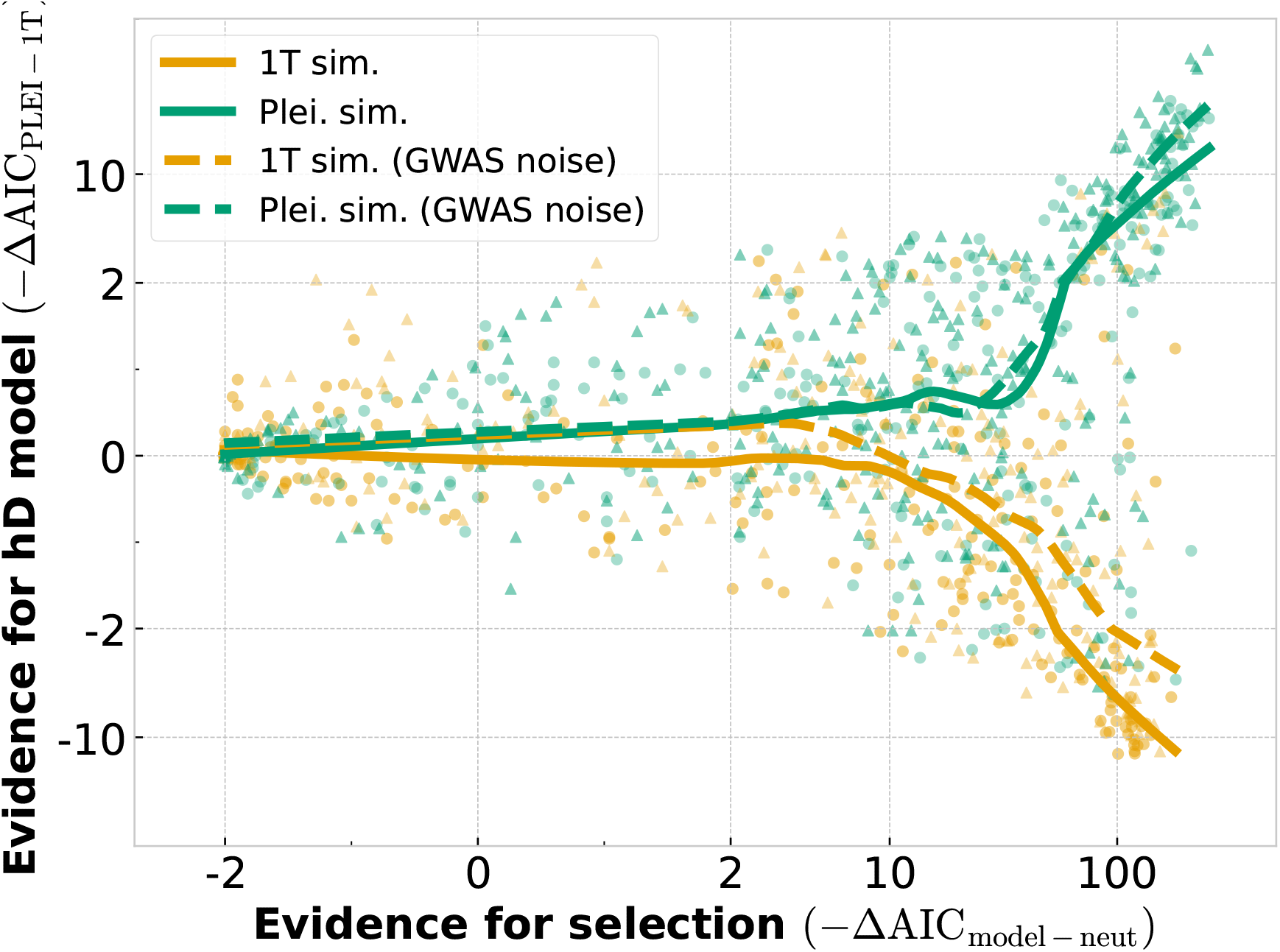
Power to discriminate between single-trait and pleiotropic stabilizing selection in simulation. Figure is based on trait-based simulations using the estimated prior distribution of IBD and a range of *I*_2_ and *I*_*p*_ values. The evidence for selection is determined by both the strength of selection and sampling noise. Circles and bold lines indicate simulations without GWAS noise, such that effect sizes are known without error. Triangles and dashed lines are for simulations where ascertainment bias was simulated and estimated effect sizes were treated as true effects for likelihood calculations. Lines show LOESS curves fit to each simulation category.

**Figure S39:**
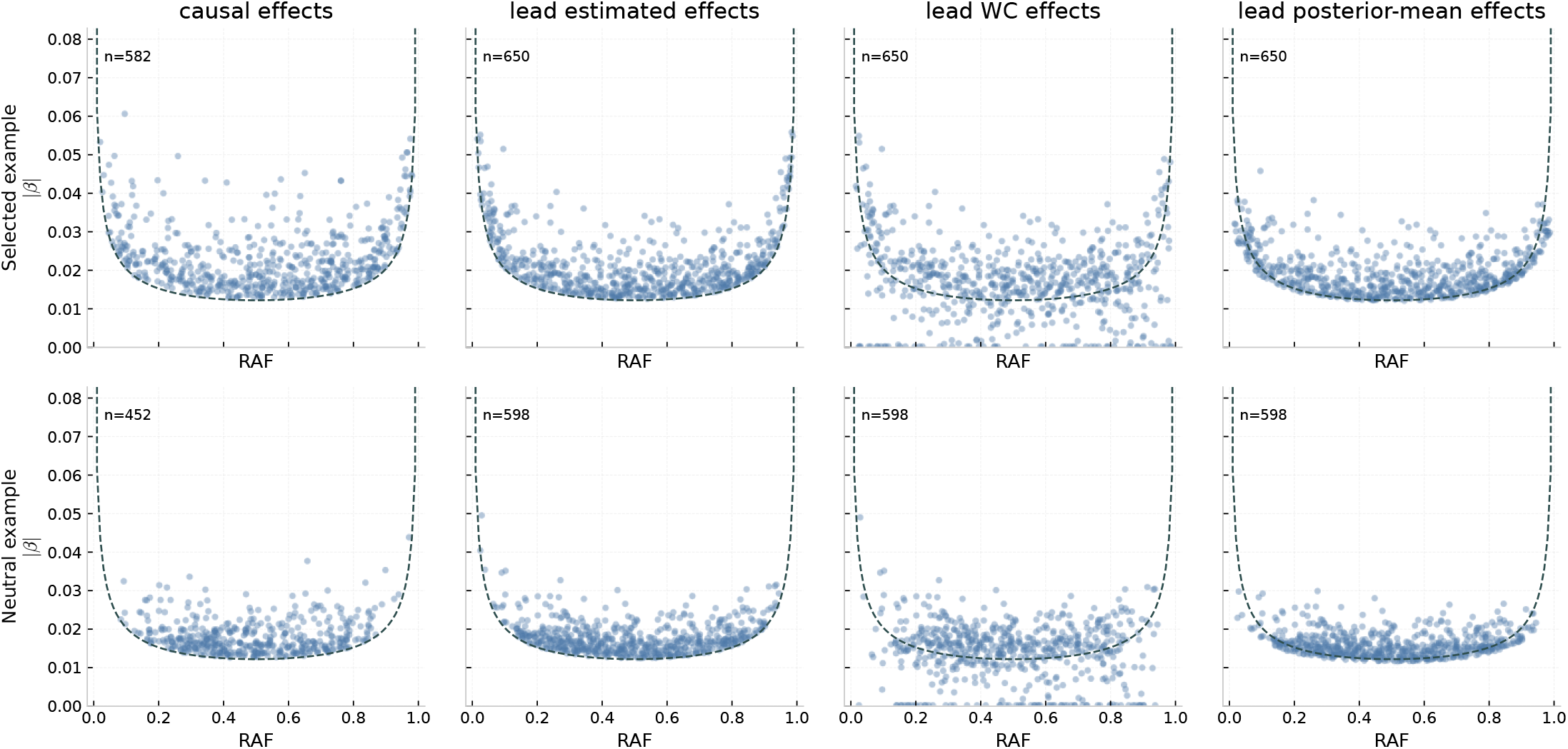
Distributions of GWAS hits from simulations using a linkage-disequilibrium graphical model. We simulated variants under a linkage-disequilibrium graphical model parameterized to produce realistic ranges of effect sizes, allele frequencies, and numbers of associated loci. The first row shows simulations under an alpha model relationship between effect sizes and frequencies. The second row shows neutral simulations with no relationship between effect size and frequency. The first column panels show causative variants and their true effect sizes, second column panels show lead variants and their marginal effect sizes, third column panels show the same lead variants after applying the winner’s-curse shrinkage to marginal effect sizes, and fourth column panels show lead variants after applying the posterior mean adjustment. Dashed curves indicate the variance-contribution threshold corresponding to a genome-wide significance cutoff of 5 × 10^−8^ computed using the effective sample size of the simulated GWAS.

**Figure S40:**
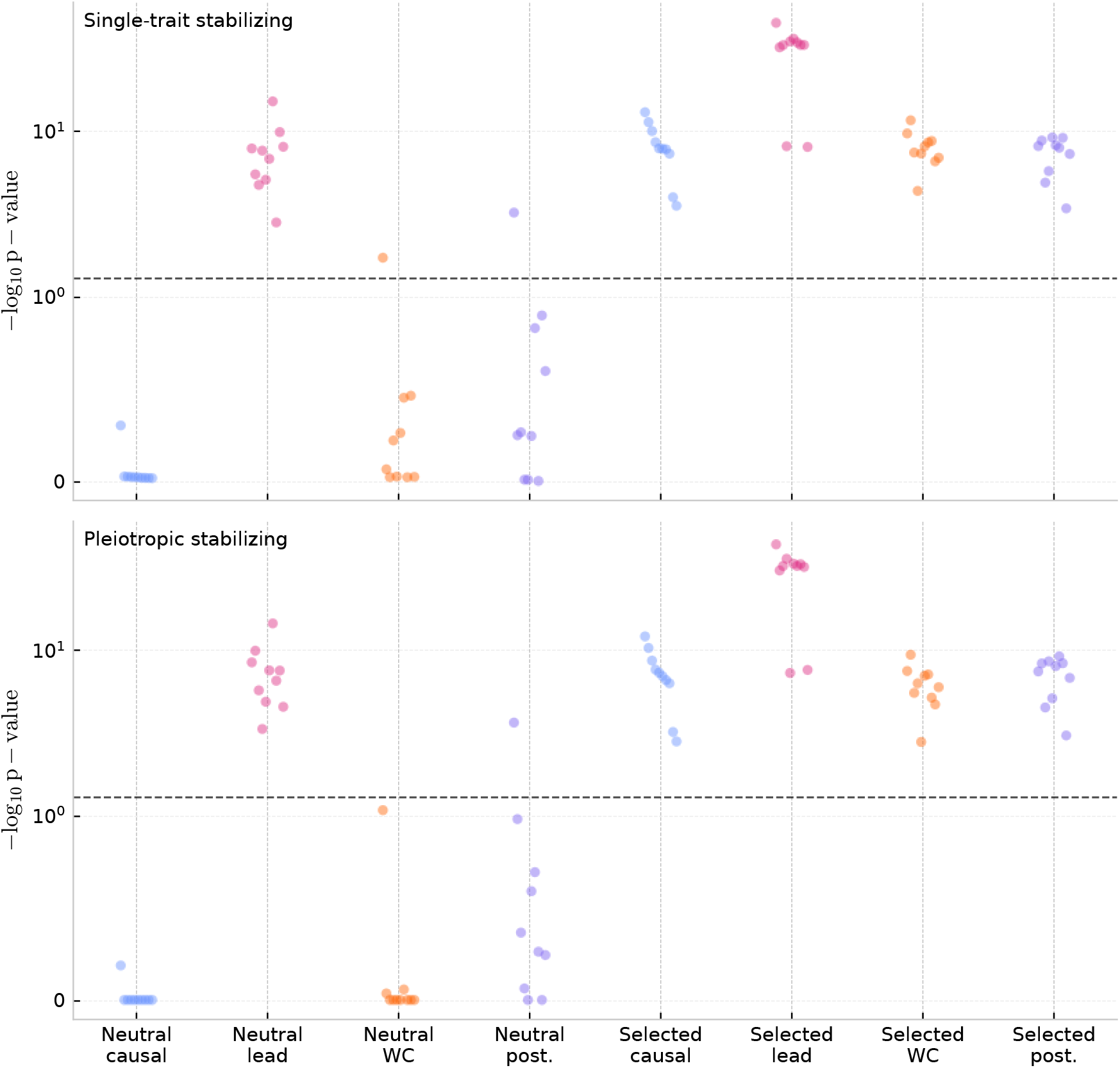
Inference of single-trait stabilizing selection on GWAS hits simulated using a linkage disequilibrium graphical model. For 11 replicate simulated datasets of the type shown in Figure S39, we plot the evidence for single-trait and pleiotropic stabilizing selection models relative to neutrality as −log_10_ *p*-value. The dashed horizontal line marks the nominal significance threshold (0.05). The use of lead variants produces spurious signals of stabilizing selection in the neutral case and inflates the evidence for selection under the alpha model used for the selection simulations. The maximum likelihood and posterior mean shrinkage adjustments reduce these effects and produces *p*-values more closely resembling what one would infer if true causative effects and frequencies were known.

**Figure S41:**
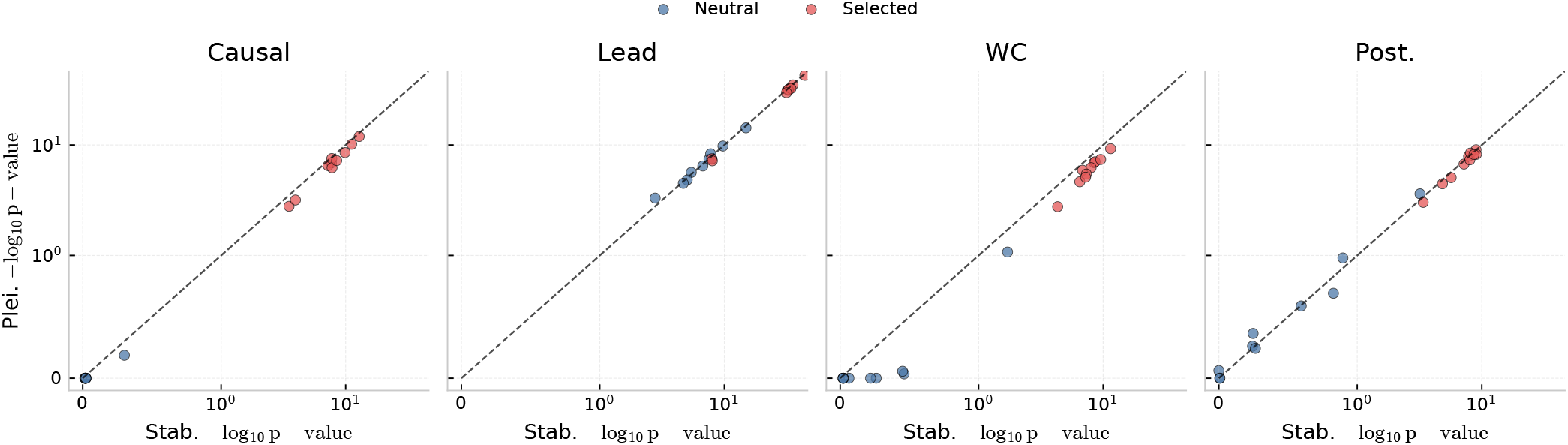
Comparing evidence for single-trait and pleiotropic stabilizing selection across simulated GraphLD replicate datasets. For the same 11 neutral and 11 selected LDL-like replicate simulations summarized in Figure S40, we compare the evidence for pleiotropic stabilizing selection against the corresponding evidence for single-trait stabilizing selection by plotting −log_10_ *p*-value for the two models against one another for causative variants, lead variants, lead variants adjusted for winner’s curse, and lead variants adjusted to posterior means. Biases associated with ascertainment and the associated corrections we implemented only systematically favor the single-trait model when using the maximum likelihood winner’s curse adjustment.

**Figure S42:**
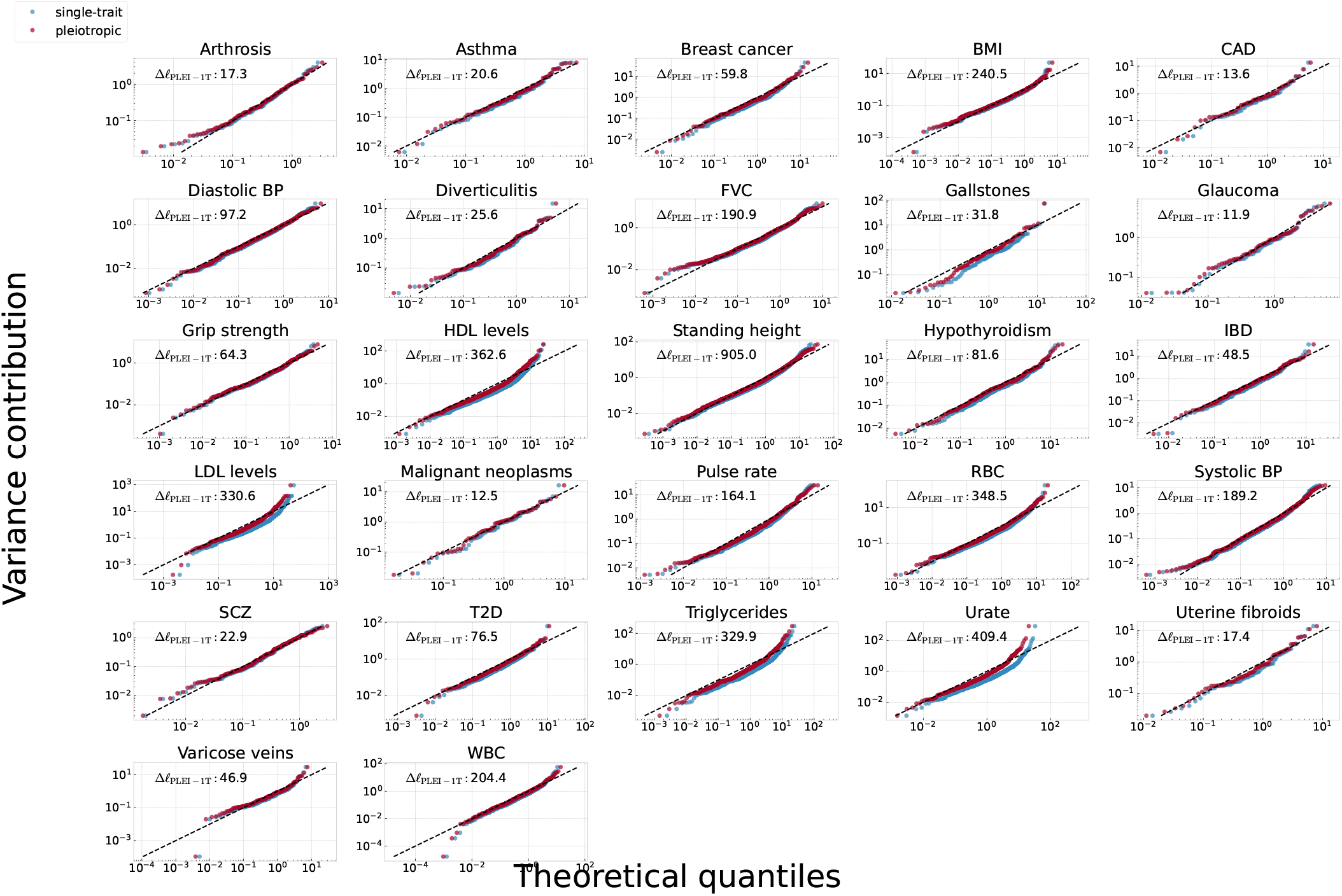
QQ plots for fits of single-trait and pleiotropic variance-contribution distributions to all trait data. The top left corner of each plot contains the excess log-likelihood for the high-dimensional variance contribution distribution. Variance contributions were transformed as (*v* − *v*^∗^)*/v*^∗^. The variance contribution threshold approximately corresponding to a *p*-value cutoff 5 × 10^−8^.

**Figure S43:**
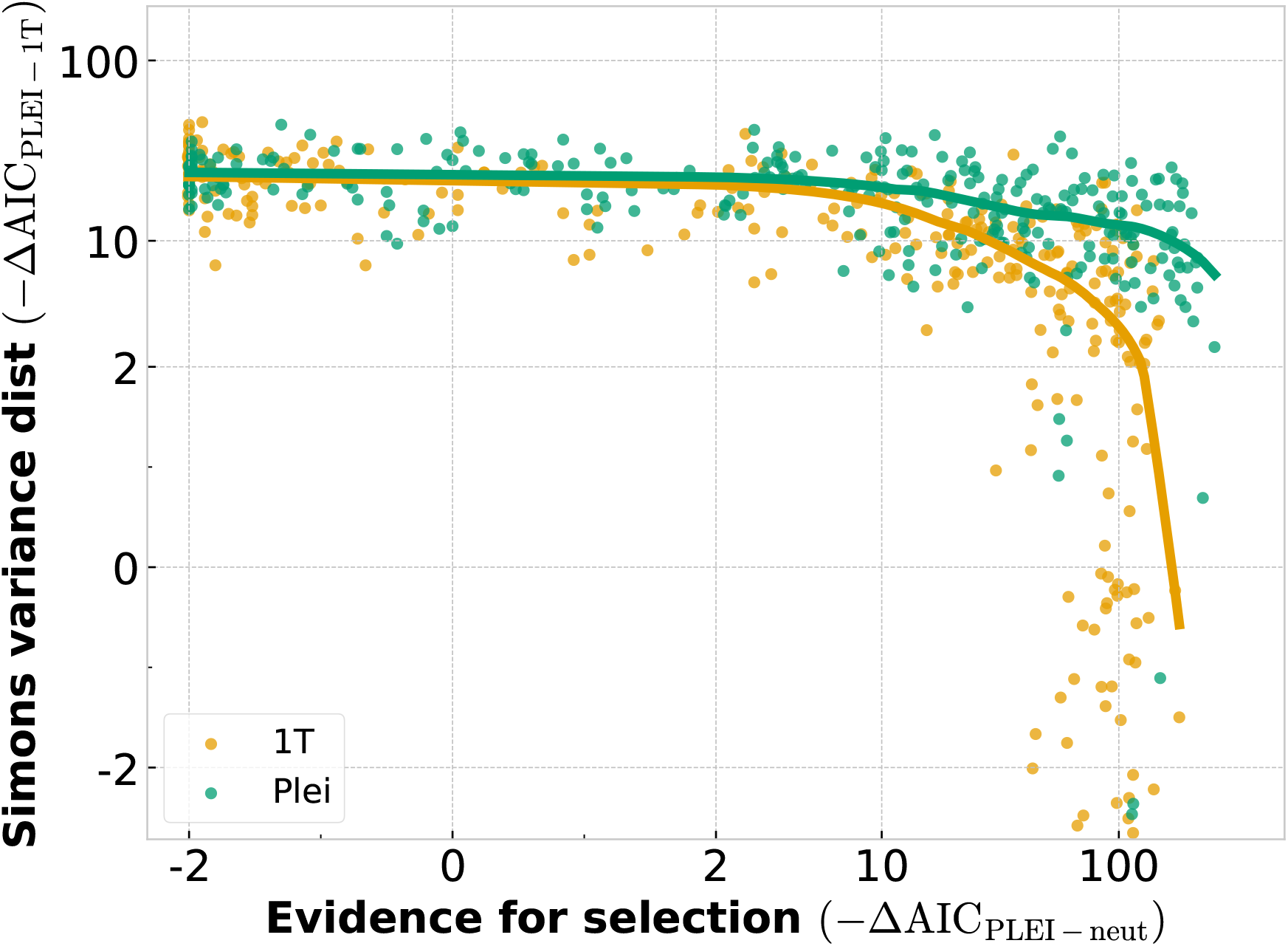
Application of variance-contribution test to trait simulations. The variance contribution test was applied to trait-based simulations of 200 GWAS hits using the estimated IBD effect size prior and a range of *I*_2_ or *I*_*p*_ values ranging from neutral to strong selection. The simulations displayed in this figure did not include ascertainment bias. The evidence for selection in each simulation was summarized using the ΔAIC value for the true simulated mode (single-trait or pleiotropic stabilizing) versus neutrality. Likelihoods for the x-axis were computed using equation (4). AIC was used instead of *p*-values in order to put both axes on the same scale. The y-axis shows the evidence for pleiotropic stabilizing selection with likelihoods computed for the tail distributions under the single-trait and pleiotropic models derived by Simons *et al*. [37]. Lines are LOESS curves showing the average behavior under each model as selection on ascertained variants becomes very strong.

**Figure S44:**
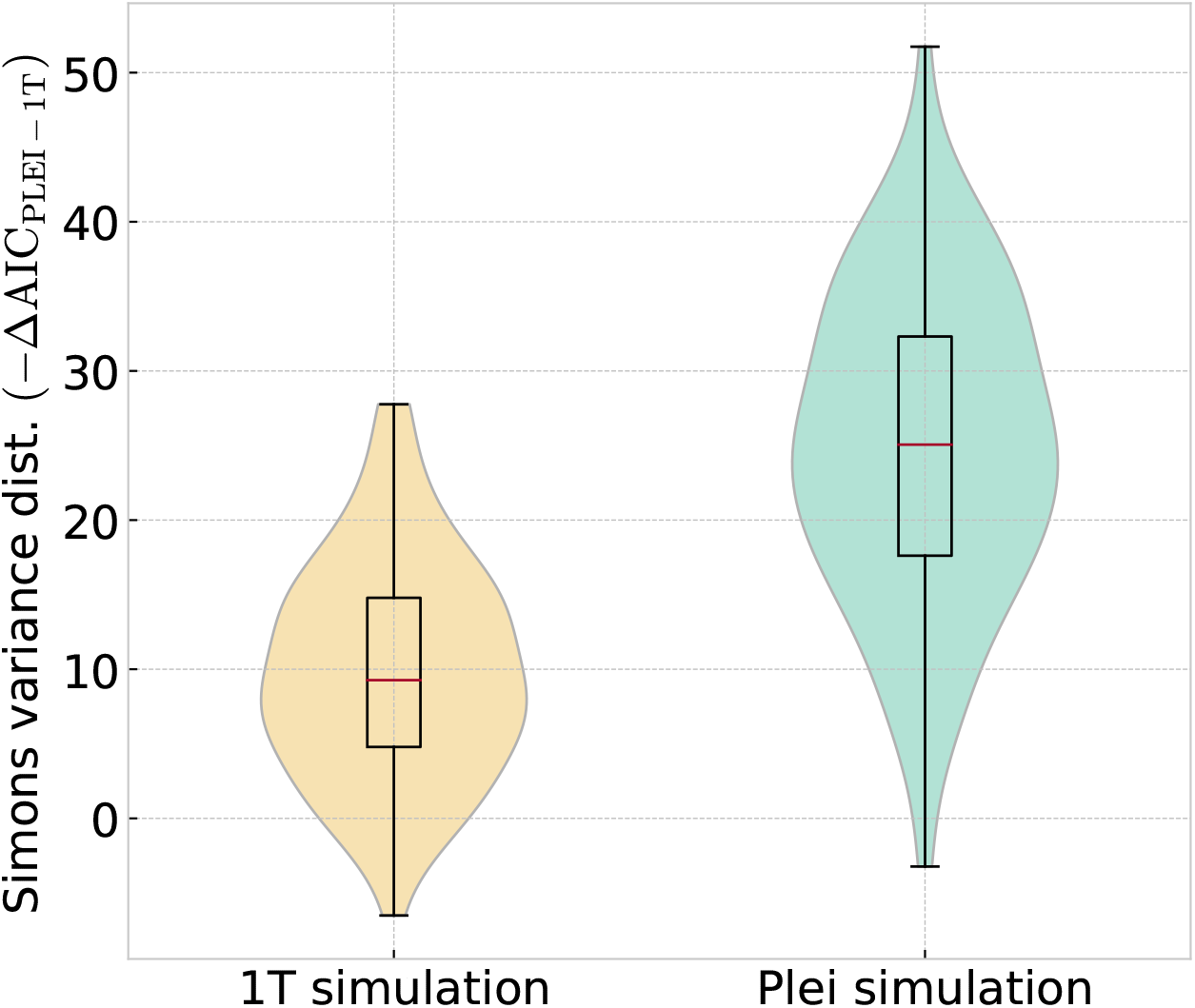
Application of variance-contribution test to DFE simulations. The variance contribution test was also applied to DFE-based simulations of 500 GWAS hits for the single-trait and pleiotropic stabilizing models. Results are displayed for a selection-scaled variance contribution cutoff of one. Boxplots summarize 100 simulations each.

**Figure S45:**
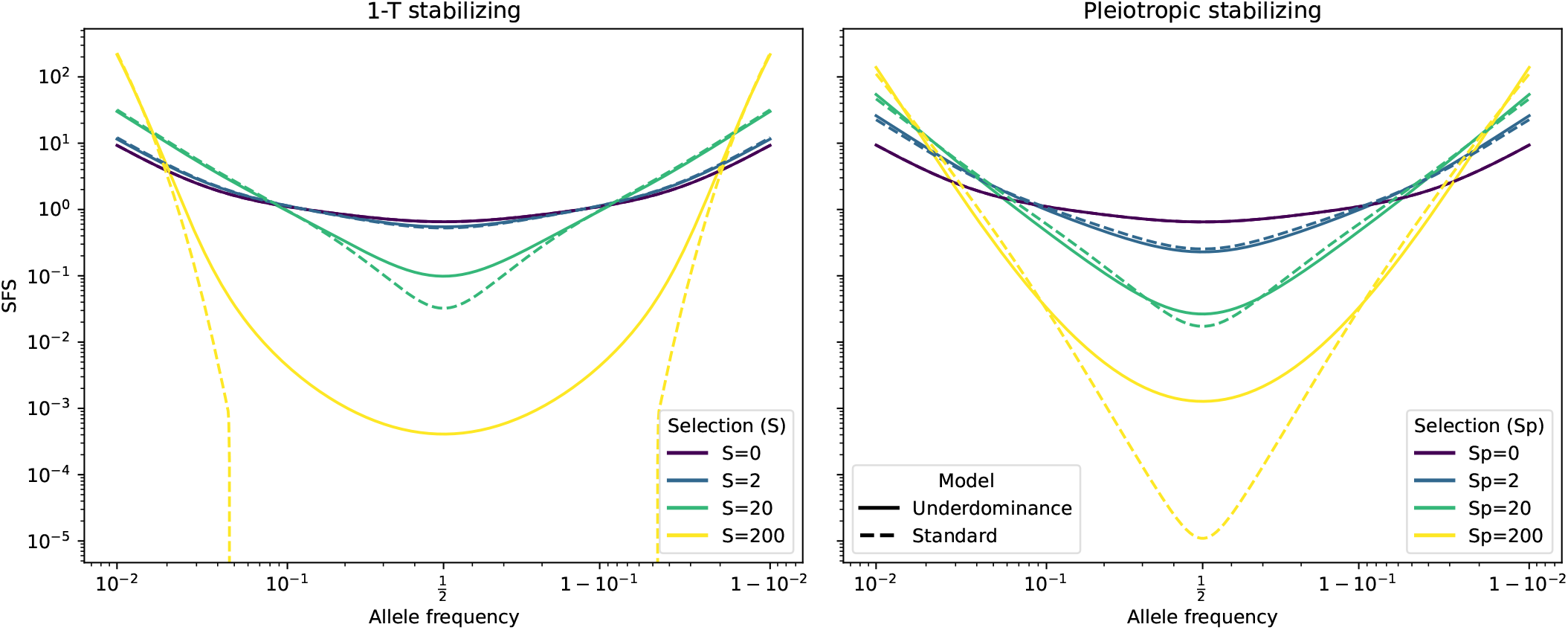
The allele frequency distribution under underdominant and additive selection. Examples of the site frequency spectrum were calculated using the equilibrium expressions for underdominant and additive selection. a) For single-trait stabilizing selection, solid lines show the underdominant model and dashed lines show the additive model for matched values of selection intensity. b) The same comparison for the pleiotropic stabilizing setting, with solid lines showing the underdominant approximation and dashed lines the additive approximation. At low frequencies the two models are similar, but the underdominant model allows more high-frequency variants.

**Figure S46:**
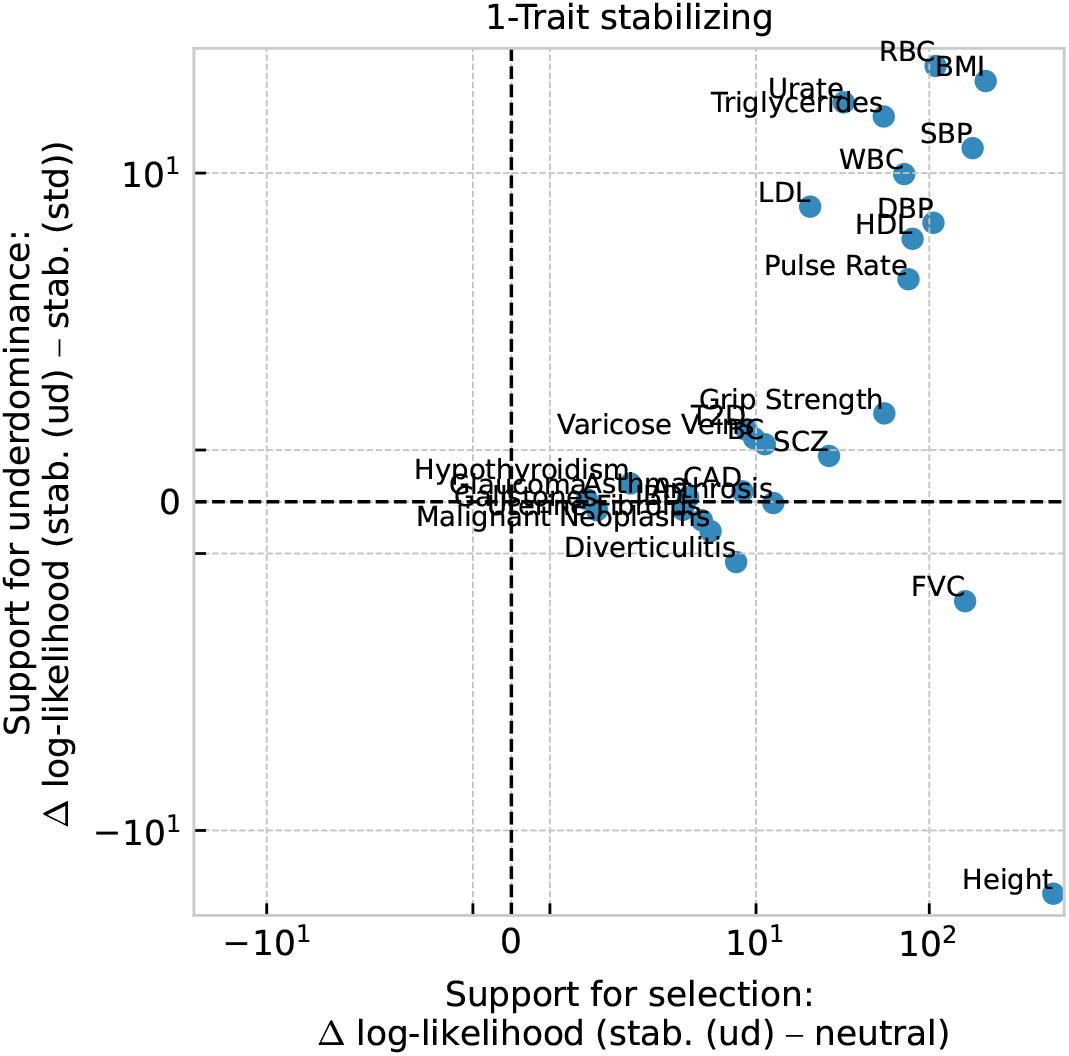
Likelihood comparison of underdominant and additive forms of single-trait stabilizing selection. For each original trait, we compare the increase in log-likelihood of the underdominant single-trait stabilizing model relative to neutrality with its increase in log-likelihood relative to the additive stabilizing approximation. Positive values on the y-axis indicate that the underdominant form provides the better fit, while positive values on the x-axis indicate support for selection relative to neutrality. Dashed lines mark zero on both axes.

**Figure S47:**
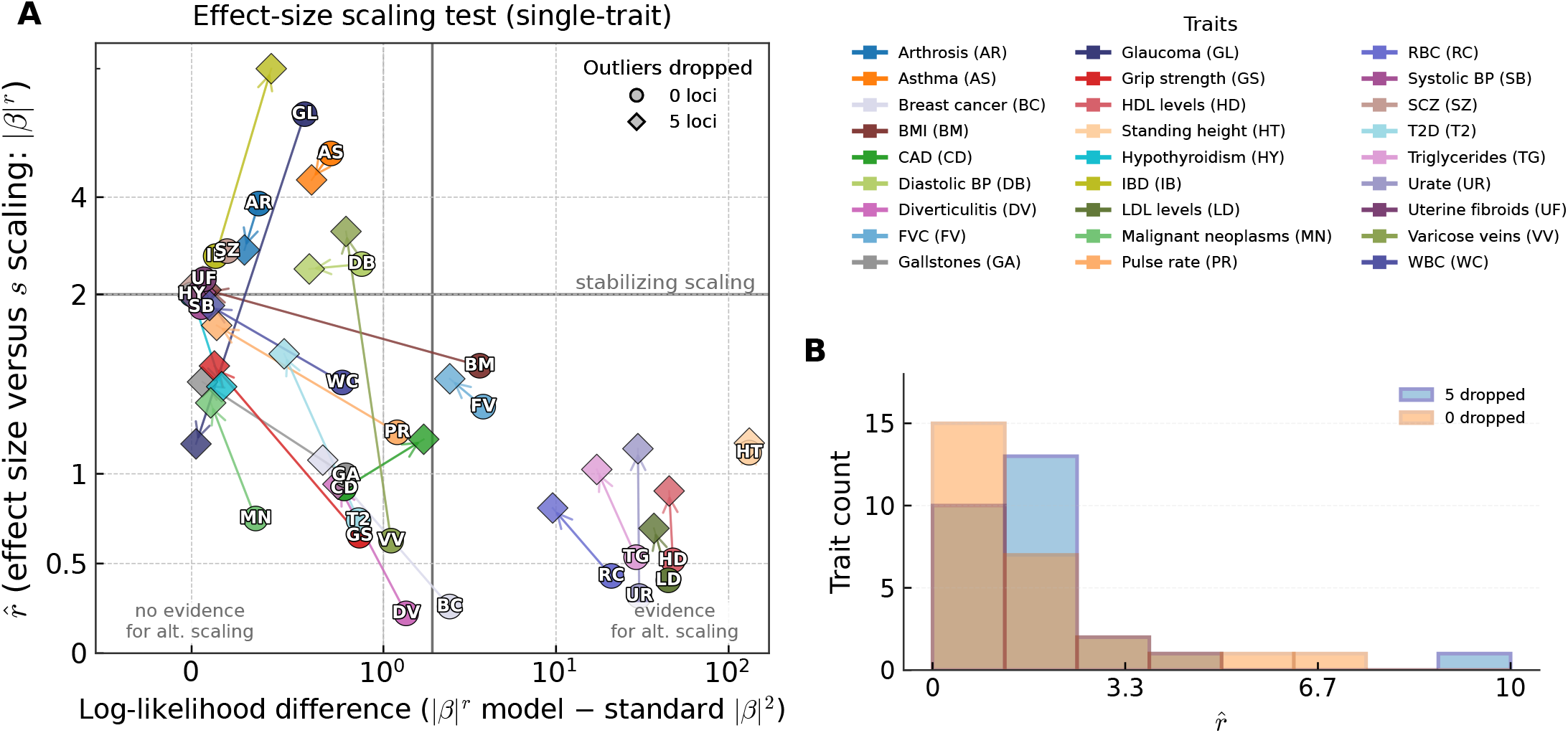
Effect-size to selection scaling under single-trait stabilizing selection with pleiotropic-model outlier removal. We fit models where the scaling exponent *r* between effect size and selection (*s* = |*β*|^*r*^) was allowed to vary from the standard stabilizing relationship (*r* = 2). Top outlier loci were identified using leave-one-out likelihood deviation under the pleiotropic stabilizing model, and traits were re-fit after removing different numbers of these loci. a) For the 27 original traits, we show the estimated 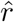 and the log-likelihood difference between the free-*r* and standard *r* = 2 single-trait stabilizing models after removing 0, 1, 2, or 5 outlier loci. The vertical reference line marks the nominal significance threshold, 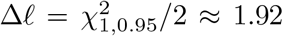, for the one-parameter extension from standard stabilizing scaling to free scaling, while the horizontal line marks the standard stabilizing expectation *r* = 2. b) The distribution of estimated scaling exponents with and without removal of the top five outlier loci.

**Figure S48:**
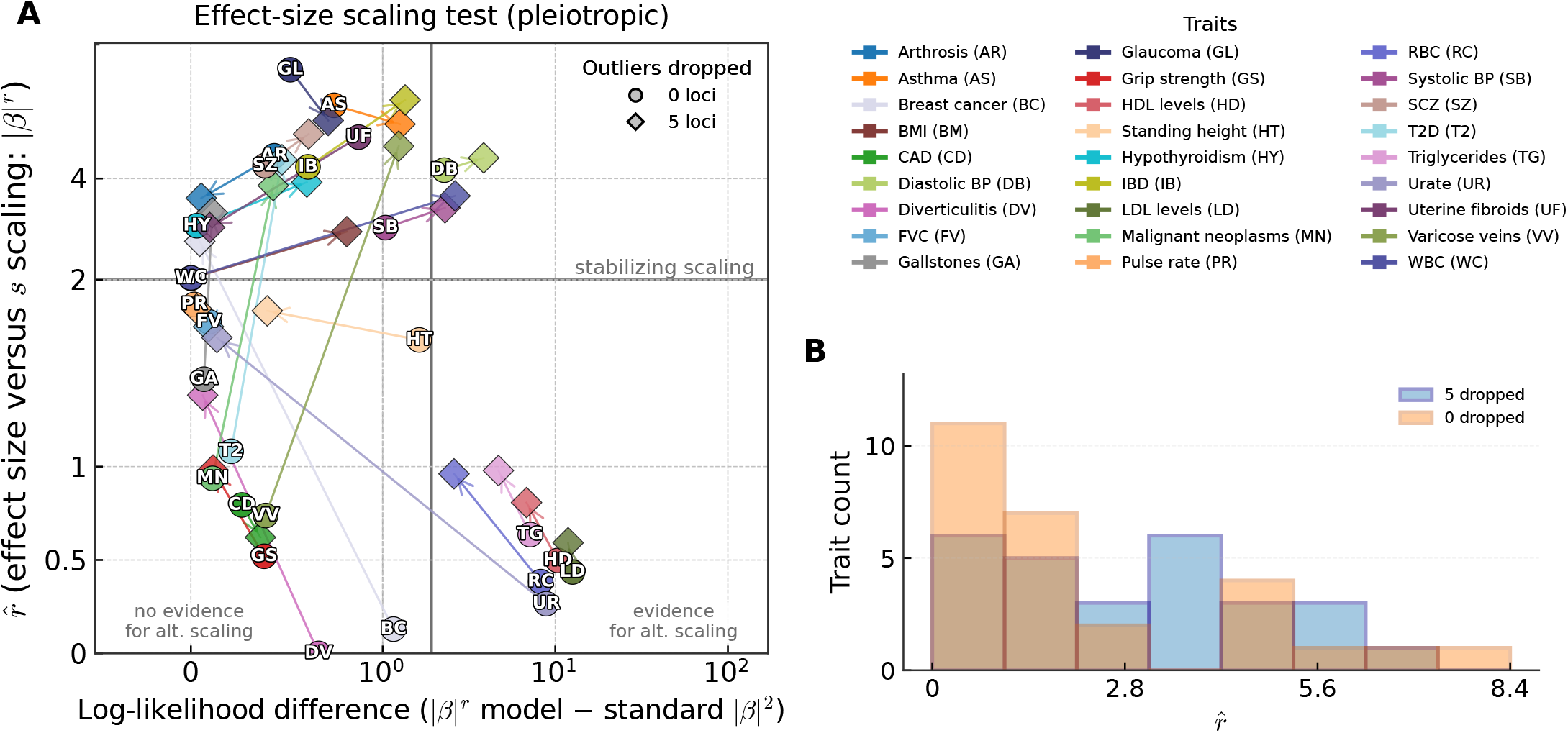
Effect-size to selection scaling under pleiotropic stabilizing selection when outliers are defined using the *r* = 0 model. We fit models where the scaling exponent *r* between effect size and selection (*s* = |*β*|^*r*^) was allowed to vary from the standard stabilizing relationship (*r* = 2). Top outlier loci were identified using leave-one-out likelihood deviation under the model with *r* = 0, and traits were re-fit after removing different numbers of these loci. The estimated *r* and the log-likelihood difference between the free-*r* and standard *r* = 2 pleiotropic stabilizing models are shown for the original trait set. The vertical reference line marks the nominal significance threshold, 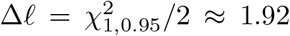, for the one-parameter extension from standard stabilizing scaling to free scaling, while the horizontal line marks the standard stabilizing expectation *r* = 2.

**Figure S49:**
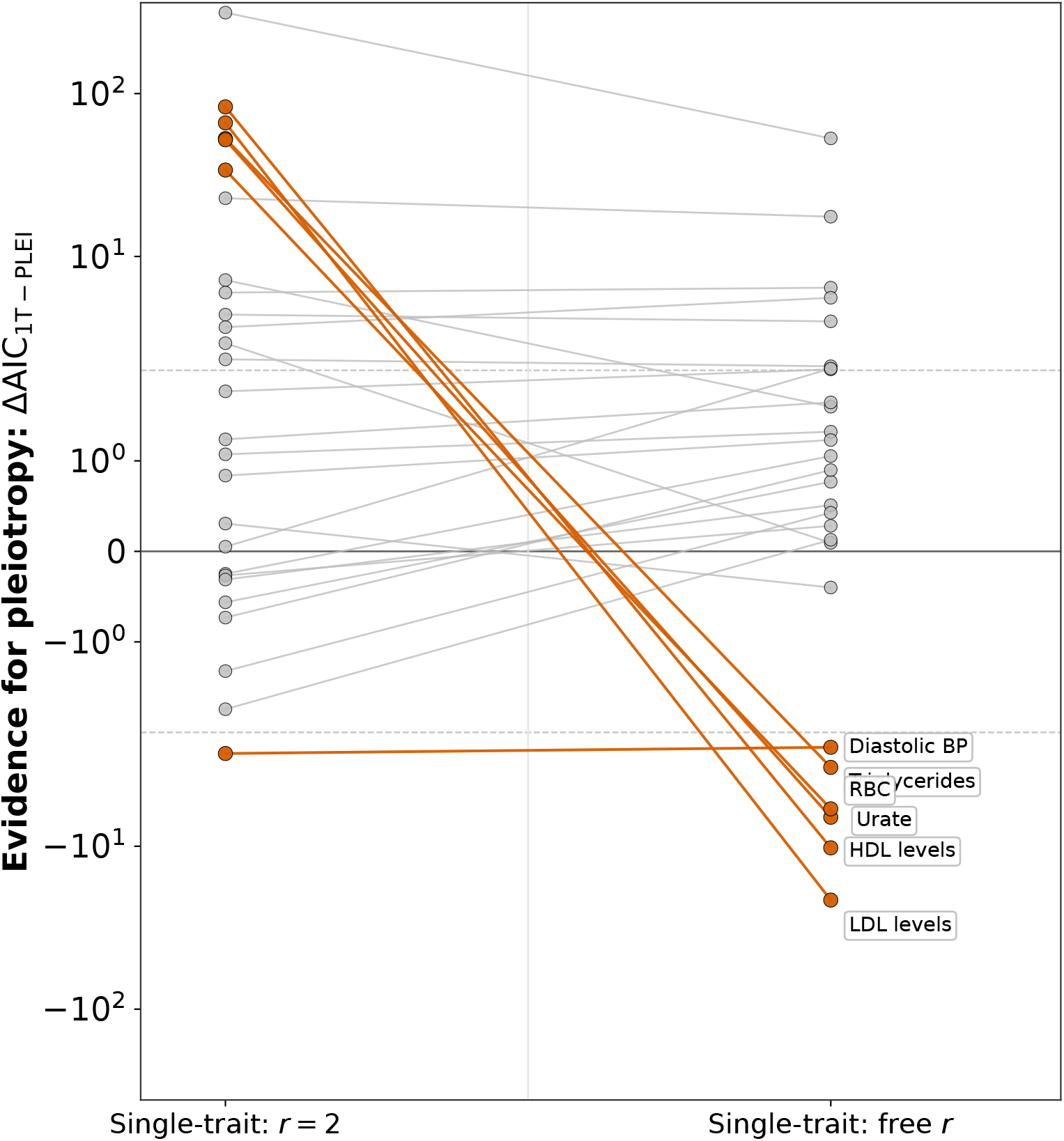
Model comparison with pleiotropic and single-trait stabilizing selection before and after allowing free effect-size scaling. We compare the AIC difference between pleiotropic and single-trait stabilizing selection across the 27 original traits before and after allowing the single-trait model to fit a free scaling exponent *r*. Positive values indicate that the pleiotropic model provides the better fit. Traits highlighted in orange are those for which the free-*r* single-trait model is favored by more than 2 AIC units.

**Figure S50:**
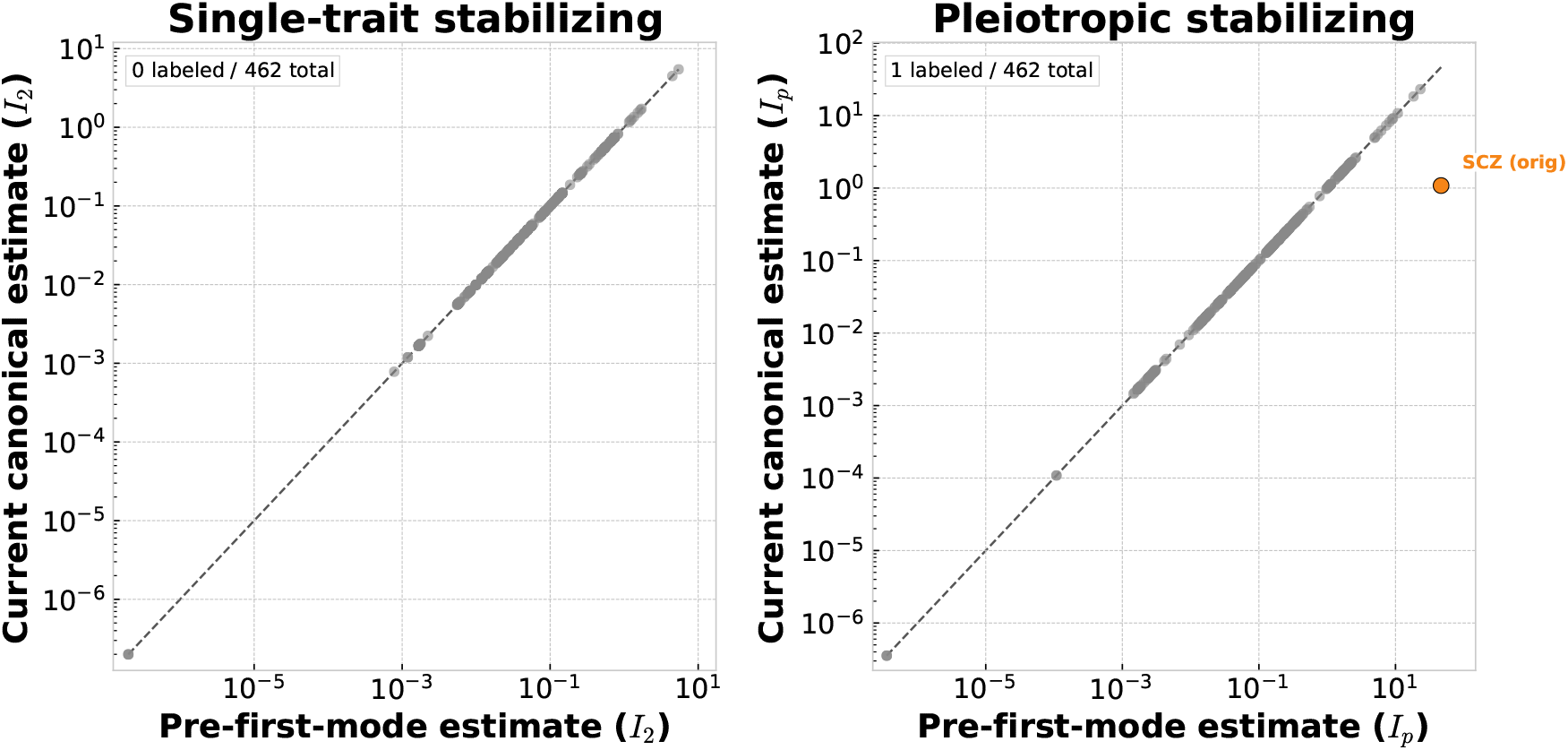
Comparison of original maximum-likelihood estimates and first-mode corrected estimates for stabilizing selection parameters. For each analysis in which first-mode correction was evaluated, we compare the original maximum-likelihood estimate to the estimate after restricting to the first mode of the likelihood surface at lower parameter values. Panel (a) shows results for the single-trait stabilizing parameter *I*_2_, while panel (b) shows results for the pleiotropic stabilizing parameter *I*_*P*_. Points falling substantially off the diagonal are labeled. In most cases the correction had little effect, indicating that the lower-parameter mode was either absent or close to the reported maximum-likelihood estimate. The only substantial change among the main reported fits was for schizophrenia under pleiotropic stabilizing selection.

